# Polygenic enrichment distinguishes disease associations of individual cells in single-cell RNA-seq data

**DOI:** 10.1101/2021.09.24.461597

**Authors:** Martin Jinye Zhang, Kangcheng Hou, Kushal K. Dey, Saori Sakaue, Karthik A. Jagadeesh, Kathryn Weinand, Aris Taychameekiatchai, Poorvi Rao, Angela Oliveira Pisco, James Zou, Bruce Wang, Michael Gandal, Soumya Raychaudhuri, Bogdan Pasaniuc, Alkes L. Price

## Abstract

Gene expression at the individual cell-level resolution, as quantified by single-cell RNA-sequencing (scRNA-seq), can provide unique insights into the pathology and cellular origin of diseases and complex traits. Here, we introduce single-cell Disease Relevance Score (`scDRS`), an approach that links scRNA-seq with polygenic risk of disease at individual cell resolution without the need for annotation of individual cells to cell types; `scDRS` identifies individual cells that show excess expression levels for genes in a disease-specific gene set constructed from GWAS data. We determined via simulations that `scDRS` is well-calibrated and powerful in identifying individual cells associated to disease. We applied `scDRS` to GWAS data from 74 diseases and complex traits (average *N* =346K) in conjunction with 16 scRNA-seq data sets spanning 1.3 million cells from 31 tissues and organs. At the cell type level, `scDRS` broadly recapitulated known links between classical cell types and disease, and also produced novel biologically plausible findings. At the individual cell level, `scDRS` identified subpopulations of disease-associated cells that are not captured by existing cell type labels, including subpopulations of CD4^+^ T cells associated with inflammatory bowel disease, partially characterized by their effector-like states; subpopulations of hippocampal CA1 pyramidal neurons associated with schizophrenia, partially characterized by their spatial location at the proximal part of the hippocampal CA1 region; and subpopulations of hepatocytes associated with triglyceride levels, partially characterized by their higher ploidy levels. At the gene level, we determined that genes whose expression across individual cells was correlated with the `scDRS` score (thus reflecting co-expression with GWAS disease genes) were strongly enriched for gold-standard drug target and Mendelian disease genes.

## Introduction

The mechanisms through which risk variants identified by genome-wide association studies (GWASs) impact critical tissues and cell types are largely unknown^1, 2^; identifying these tissues and cell types is central to our understanding of disease etiologies and will inform efforts to develop therapeutic treatments^3^. Single-cell RNA sequencing (scRNA-seq) has emerged as the tool of choice for measuring gene abundances at single-cell resolution^4, 5^, providing an increasingly clear picture of which genes are active in which cell types and also being able to identify novel cell populations within classically defined cell types. Integrating scRNA-seq with GWAS data offers the potential to identify critical tissues, cell types, and cell populations through which GWAS risk variants impact disease^6–8^, thus providing finer resolution than studies using bulk transcriptomic data^9–12^.

Previous studies integrating scRNA-seq with GWAS have largely focused on predefined cell type annotations (e.g., classical cell types defined using known marker genes), aggregating cells from the same cell type followed by evaluating overlap of the cell type-level information with GWAS^6–8^. However, this approach overlooks the considerable heterogeneity of individual cells within cell types that has been reported in studies of scRNA-seq data alone^13–18^; the underlying methods (e.g., Seurat cell-scoring function^15^, Vision^16^, and VAM^18^) have sought to explain transcriptional heterogeneity in scRNA-seq data by scoring cells based on predefined gene sets such as pathway gene sets, but do not consider polygenic disease risk from GWAS and generally do not provide individual cell-level association p-values. Integrating information from scRNA-seq data at fine-grained resolution (e.g., individual cells both within and across cell types) with polygenic signals from disease GWAS has considerable potential to produce new biological insights.

Here, we introduce *single-cell Disease Relevance Score* (`scDRS`), a method to evaluate polygenic disease enrichment of individual cells in scRNA-seq data. `scDRS` assesses whether a given cell has excess expression levels across a set of putative disease genes derived from GWAS, using an appropriately matched empirical null distribution to estimate well-calibrated p-values. To our knowledge, `scDRS` is the first method to associate individual cells in scRNA-seq data to disease GWAS. We performed extensive simulations to assess the calibration and power of `scDRS`. We then applied `scDRS` to 74 diseases and complex traits (average GWAS *N* =346K) in conjunction with 16 scRNA-seq data sets (including the Tabula Muris Senis (TMS) mouse cell atlas^19^), assessing cell type-disease associations and within-cell type association heterogeneity, including heterogeneity of T cells in their association with inflammatory bowel disease (IBD) and other autoimmune diseases, neurons in their association with schizophrenia (SCZ) and other brain-related diseases/traits, and hepatocytes in their association with triglyceride levels (TG) and other metabolic traits; we analyzed a broader set of scRNA-seq data sets to provide validation across species (human vs. mouse) and across sequencing platforms, and to include scRNA-seq data sets with experimentally determined cell types and cell states.

## Results

### Overview of methods

`scDRS` integrates gene expression profiles from scRNA-seq with polygenic disease information from GWAS to associate individual cells to disease without the need for annotation of individual cells to cell types, by assessing the excess expression of putative disease genes from GWAS in a given cell relative to other genes with similar expression levels across all cells. `scDRS` consists of three steps (Fig. 1, Methods, and Supp. Note). First, `scDRS` constructs a set of putative disease genes from GWAS summary statistics using MAGMA^20^, an existing gene scoring method (top 1,000 MAGMA genes; see Methods for other choices evaluated). Second, `scDRS` quantifies the aggregate expression of the putative disease genes in each cell to generate cell-specific *raw disease scores*; to maximize power, each putative disease gene is weighted by its GWAS MAGMA z-score and inversely weighted by its gene-specific technical noise level in the single-cell data, estimated via modeling the mean-variance relationship across genes^18, 21^ (alternative choices of cell scores are evaluated in Methods). To determine statistical significance, `scDRS` also generates 1,000 sets of cell-specific *raw control scores* at Monte Carlo (MC) samples of matched control gene sets (matching the gene set size, mean expression, and expression variance of the putative disease genes); cell-specific MC p-values are defined as the proportion of the 1,000 raw control scores for a given cell exceeding the raw disease score for that cell^22^. Third, `scDRS` approximates the ideal MC p-values (obtained using ≫ 1,000 MC samples) by pooling control scores across cells. Specifically, it normalizes the raw disease score and raw control scores for each cell (producing the *normalized disease score* and *normalized control scores*), and then computes cell-level p-values based on the empirical distribution of the pooled normalized control scores across all control gene sets and all cells; this approximation relies on the assumption that the raw control score distributions (across the 1,000 control gene sets, for each cell) are from the same parametric distribution (e.g., normal distributions with different parameters), a reasonable assumption when the disease gene set is neither too small nor too large (e.g., *>*50 genes and *<*20% of all genes; Methods). Importantly, `scDRS` does not use cell type or other cell-level annotations, although these annotations can be of value when interpreting its results. `scDRS` is computationally efficient and scales linearly with the number of cells and number of control gene sets for both running time and memory (Methods).

**Figure 1.**
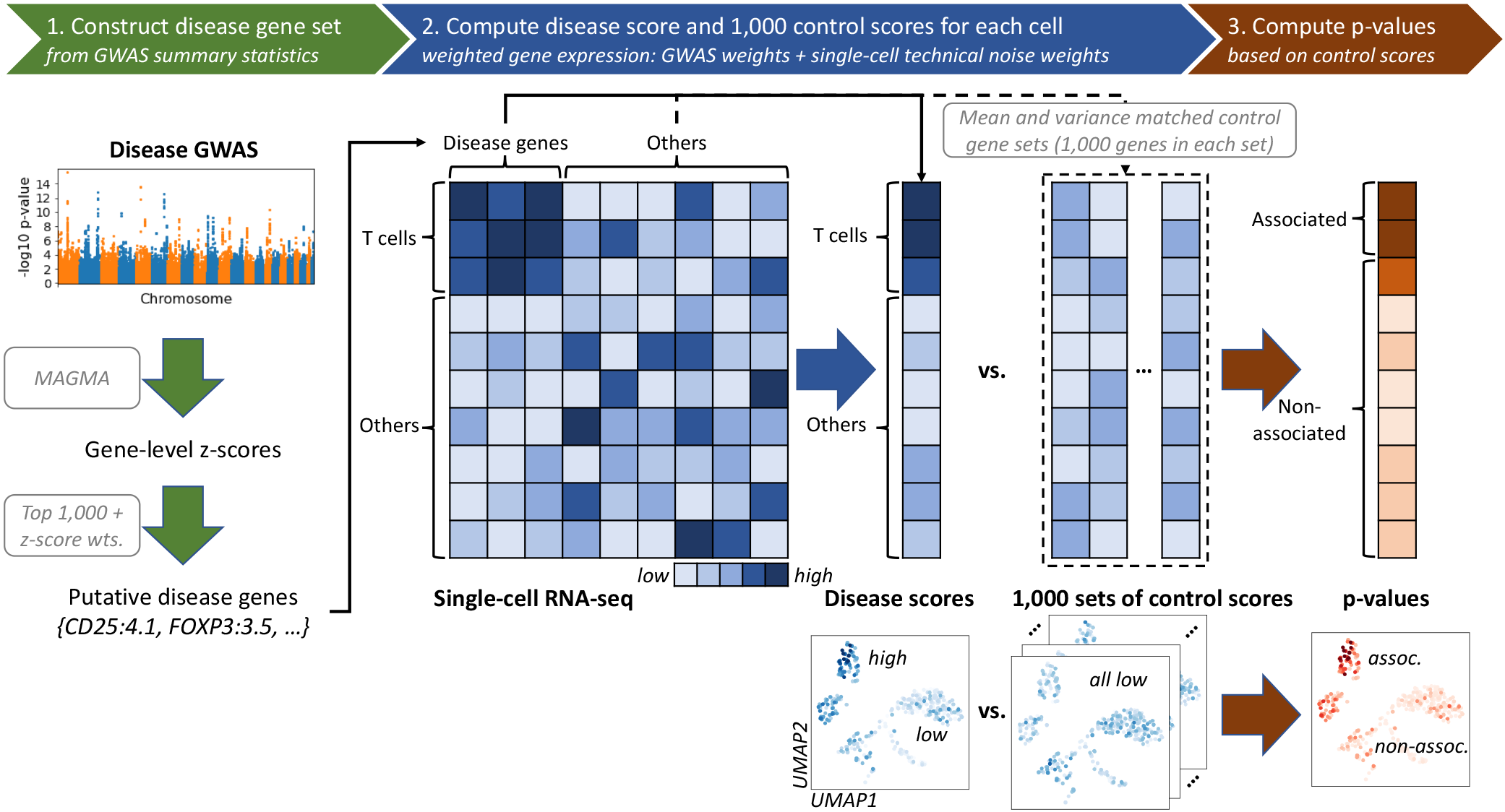
Overview of scDRS method. `scDRS` takes a disease GWAS and an scRNA-seq data set as input and outputs individual cell-level p-values for association with the disease. **(1)** `scDRS` constructs a set of putative disease genes from GWAS summary statistics by selecting the top 1,000 MAGMA genes; these putative disease genes are expected to have higher expression levels in the relevant cell population. **(2)** `scDRS` computes a raw disease score for each cell, quantifying the aggregate expression of the putative disease genes in that cell; to maximize power, each putative disease gene is weighted by its GWAS MAGMA z-score and inversely weighted by its gene-specific technical noise level in scRNA-seq. `scDRS` also computes a set of 1,000 Monte Carlo raw control scores for each cell, in each case using a random set of control genes matching the gene set size, mean expression, and expression variance of the putative disease genes. **(3)** `scDRS` normalizes the raw disease score and raw control scores across gene sets and across cells, and then computes a p-value for each cell based on the empirical distribution of the pooled normalized control scores across all control gene sets and all cells. The choice of 1,000 for the number of putative disease genes and the choice of 1,000 for the number of control scores are independent.

`scDRS` outputs individual cell-level p-values (testing for cell-disease associations as described above), normalized disease scores, and 1,000 sets of normalized control scores (referred to as “disease scores” and “control scores” in the rest of the paper) that can be used for a wide range of downstream applications (Methods). Here, we focus on three downstream analyses. First, we perform *cell type-level* analyses to associate predefined cell types to disease and assess heterogeneity in association to disease across cells within a predefined cell type. Second, we perform *individual cell-level* analyses to associate individual cells to disease and correlate individual cell-level variables to the `scDRS` disease score. Third, we perform *gene-level* analyses to prioritize disease-relevant genes whose expression is correlated with the `scDRS` disease score, reflecting co-expression with genes implicated by disease GWAS.

We analyzed publicly available GWAS summary statistics of 74 diseases and complex traits (average *N*=346K; Supp. Table 1) in conjunction with 16 scRNA-seq or single-nucleus RNA-seq (snRNA-seq) data sets spanning 1.3 million cells from 31 tissues and organs from mouse (*mus musculus*) and human (*homo sapiens*) (Supp. Table 2; 15 out of 16 data sets publicly available; Data Availability). The single-cell data sets include two mouse cell atlases from the Tabula Muris Senis (TMS)^19^ collected using different technologies (fluorescence-activated cell sorting followed by Smart-seq2 amplification^23^ for the TMS FACS data and 10x microfluidic droplet capture and amplification^24^ for the TMS droplet data), the unpublished Tabula Sapiens (TS) human cell atlas^25^, and other data sets focusing on specific tissues containing well-annotated cell types and cell states. We focused on the TMS FACS data in our primary analyses due to its comprehensive coverage of 23 tissues and 120 cell types and more accurate quantification of gene expression levels (via Smart-seq2); we used the other 15 data sets to validate our results. We note the extensive use of mouse gene expression data to study human diseases and complex traits (see Bryois et al.^8^, other studies^6, 7, 9, 12, 26^, and Discussion).

### Simulations assessing calibration and power

We performed null simulations and causal simulations to assess the calibration and power of `scDRS`, comparing `scDRS` to three state-of-art methods for scoring individual cells with respect to a specific gene set: Seurat (cell-scoring function)^15^, Vision^16^, and VAM^18^. To our knowledge, VAM is the only method for scoring individual cells that provides cell-level association p-values; Seurat and Vision provide quantitative cell-level scores that we transformed to p-values based on the standard normal distribution (Methods).

First, we evaluated each method in null simulations in which no cells have systematically higher expression across the putative disease genes analyzed. We subsampled 10,000 cells from the TMS FACS data and randomly selected 1,000 putative disease genes. We simulated GWAS gene weights for `scDRS` matching the MAGMA z-score distributions in real traits and used binary disease gene sets for the other 3 methods. `scDRS` and Seurat produced well-calibrated p-values, Vision suffered slightly inflated type I error, and VAM suffered severely inflated type I error (Fig. 2A and Supp. Table 11). The slight miscalibration of Vision may be due to the mismatch between the normal distribution used for computing p-values and the actual null distribution of the cell-level scores. The poor calibration of VAM may be because it uses a permutation-based test that assumes independence between genes under the null, an assumption that is likely to be violated in scRNA-seq data. Secondary analyses are reported in the Supp. Note, including null simulations with other numbers of putative disease genes or biased sets of putative disease genes (e.g., randomly selected from genes with high mean expression) (Supp. Fig. 4,5), and null simulations for `scDRS` cell type-level association analysis (Supp. Table 12).

**Figure 2.**
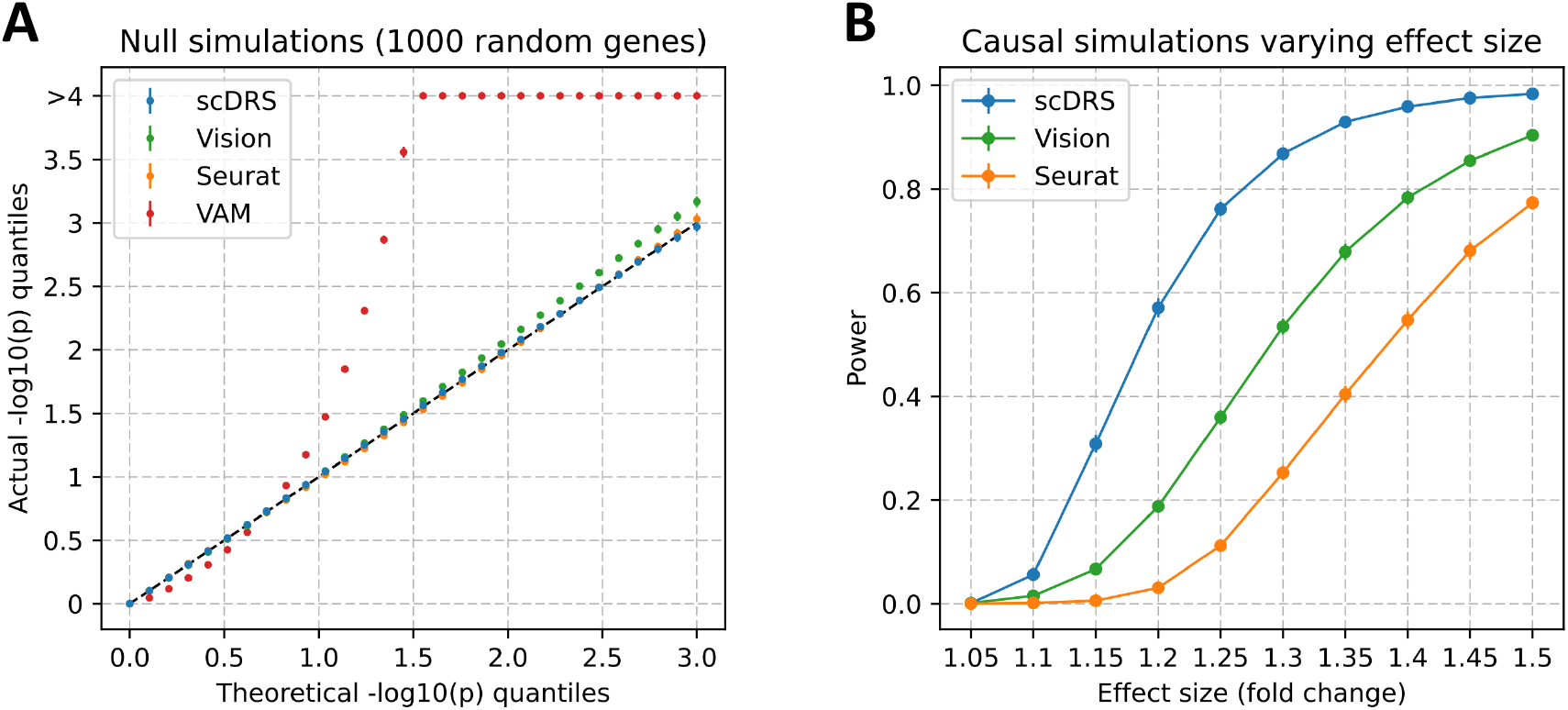
Results for null and causal simulations. **(A)** Q-Q plot for null simulations using 1,000 randomly selected genes as the putative disease genes. Random GWAS gene weights were used for `scDRS` matching the MAGMA z-score distributions in real traits while binary gene sets were used for the other 3 methods. The x-axis denotes theoretical −log_10_ p-value quantiles and the y-axis denotes actual −log_10_ p-value quantiles for different methods. Each point is based on 100 simulation replicates (with 10,000 cells per simulation replicate); error bars denote 95% confidence intervals (all error bars are *<*0.05 from the point estimate). Numerical results are reported in Supp. Table 11 and additional results are reported in Supp. Fig. 4. **(B)** Power for casual simulations with perturbed expression of causal genes in causal cells. We report the power at FDR=0.1 for different methods and different effect sizes. Each point is based on 100 simulation replicates (with 10,000 cells per simulation replicate); error bars denote 95% confidence intervals (all error bars are *<*0.02 from the point estimate). Numerical results are reported in Supp. Table 13 and additional results are reported in Supp. Fig. 6.

Next, we evaluated `scDRS`, Seurat and Vision in causal simulations in which a subset of causal cells has systematically higher expression across putative disease genes (we did not include VAM, which was not well-calibrated in null simulations). We used the same 10,000 cells subsampled from the TMS FACS data, randomly selected 1,000 causal disease genes, randomly selected 500 of the 10,000 cells as causal cells and artificially perturbed their expression levels to be higher (1.05-1.50 times for different simulations) across the 1,000 causal disease genes, and randomly selected 1,000 putative disease genes (provided as input to each method) with 25% overlap with the 1,000 causal disease genes. We used the binary gene set for all 3 methods because there were no GWAS weights involved in generating the data. We determined that `scDRS` attained higher power than Seurat and Vision to detect individual cell-disease associations at FDR*<*0.1 (Fig. 2B and Supp. Table 13); the improved power of `scDRS` may be due to its incorporation of gene-specific weights that downweight genes with higher levels of technical noise. Please see secondary analyses in Supp. Note, including simulations with other levels of overlap between the 1,000 causal genes and 1,000 putative disease genes (Supp. Fig. 6).

### Results across 120 TMS cell types for 74 diseases and complex traits

We analyzed GWAS data from 74 diseases and complex traits (average *N*=346K; Supp. Table 1,8) in conjunction with the TMS FACS data with 120 cell types (cells from different tissues were combined for a given cell type; Supp. Table 5). We first report `scDRS` cell type-level results, aggregated for each cell type from the `scDRS` individual cell-level results; the individual cell-level results are discussed in subsequent sections. Results for a representative subset of 19 cell types and 22 diseases/traits are reported in Fig. 3 (complete results in Supp. Fig. 7 and Supp. Table 14). Within this subset, `scDRS` identified 80 associated cell type-disease pairs (FDR*<*0.05; squares in Fig. 3) and detected significant within-cell type disease-association heterogeneity for 43 of these 80 associated cell type-disease pairs (FDR*<*0.05; cross symbols in Fig. 3; 273 of 597 across all pairs of the 120 cell types and 74 diseases/traits). We also report the proportion of significantly associated individual cells for each cell type-disease pair (FDR*<*0.1, a less stringent threshold as false positive associations of individual cells are less problematic and we do not focus on the results for any one specific cell; heatmap colors in Fig. 3). We note these associated cell type-disease pairs (and individual cell-disease associations discussed in subsequent sections) may reflect indirect tagging of causal cell types rather than direct causal associations, analogous to previous works (see Discussion).

**Figure 3.**
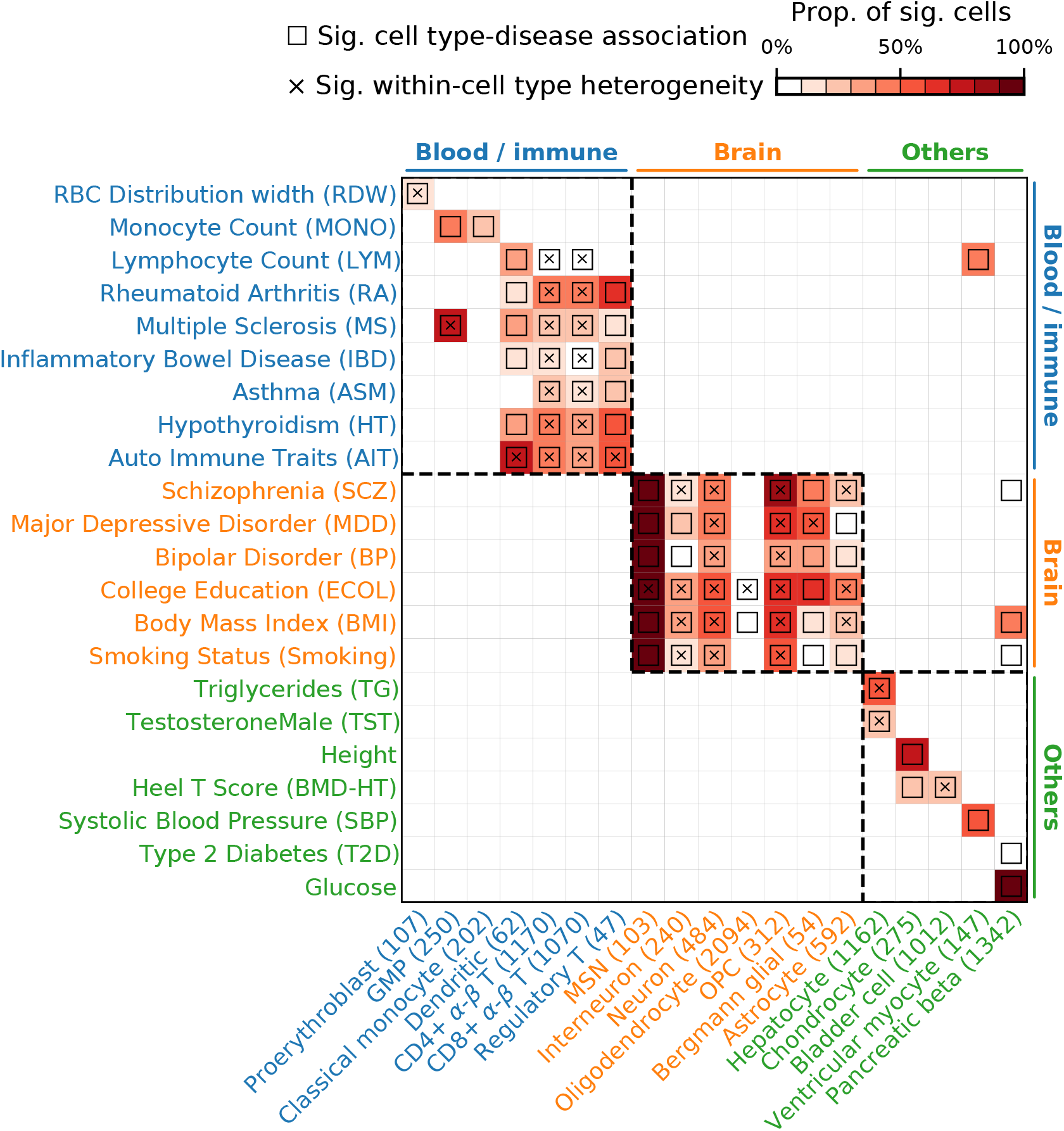
Disease associations at the cell type-level. We report `scDRS` results for individual cells aggregated at the cell type-level for a subset of 19 cell types and 22 diseases/traits in the TMS FACS data. Each row represents a disease/trait and each column represents a cell type (with number of cells indicated in parentheses). Heatmap colors for each cell type-disease pair denote the proportion of significantly associated cells (FDR*<*0.1 across all cells for a given disease). Squares denote significant cell type-disease associations (FDR*<*0.05 across all pairs of the 120 cell types and 74 diseases/traits; p-values via MC test; Methods). Cross symbols denote significant heterogeneity in association with disease across individual cells within a given cell type (FDR*<*0.05 across all pairs; p-values via MC test; Methods). Heatmap colors (*>*10% of cells associated) and cross symbols are omitted for cell type-disease pairs with non-significant cell type-disease associations via MC test (heatmap colors omitted for 1 pair (Dendritic-ASM) and cross symbols omitted for 6 pairs (CD4^+^ *α*-*β* T-MONO, CD8^+^ *α*-*β* T-MONO, bladder cell-RA, bladder cell-ASM, oligodendrocyte-BP, and dendritic-BMD-HT)). Auto Immune Traits (AIT) represents a collection of diseases in the UK Biobank that characterize autoimmune physiopathogenic etiology^124, 125^. Abbreviated cell type names include red blood cell (RBC), granulocyte monocyte progenitor (GMP), medium spiny neuron (MSN), and oligodendrocyte precursor cell (OPC). Neuron refers to neuronal cells with undetermined subtypes (whereas MSN and interneuron (non-overlapping with neuron) refer to neuronal cells with those inferred subtypes). Complete results for 120 cell types and 74 diseases/traits are reported in Supp. Fig. 7 and Supp. Table 14.

For cell type-disease associations, as expected, `scDRS` broadly associated blood/immune cell types with blood/immune-related diseases/traits, brain cell types with brain-related diseases/traits, and other cell types with other diseases/traits (block-diagonal pattern in Fig. 3; exceptions are discussed in Supp. Note).

We discuss 3 main findings for the blood/immune-related diseases/traits (upper left block in Fig. 3). First, different blood/immune cell types were associated with the corresponding blood cell traits, including proerythroblasts with RDW, classical monocytes with monocyte count, and adaptive immune cells with lymphocyte count. We detected significant heterogeneity across cells for the proerythroblast-RDW association, which may correspond to erythrocytes at different differentiation stages^27^ (see Supp. Fig. 8). Second, immune cell types were associated with immune diseases, including dendritic cells, CD4^+^ *α/β* T cells, CD8^+^ *α/β* T cells, and/or regulatory T cells with rheumatoid arthritis (RA), multiple sclerosis (MS), and IBD, consistent with previous findings^12, 28^. We detected significant heterogeneity across cells for many of these cell type-disease associations, consistent with the known diversity within the T cell population (see the T cell subsection below). Third, granulocyte monocyte progenitors (GMP) were strongly associated with MS, highlighting the role of myeloid cells in MS^29, 30^.

We discuss 2 main findings for brain-related diseases/traits (middle block in Fig. 3). First, neuronal cell types, including medium spiny neurons (MSNs), interneurons, and neurons (neuronal cells with undetermined subtypes), were associated with schizophrenia (SCZ), major depressive disorder (MDD), bipolar disorder (BP), college education (ECOL), and several other brain-related traits; the role of MSN in SCZ, MDD, BP, and ECOL is supported by previous genetic studies^8, 26, 31, 32^. We detected significant heterogeneity across neurons in their association with most brain-related diseases/traits (see the neuron subsection below). Second, oligodendrocytes, oligodendrocyte precursor cells (OPCs) were also associated with multiple brain-related diseases/traits. These associations are less clear in existing genetic studies^6, 8, 26, 33^, but are biologically plausible, consistent with the increasingly discussed role of oligodendrocyte lineage cells in brain diseases/traits: the differentiation and myelination of oligodendrocyte lineage cells are important to maintain the functionality of neuronal cells^34, 35^. We detected significant heterogeneity across OPCs in their association with many brain-related diseases/traits, consistent with recent evidence of functionally diverse states of OPCs^36^, traditionally considered to be a homogeneous population (see Supp. Fig. 9).

We discuss 2 main findings for other diseases/traits (lower right block in Fig. 3). First, hepatocytes were associated with several metabolic traits including TG and testosterone (TST) (and other lipid traits; Supp. Fig. 7); hepatocytes are known to play an important role in metabolism^37^. We detected significant heterogeneity across hepatocytes in their association with TG and TST (see the hepatocyte subsection below). Second, other cell types, including chondrocytes, bladder cells, ventricular myocytes and pancreatic beta cells, were associated with their corresponding expected diseases/traits, consistent with previous genetic studies^38–41^.

We performed 4 secondary analyses to assess robustness of these results; further details are provided in the Supp. Note. First, we determined that `scDRS` cell type-disease associations are highly consistent between data sets collected using different technologies (TMS FACS vs. TMS droplet) and reasonably consistent between mouse and human data (TMS FACS vs. TS FACS) (Supp. Fig. 10). Second, we determined that cell type-disease associations are highly consistent between `scDRS` and 4 existing cell type-level association methods (LDSC-SEG^12^ and 3 methods in Bryois et al.^8^; Supp. Fig. 11). Third, since the `scDRS` results may be biased towards major cell types with many cells, we implemented a version of `scDRS` that adjusts for cell type proportions, and determined that it was highly consistent with the default version (median of 0.97 across 74 diseases for the `scDRS` disease score correlation computed across all TMS FACS cells) and well-calibrated in null simulations (Supp. Fig. 4; Methods). Fourth, we determined that `scDRS` is robust to different scaling factors for size-factor normalization (median of 0.90 across 74 diseases for the `scDRS` disease score correlation between scaling to the default 10,000 vs. 1 million reads per cell computed across all TMS FACS cells; Methods).

We performed 2 secondary analyses to assess alternative versions of `scDRS`; further details are provided in the Supp. Note. First, we determined that the default version of `scDRS` outperformed alternative versions using different disease gene selection methods (top 100, top 500, top 2,000, FWER*<*5%, FDR*<*1%, instead of top 1,000), weighting methods for the selected disease genes (no weights, GWAS MAGMA z-score weights, single-cell technical noise weights, instead of using both sets of weights), or MAGMA gene window sizes (0 kb, 50 kb, instead of 10 kb) (Supp. Fig. 12,13, Supp. Table 17,18; Methods). Second, we determined that the default weighted score (only capturing overexpression of putative disease genes in the relevant cell population) substantially outperformed an overdispersion score capturing both overexpression and underexpression (Supp. Fig. 14; Methods).

### Heterogeneous subpopulations of T cells associated with autoimmune disease

We sought to further understand the heterogeneity across T cells in the TMS FACS data in their association with autoimmune diseases (Fig. 3). We jointly analyzed all T cells in the TMS FACS data (3,769 cells, spanning 15 tissues). Since the original study clustered cells from different tissues separately^19^, we reclustered these T cells, resulting in 11 clusters (Fig. 4A; Methods); we verified that batch effects were not observed for tissue, age, or sex (Supp. Fig. 15). We considered 10 autoimmune diseases: IBD, Crohn’s disease (CD), ulcerative colitis (UC), RA, MS, AIT, hypothyroidism (HT), eczema, asthma (ASM), and respiratory and ear-nose-throat diseases (RR-ENT) (Supp. Table 1); we also considered height as a negative control trait.

**Figure 4.**
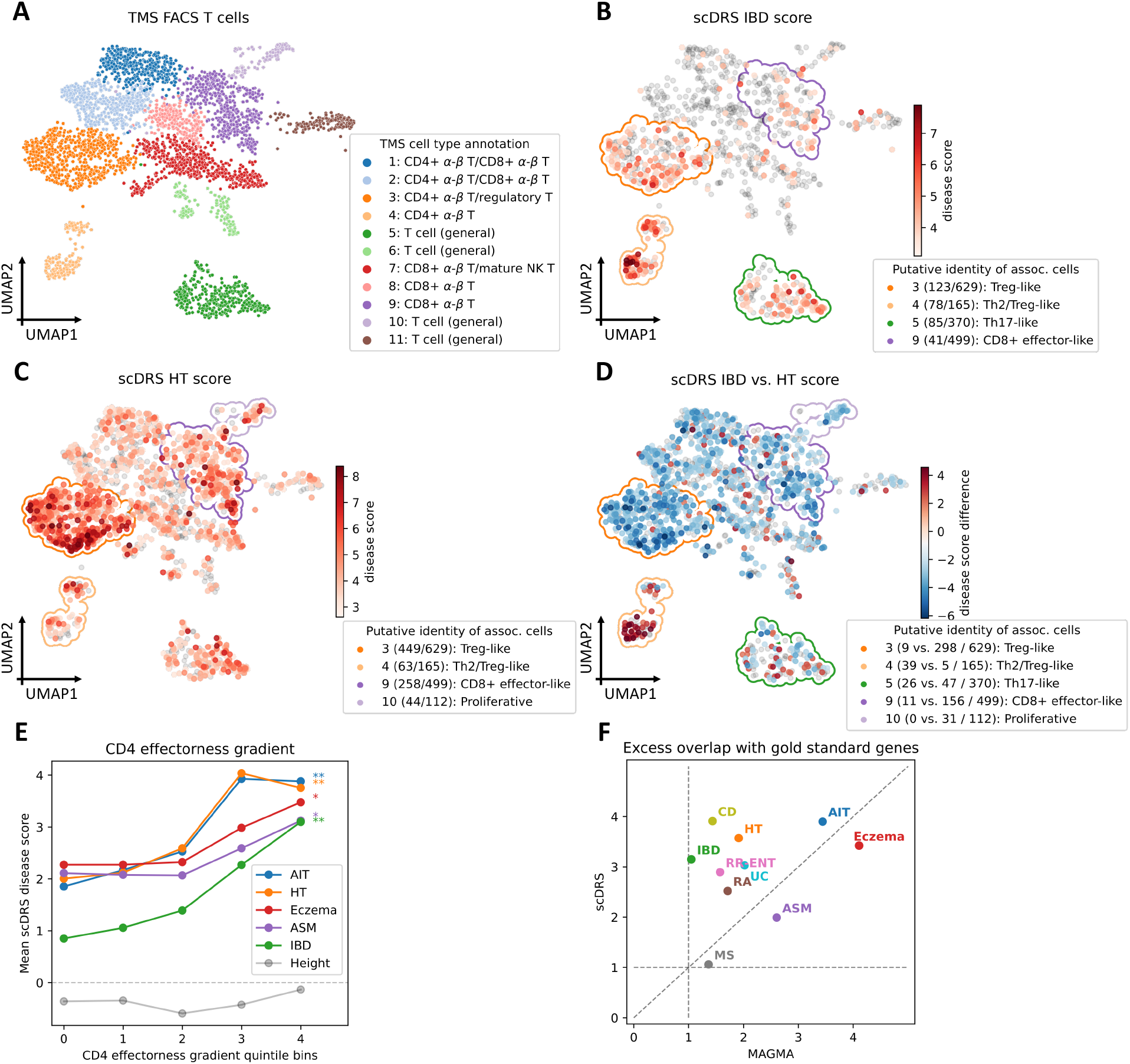
Associations of T cells with autoimmune diseases. **(A)** UMAP visualization of T cells in the TMS FACS data. In the legend, cluster labels are based on annotated TMS cell types in the cluster. Compositions of tissue, sex, and age of cells in each cluster are reported in Supp. Fig. 15. **(B-C)** Subpopulations of T cells associated with IBD and HT, respectively. Significantly associated cells (FDR*<*0.1) are denoted in red, with shades of red denoting `scDRS` disease scores; other cells are denoted in grey. Cluster boundaries indicate the corresponding T cell clusters from panel A. In the figure legend, the number of disease-associated cells and total number of cells are provided in parentheses, and cluster labels are based on the putative identities of the associated cells in the cluster, for the top 4 clusters (out of 11) with the strongest level of association (highest average disease score for associated cells in the cluster). Results for the other 8 autoimmune diseases and height are reported in Supp. Fig. 16. **(D)** Differences in individual cell-level associations between IBD and HT. Differentially associated cells (absolute `scDRS` disease score difference*>*2) are denoted in red and blue, with shades of colors denoting `scDRS` disease score differences; other cells are denoted in grey. Cluster boundaries indicate the corresponding T cell clusters from panel A. Clusters are annotated as in panels B and C; the number of IBD-enriched cells, HT-enriched cells, and all cells in the cluster are provided in parentheses. Differences in individual cell-level associations between IBD and the other 8 autoimmune diseases are reported in Supp. Fig. 21. **(E)** Association between `scDRS` disease score and CD4 effectorness gradient across CD4^+^ T cells for 5 representative autoimmune diseases and height, a negative control trait. The x-axis denotes CD4 effectorness gradient quintile bins and the y-axis denotes the average `scDRS` disease score in each bin for each disease. * denotes *P <*0.05 and ** denotes *P <*0.005 (MC test). Numerical results for all 10 autoimmune diseases are reported in Supp. Table 20. **(F)** Excess overlap of genes prioritized by `scDRS` with gold standard gene sets. The x-axis denotes the excess overlap of genes prioritized by MAGMA and the y-axis denotes the excess overlap of genes prioritize by `scDRS`, for each of 10 autoimmune diseases. The median ratio of (excess overlap − 1) for `scDRS` vs. MAGMA was 2.07. Numerical results are reported in Supp. Table 22.

We focused on individual cells associated with IBD, a representative autoimmune disease (Fig. 4B; results for HT in Fig. 4C; results for the other 8 autoimmune diseases and height in Supp. Fig. 16). The 387 IBD-associated cells (FDR*<*0.1) formed subpopulations of 4 of the 11 T cell clusters; we characterized these subpopulations based on marker gene expression, automatic T cell subtype annotation^42^, and overlap of specifically expressed genes in each subpopulation with T cell signature gene sets (Supp. Fig. 17,18,19,20; Methods). First, the subpopulation of 123 IBD-associated cells in cluster 3 (labeled as “Treg”) had high expression of regulatory T cell (Treg) marker genes (e.g., *FOXP3*^+^, *CTLA4*^+^, *LAG3*^+^; Supp. Fig. 20A), and their specifically expressed genes significantly overlapped with Treg signatures (*P* =6.0 × 10^−8^ for MSigDB signatures and *P* =4.0×10^−68^ for an effector-like Treg program^43^, Fisher’s exact test; Supp. Fig. 20C,D), suggesting these cells had Treg immunosuppressive functions. Interestingly, these 123 IBD-associated cells were non-randomly distributed in cluster 3 on the UMAP plot (*P <*0.001, MC test; Methods). Genes specifically expressed in these IBD-associated cells were preferentially enriched (compared to the 506 non-IBD-associated cells in the same cluster) in pathways defined by NF-*κ*B signaling, T helper cell differentiation, and tumor necrosis factor-mediated signaling (Supp. Fig. 20E); these pathways are closely related to inflammation, a distinguishing feature of IBD^44^. Second, the 78 IBD-associated cells in cluster 4 had high expression of T helper 2 (Th2) markers in the lower part of the cluster (e.g., *CCR8*^+^, *IL2*^+^) and Treg markers in the upper part (e.g., *FOXP3*^+^, *CTLA4*^+^; Supp. Fig. 17,18A-C), suggesting a mixed cluster identity (labeled as “Th2/Treg-like”); the role of Th2 cells in IBD has been discussed in literature^45^. Third, the 85 IBD-associated cells in cluster 5 (*IL23R*^+^ *RORC*^+^ *IL17A*^+^; labeled as “Th17-like”) were characterized as having T helper 17 (Th17) proinflammatory functions. Interestingly, drugs targeting *IL17A* (secukinumab and ixekizumab) have been considered for treatment of IBD but their use was associated with the onset of paradoxical effects (disease exacerbation after treatment with a putatively curative drug); the mechanisms underlying these events are not well understood^46^. Fourth, the 41 IBD-associated cells in cluster 9 (*IFNG*^+^ *GZMB*^+^ *FASL*^+^; labeled as “CD8^+^ effector-like”) were characterized as having effector CD8^+^ (cytotoxic) T cell functions. Overall, these findings are consistent with previous studies associating subpopulations of effector T cells to IBD, particularly Tregs and Th17 cells^44, 47–49^.

We further compared the individual T cell associations of IBD to HT, another representative autoimmune disease (Fig. 4C,D; results for comparison of IBD to the other 8 autoimmune diseases are reported in Supp. Fig. 21). The top 4 HT-associated subpopulations included 3 IBD-associated subpopulations (cells in clusters 3,4,9; Fig. 4C), but also a unique subpopulation of cells in cluster 10 (labeled as “proliferative”). The association strength was also different between the two diseases. Despite the stronger associations to HT overall (possibly due to higher GWAS power), IBD was more strongly associated with cells in cluster 4 (labeled as “Th2/Treg-like”; Fig. 4D). Across the 10 autoimmune diseases, pairwise `scDRS` disease score correlations (across all TMS FACS cells) were moderate (average of 0.51), implying differences between these diseases; the score correlations were not entirely driven by gene set overlap (average overlap of 231/1,000 genes; average `scDRS` disease score correlation of 0.16 when restricting to non-overlapping genes, substantially higher than the average of -0.10 across traits in different categories; Supp. Fig. 22, Supp. Table 19). Furthermore, the 10 autoimmune diseases formed 3 clusters based on hierarchical clustering of `scDRS` disease score correlations: IBD-related (IBD, UC, CD), allergy-related (Eczema, ASM, RR-ENT), and others (MS, RA, AIT, HT) (Supp. Fig. 22); these 3 groups represent biologically more similar subtypes of autoimmune diseases^50^, suggesting that `scDRS` can differentiate between subgroups of diseases from the same category.

We investigated whether the heterogeneity of T cells in association with autoimmune diseases was correlated with T cell effectorness gradient, a continuous classification of T cells defined by naive T cells on one side (immunologically naive T cells matured from the thymus) and effector T cells on the other (differentiated from naive T cells upon activation and capable of mediating effector immune responses); we hypothesized that such a correlation might exist given the effector-like T cell subpopulations associated to IBD above. Following a recent study^51^, we separately computed the effectorness gradients for CD4^+^ T cells (1,686 cells) and CD8^+^ T cells (2,197 cells) using pseudotime analysis^52^ (Supp. Fig. 23A,B; Methods), and confirmed that the inferred effectorness gradients were significantly negatively correlated with naive T cell signatures and positively correlated with memory and effector T cell signatures (Supp. Fig. 23C,D; Methods). We assessed whether the CD4 (resp., CD8) effectorness gradient was correlated with `scDRS` disease scores for IBD or other autoimmune diseases, across CD4^+^ T cells (resp., CD8^+^ T cells). Results are reported in Fig. 4E and Supp. Table 20. We determined that the CD4 effectorness gradient was strongly associated with IBD, CD, UC, AIT, and HT (*P <*0.005, MC test; 15%-28% of variance in `scDRS` disease score explained by CD4 effectorness gradient), weakly associated with Eczema and ASM (*P <*0.05; 6%-9% variance explained), but not significantly associated with RA, MS, or RR-ENT. This implies that these autoimmune diseases are associated with more effector-like CD4^+^ T cells. We also determined that the CD8 effectorness gradient was weakly associated with IBD, CD, and AIT (*P <*0.05, MC test; 6%-9% variance explained), but not significantly associated with the other autoimmune diseases, suggesting that CD4^+^ effector T cells may be more important than CD8^+^ effector T cells for these diseases. Notably, after conditioning on the 11 cluster labels, the associations with CD4 effectorness gradient remained significant for IBD and CD (*P <*0.005, MC test), AIT and HT (*P <*0.05), and the associations with CD8 effectorness gradient remained significant for IBD and CD (*P <*0.05), indicating that `scDRS` distinguishes effectorness gradients within clusters. In addition, as a negative control, height was not significantly associated in any of these analyses. The association of T cell effectorness gradients with autoimmune diseases has not previously been formally evaluated, but is consistent with previous studies linking T cell effector functions to autoimmune disease^53, 54^; the results also suggest that different subpopulations of effector T cells share certain similarities in their association with autoimmune diseases, consistent with previous studies characterizing the similarities among different subtypes of effector T cells, such as an increase in the expression of cytokines and chemokines^51, 55, 56^.

Finally, we prioritized disease-relevant genes by computing the correlation (across all TMS FACS cells) between the expression of a given gene and the `scDRS` score for a given disease; this approach identifies genes that are co-expressed with genes implicated by disease GWAS. We compared the top 1,000 genes prioritized using this approach with gold-standard disease-relevant genes based on putative drug targets from Open Targets^57^ (phase 1 or above; 8 gene sets with 27-250 genes; used for 8 autoimmune diseases except RR-ENT and HT; Supp. Table 21) or genes known to cause a Mendelian form of the disease^58^ (550 genes corresponding to “immune dysregulation”, used for RR-ENT and HT; Supp. Table 21). Results are reported in Fig. 4F and Supp. Table 22. We determined that `scDRS` attained a more accurate prioritization of disease-relevant genes compared to the top 1,000 MAGMA genes (median ratio of (excess overlap −1) was 2.07, median ratio of −log_10_ p-value was 2.86; Methods), likely by capturing disease-relevant genes with weak GWAS signal^59^. For example, *ITGB7* was prioritized by `scDRS` for association with IBD (rank 11) but was missed by MAGMA (rank 10565, MAGMA *P* =0.54); *ITGB7* impacts IBD via controlling lymphocyte homing to the gut and is a drug target for IBD (using vedolizumab)^60, 61^. In addition, *JAK1* was prioritized by `scDRS` for association with RA (rank 358) but was missed by MAGMA (rank 5228, MAGMA *P* =0.26); *JAK1* plays a role in regulating immune cell activation and is a drug target for RA (using tofacitinib, baricitinib, or upadacitinib)^62, 63^.

Additional secondary analyses are reported in the in Supp. Note, including replication results in 2 human scRNA-seq data sets^51, 64^ (Supp. Table 23), comparison to cluster-level LDSC-SEG analysis (Supp. Fig. 24), and additional results on prioritization of disease-relevant genes (Supp. Fig. 25, Supp. Table 21).

### Heterogeneous subpopulations of neurons associated with brain-related diseases and traits

We sought to further understand the heterogeneity across neurons (in the non-myeloid brain tissue) in the TMS FACS data (484 cells labeled as “neuron”) in association with brain-related diseases and traits (Fig. 3). We considered 7 brain-related diseases and traits: SCZ, MDD, BP, neuroticism (NRT), ECOL, BMI, Smoking (Supp. Table 1); we also considered height as a negative control trait. While these traits were broadly associated with neurons, pairwise `scDRS` disease score correlations were moderate across all TMS FACS cells (average of 0.44; Supp. Fig. 22, Supp. Table 19); there were also notable differences in associated brain cell populations, e.g., oligodendrocytes produced stronger associations for SCZ than for Smoking (Supp. Fig. 26). The TMS FACS data includes a partition of neurons into four brain subtissues (cerebellum, cortex, hippocampus, and striatum), but significant heterogeneity remained when we stratified our heterogeneity analyses by subtissue (Supp. Fig. 27).

Since the TMS FACS data has limited coverage of neuronal subtypes, we focused our subsequent analyses on a separate mouse brain scRNA-seq data set (Zeisel & Muñoz-Manchado et al.^65^; 3,005 cells), which has better coverage of neuronal subtypes and has been analyzed at cell type level in several previous genetic studies^8, 26, 66^. We first investigated cell type-trait associations using `scDRS`, which associated several neuronal subtypes (CA1 pyramidal neurons, SS pyramidal neurons, and interneurons) with the 7 brain-related traits (Supp. Fig. 28A, Supp. Table 24), consistent with previous genetic studies^8, 26, 66^. We focused on the CA1 pyramidal neurons from the hippocampus (827 cells), which exhibited the strongest within-cell type heterogeneity (FDR*<*0.005 for all 7 brain traits, MC test; Supp. Table 24). Individual cell-trait associations for SCZ are reported in Fig. 5A (results for all 7 brain-related traits in Supp. Fig. 28B). We observed a continuous gradient of CA1 pyramidal neuron-SCZ associations, with similar results for other traits.

**Figure 5.**
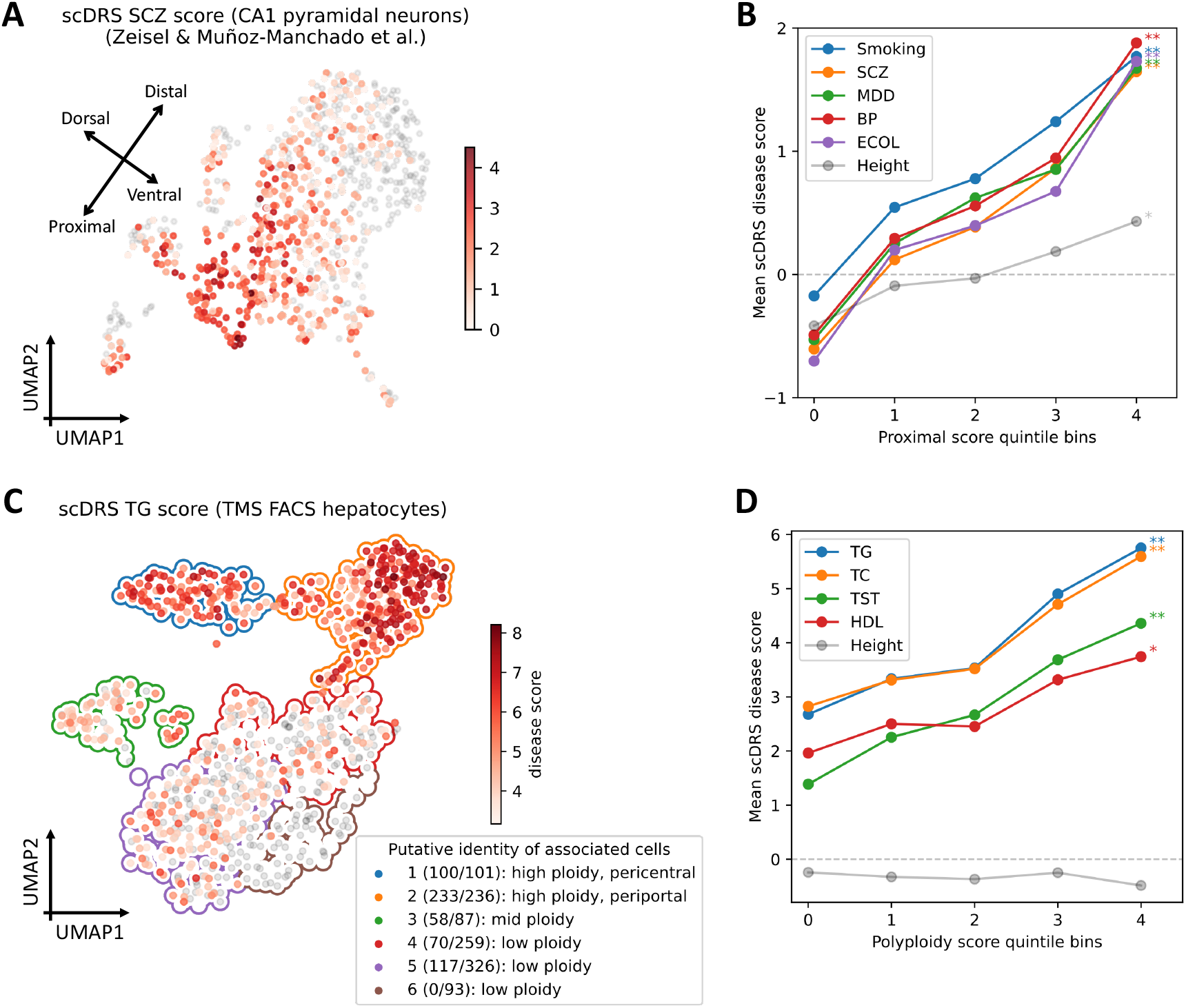
Associations of neurons with brain-related disease/traits and hepatocytes with metabolic traits. **(A)** Subpopulations of CA1 pyramidal neurons associated with SCZ in the Zeisel & Muñoz-Manchado et al. data. Colors of cells denote `scDRS` disease scores (negative disease scores are denoted in grey). We include a visualization of putative dorsal-ventral and proximal-distal axes (see text). Results for all 7 brain-related diseases/traits and height are reported in Supp. Fig. 28B. **(B)** Association between `scDRS` disease score and proximal score across CA1 pyramidal neurons for 5 representative brain-related disease/traits and height, a negative control trait. The x-axis denotes proximal score quintile bins and the y-axis denotes average `scDRS` disease score in each bin for each disease. * denotes *P <*0.05 and ** denotes *P <*0.005 (MC test). Results for all 6 spatial scores and all 7 brain traits (and height) are reported in Supp. Fig. 30 and Supp. Table 25. **(C)** Subpopulations of hepatocytes associated with TG in the TMS FACS data. Significantly associated cells (FDR*<*0.1) are denoted in red, with shades of red denoting `scDRS` disease scores; non-significant cells are denoted in grey. Cluster boundaries indicate the corresponding hepatocyte clusters. In the legend, numbers in parentheses denote the number of TG-associated cells vs. the total number of cells and cluster labels are based on the putative identity of cells in the cluster. Results for the other 8 metabolic traits and height are reported in Supp. Fig. 31. **(D)** Association between `scDRS` disease score and polyploidy score for 4 representative metabolic traits and height, a negative control trait. The x-axis denotes polyploidy score quintile bins and the y-axis denotes average `scDRS` disease score in each bin for each disease. * denotes *P <*0.05 and ** denotes *P <*0.005 (MC test). Results for all 3 scores (polyploidy score, pericentral score, periportal score) and all 9 metabolic traits (and height) are reported in Supp. Fig. 34 and Supp. Table 26.

We investigated whether the heterogeneity observed in Fig. 5A was correlated with spatial location; we hypothesized that such a correlation might exist because of the known location-specific functions of hippocampal neurons^17, 67^. We inferred spatial coordinates of the CA1 pyramidal neurons along the natural CA1 spatial axes^68^ (dorsal-ventral long axis, proximal-distal transverse axis, and superficial-deep radial axis) for each cell in terms of continuous individual cell-level scores for these 6 spatial regions by applying `scDRS` to published spatial signature gene sets (instead of MAGMA putative disease gene sets; Supp. Fig. 28C, Supp. Table 10; Methods). We verified that this procedure produced spatial scores significantly correlated with annotated spatial coordinates in independent mouse and human data sets^69, 70^ (Supp. Fig. 29). The inferred spatial scores for the long (dorsal, ventral) and transverse (proximal, distal) axes varied along the top two UMAP axes, providing visual evidence of stronger neuron-SCZ associations in dorsal and proximal regions (Fig. 5A, Supp. Fig. 28).

We used the results of `scDRS` for individual cells to assess whether the inferred spatial scores for each of the 6 spatial regions (dorsal/ventral/proximal/distal/superficial/deep) were correlated to the `scDRS` disease scores for each of the 7 brain-related traits (and height, a negative control trait) across CA1 pyramidal neurons (Methods). Results are reported in Fig. 5B (for the proximal region, which had the strongest associations), Supp. Fig. 30, and Supp. Table 25. We determined that the proximal score was strongly associated with all 7 brain-related traits (all *P <*0.002, MC test; 15%-29% of `scDRS` disease score variance explained by proximal score; *P* =0.006 for height), suggesting proximal CA1 pyramidal neurons may be more relevant to these brain-related traits (instead of distal CA1 pyramidal neurons). The association between the proximal region and brain-related traits is consistent with the fact that the proximal region of the hippocampus receives synaptic inputs in the perforant pathway, which is the main input source of the hippocampus^71, 72^.

We reapplied `scDRS` to 3 additional mouse single-cell data sets^69, 73, 74^ and 3 human single-cell data sets^70, 75, 76^ (Supp. Table 2), computing both spatial scores and disease scores for each cell as above. Results are reported in Supp. Fig. 30. We determined that the proximal score was consistently associated with the 7 brain-related traits across these 7 data sets (while the distal score was consistently non-associated). For the long (dorsal-ventral) and radial (superficial-deep) axes, while the dorsal and deep scores were consistently associated with the 7 brain-related traits across the 7 data sets, the corresponding ventral and superficial scores were consistently associated across the 3 human data sets but consistently non-associated across the 4 mouse data sets, possibly due to differences in brain biology between human and mouse^67, 77^.

### Heterogeneous subpopulations of hepatocytes associated with metabolic traits

We sought to further understand the heterogeneity across hepatocytes (in the liver) in the TMS FACS data in their association with metabolic traits (Fig. 3). Since the original study clustered all cells from the liver together^19^ (limiting the resolution for distinguishing cell states within hepatocytes), we reclustered the hepatocytes alone, resulting in 6 clusters (1,102 cells; Fig. 5C; Methods). We considered 9 metabolic traits: TG, high-density lipoprotein (HDL), low-density lipoprotein (LDL), total cholesterol (TC), TST, alanine aminotransferase (ALT), alkaline phosphatase (ALP), sex hormone-binding globulin (SHBG), and total bilirubin (TBIL) (Supp. Table 1); we also considered height as a negative control trait.

We focused on individual cells associated with TG, a representative metabolic trait (Fig. 5C; results for the other 8 metabolic traits and height in Supp. Fig. 31). The 530 TG-associated cells (FDR*<*0.1) formed subpopulations of 5 of the 6 hepatocyte clusters; we characterized these subpopulations based on ploidy level (number of sets of chromosomes in a cell) and zonation (pericentral/mid-lobule/periportal spatial location in the liver lobule), which have been extensively investigated in previous studies of hepatocyte heterogeneity^78–80^. We inferred the ploidy level and zonation for each individual cell in terms of a polyploidy score, a pericentral score, and a periportal score by applying `scDRS` to published polyploidy/zonation signature gene sets^81–83^ (instead of MAGMA putative disease gene sets; Supp. Fig. 32; Methods); we validated these inferred scores using expression signatures and independent data sets with experimentally determined annotations of ploidy level^82^ and zonation^83^ (Supp. Note). The inferred ploidy level and zonation varied across clusters, providing visual evidence of stronger cell-TG associations in high-ploidy clusters (clusters 1,2), particularly the periportal high-ploidy cluster (cluster 2; Fig. 5C). We further compared the associations between the 9 metabolic traits. While these traits were broadly associated with hepatocytes, pairwise `scDRS` disease score correlations were moderate across all TMS FACS cells (average of 0.36; Supp. Fig. 22, Supp. Table 19); however, there were no notable differences in the associated hepatocyte subpopulations across traits (Supp. Fig. 33).

We used the results of `scDRS` for individual cells to assess whether the inferred polyploidy, pericenteral and periportal scores were correlated to the `scDRS` disease score for each of the 9 metabolic traits (and height, a negative control trait) across hepatocytes; we jointly regressed the `scDRS` disease score for each trait on the polyploidy score, pericentral score, and periportal score (because the polyploidy score was positively correlated with the other 2 scores; Methods). Results are reported in Fig. 5D (for the polyploidy score which had the strongest associations), Supp. Fig. 34 and Supp. Table 26. The polyploidy, pericentral, and periportal scores jointly explained 42%-62% of variance of the `scDRS` disease scores across the 9 metabolic traits. We determined that the polyploidy score was strongly associated with all 9 metabolic traits (all *P <*0.005 except *P* =0.006 for HDL and *P* =0.007 for LDL, MC test; *P* =0.63 for height), suggesting that high-ploidy hepatocytes may be more relevant to these metabolic traits. The association between ploidy level and metabolic traits is consistent with previous findings that ploidy levels are associated with changes in the expression level of genes for metabolic processes such as de novo lipid biosynthesis and glycolysis^80, 81^, and supports the hypothesis that liver functions are enhanced in polyploid hepatocytes^80^. In addition, the periportal score was associated with the 9 metabolic traits (*P <*0.005 for TC, TST, ALP, MC test; all *P <*0.05 except *P* =0.24 for TBIL; *P* =0.24 for height). While the pericentral score was not significantly associated with these traits in the TMS FACS data, we detected significant associations across multiple other data sets (see below). These results suggest that these metabolic traits are impacted by complex processes involving both pericentral and periportal hepatocytes.

The association between hepatocyte ploidy level and metabolic traits may imply that there are metabolic trait GWAS variants associated with ploidy (ploidyQTL). This is supported by the excess overlap between the metabolic trait GWAS gene sets and a polyploidy signature gene set^81^, but is difficult to assess directly as genetic studies of ploidy level have largely focused on organisms other than humans^84^. Further details are provided in the Supp. Note, including results on 5 additional mouse and human data sets^19, 82, 83, 85, 86^ and validation of the polyploidy score using independent signature gene sets^81^ (Supp. Fig. 34 and Supp. Table 27).

## Discussion

We have introduced `scDRS`, a method that leverages polygenic GWAS signals to associate individual cells in scRNA-seq data with diseases and complex traits; we showed via extensive simulations that `scDRS` is well-calibrated and powerful. We applied `scDRS` to 74 diseases and complex traits in conjunction with 16 scRNA-seq data sets and detected extensive heterogeneity in disease associations of individual cells within classical cell types, including subpopulations of T cells associated with IBD partially characterized by their effector-like states, subpopulations of neurons associated with SCZ partially characterized by their spatial location, and subpopulations of hepatocytes associated with TG partially characterized by their higher ploidy levels. These findings have improved our understanding of these diseases/traits, and may prove useful for targeting the relevant cell populations for in vitro experiments to elucidate the molecular mechanisms through which GWAS risk variants impact disease. To ensure a reasonable number of `scDRS` discoveries, we recommend using GWAS data with a heritability z-score greater than 5, or sample size greater than 100K if heritability z-score is not available (although less stringent thresholds can be used for less polygenic traits) (Supp. Fig. 35). We also recommend using single-cell RNA-seq data with a diverse set of cells potentially relevant to disease, although a smaller number of cells should not affect the `scDRS` power. However, `scDRS` will not produce false positives for less ideal GWAS or single-cell data sets.

`scDRS` does not rely on annotations of classical cell types based on known marker genes, a standard approach for integrating GWAS with scRNA-seq data^6–8^ (and bulk gene expression data^9–12^; see Supp. Note), because the `scDRS` analysis uses the gene expression levels measured in individual cells. Thus, `scDRS` is particularly well-suited for analyzing data sets that are less well-annotated (e.g., large-scale cell atlases^19, 25^) or contain less well-studied cell populations. In addition, `scDRS` characterizes heterogeneity across individual cells in their associations to common diseases and complex traits, providing a unique perspective relative to studies of single-cell transcriptional heterogeneity focusing on scRNA-seq data alone^13–16, 18, 87, 88^; it also improves upon recent methods for scoring individual cells with respect to a given gene set (e.g., Seurat^15^, Vision^16^, and VAM^18^) by providing robust individual cell-level association p-values and higher detection power (see Supp. Note).

We have demonstrated the value of `scDRS` in associating individual cells to disease; assessing the heterogeneity across individual cells within predefined cell types in their association to disease; identifying cell-level variables partially characterizing the individual cells that are associated to disease; and broadly associating predefined cell types to disease. We anticipate that application of `scDRS` to future scRNA-seq/snRNA-seq and GWAS data sets will continue to further these goals.

We note several limitations and future directions of our work. First, identifying a statistical correlation between individual cells (or cell types) and disease does not imply causality, but may instead reflect indirect tagging of causal cells/cell types, analogous to previous work^6, 7, 12, 20^. However, even in such cases, the implicated cells/cell types are likely to be closely biologically related to the causal cells/cell types, based on their similar expression patterns. Second, we identified putative disease genes using MAGMA, a widely used method^20^. However, `scDRS` can be applied to any disease gene sets and gene weights, and it may be possible to construct more accurate sets of disease genes by incorporating other types of data, such as eQTL data^89^, protein-protein interaction data^90^ or functionally informed SNP-to-gene linking strategies^91^; we caution that such efforts must strive to avoid biases towards well-studied tissues. Third, since results of `scDRS` depend on the set of cells (and cell types) in the data set, it is appropriate to interpret the results with respect to other cells (or cell types) in the data set. We have implemented an option to adjust for cell type proportions (or any cell group annotations) so that the results will only depend on the set of cell types in the data set (but not the number of cells of each cell type), analogous to other disease-cell type association methods^7, 8, 12^; this option is recommended only for extremely unbalanced data sets (see Results and Methods). Fourth, while we have primarily focused on the associations involving a single disease/trait, further investigation of differences between diseases/traits within the same category is an important future direction. Please see more discussions, including use of mouse vs. human single-cell data, in Supp. Note. Despite all these limitations, `scDRS` is a powerful method for distinguishing disease associations of individual cells in single-cell RNA-seq data.

## Methods

### scDRS method

We consider a scRNA-seq data set with *n*_cell_ cells (not cell types) and *n*_gene_ genes. We denote the cell-gene matrix as 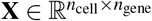, where *X*_*cg*_ represents the expression level of cell *c* and gene *g*. We assume that **X** is size-factor-normalized (e.g., 10,000 counts per cell) and log-transformed (log(*x* + 1)) from the original raw count matrix^21^. We regress the covariates out from the normalized data^21^ (with a constant term in the regressors to center the data), before adding the original log mean expression of each gene back to the residual data. Such a procedure preserves the mean-variance relationship in the covariate-corrected data, which is needed for estimating the gene-specific technical noise levels (see Supp. Note). Please see Supp. Fig. 2 for distributions of gene-level statistics for the TMS FACS, TMS droplet, and TS FACS data (gene-level statistics for all 16 data sets are reported in Supp. Table 3). The technical noise levels are moderately correlated across genes between the 16 data sets (avg. cor. 0.34) and are highly correlated between data sets with similar cell type compositions (e.g., 0.74 between TMS FACS and TS FACS; Supp. Table 4).

The `scDRS` algorithm is described in Box 1. Given a disease GWAS and an scRNA-seq data set, `scDRS` computes a p-value for each individual cell for association with the disease. `scDRS` also outputs cell-level normalized disease scores and *B* sets of normalized control scores (default *B* =1,000) that can be used for data visualization and Monte Carlo-based statistical inference (see Downstream applications and MC test). `scDRS` consists of three steps. First, `scDRS` constructs a set of putative disease genes from the GWAS summary statistics. Second, `scDRS` computes a raw disease score and *B* MC samples of raw control scores for each cell. Third, after gene set-wise and cell-wise normalization, `scDRS` computes an association p-value for each cell by comparing its normalized disease score to the empirical distribution of the pooled normalized control scores across all control gene sets and all cells. These steps are detailed below.

#### Step 1: Constructing disease gene set

We use MAGMA^20^ to compute gene-level association p-values from disease GWAS summary statistics (Box 1, step 1). We use a reference panel based on individuals of European ancestry in the 1000 Genomes Project^92^. We use a 10-kb window around the gene body to map SNPs to genes. We select the top 1,000 genes based on MAGMA p-values as putative disease genes and use their MAGMA z-scores as the GWAS gene weights. We denote the disease gene set as *G* ⊂{1, 2, …, *n*_gene_} and their GWAS gene weights as {*w*_*g*_}_*g*∈*G*_. Alternative parameter choices and methods for constructing putative disease gene sets are considered below (see Alternative versions of `scDRS` method).

#### Step 2: Computing disease scores and control scores

We construct *B* sets of control genes 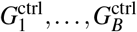 by randomly selecting genes matching the mean expression and expression variance of the disease genes calculated across all cells in the data set (Box 1, step 2a). Specifically, each control gene set 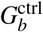 has the same size as the disease gene set *G* and is constructed by first dividing all genes into 20×20 equal-sized mean-variance bins and then for each gene in the disease gene set, randomly sampling a control gene from the same bin (containing the disease genes) without replacement. Next, we estimate the technical noise level for each gene 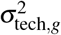 in the scRNA-seq data, the part of the variance due to sequencing noise, using a procedure similar to previous works^18, 21^ by modeling the mean-variance relationship across genes; we further compute the raw disease score and raw control scores for each cell as weighted average expression of genes in the corresponding gene set (Box 1, steps 2b-2c, Supp. Note). The weight for gene *g* is proportional to 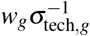 (capped at 10 for both the MAGMA z-score *w*_*g*_ and single-cell weight 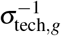), which upweights genes with stronger GWAS associations and downweights genes with higher levels of technical noise to increase detection power. The single-cell weight 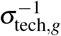 was adapted from VAM^18^, where the cell-specific score is proportional to 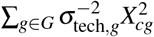 and was shown to have a superior classification accuracy. Alternative cell scores (instead of the weighted average score) are evaluated below (see Alternative versions of `scDRS` method).

##### Box 1

Single-cell disease relevance score (`scDRS`)

**Input:** Disease GWAS summary statistics (or putative disease gene set *G* with GWAS gene weights {*w*_*g*}*g*∈*G*_), scRNA-seq data 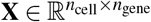.

**Parameters:** Number of MC samples of control gene sets *B* (default 1,000).

1. **Construct putative disease gene set**
  a. Construct putative disease gene set *G* ⊂ {1, 2, …, *n*_gene_} with GWAS gene weights {*w*_*g*_} _*g*∈*G*_ from GWAS summary statistics using MAGMA.
2. **Compute disease scores and control scores**
  a. Sample *B* sets of control genes 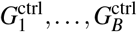 matching mean expression and expression variance of disease genes.
  b. Estimate gene-specific technical noise level 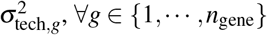.
  c. Compute raw disease score and *B* raw control scores for each cell *c* = 1, …, *n*_cell_,

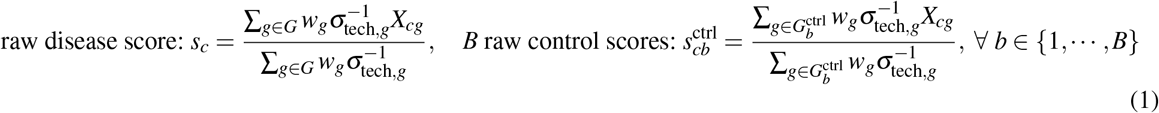
3. **Compute disease association p-values**
  a. First gene set alignment by mean and variance. Let 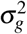 be the expression variance of gene *g*. For each cell *c*,

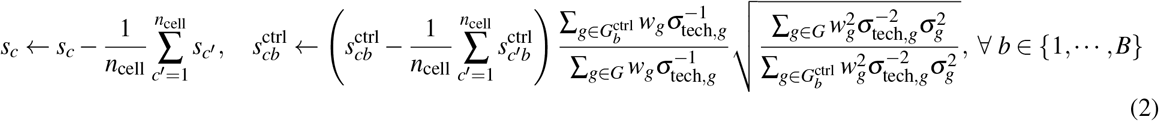
  b. Cell-wise standardization for each cell *c* by the mean 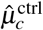 and variance 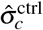 of control scores 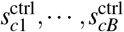 of that cell,

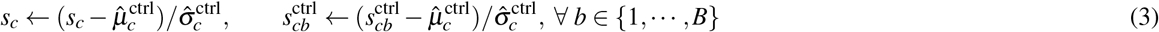
  c. Second gene set alignment by mean. For each cell *c*,

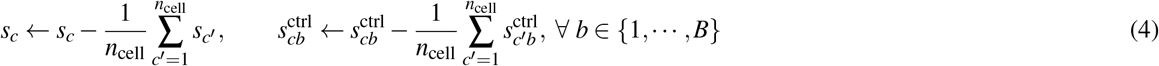
  d. Compute cell-level p-values based on the empirical distribution of the pooled normalized control scores for each cell *c*,

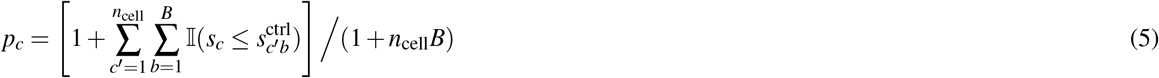

**Output:** cell-level p-values *p*_*c*_, normalized disease scores *s*_*c*_, and normalized control scores 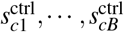.

#### Step 3: Computing disease-association p-values

We first describe the alternative distribution that `scDRS` aims to detect. Since the control genes match the mean expression and expression variance of the disease genes across cells, it can be shown that the raw disease score has the same mean but a higher variance compared to each set of raw control scores; the higher variance is because the disease genes are more positively correlated with each other due to co-expression in the associated cell population (Supp. Fig. 1A-C). As a result, the disease-relevant cells, with high expression of the disease genes, are expected to have larger raw disease scores than raw control scores. We caution that the disease genes may be more positively correlated due to other reasons such as being physically close to each other, but `scDRS` will produce much weaker signals in these cases (Supp. Fig. 5). Please see more details in Supp. Note.

The first gene set alignment (Box 1, step 3a) corrects for the potential mismatch of control gene sets by first centering the scores and then aligning the variance level for each gene set. The variance of the raw disease score is estimated as 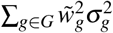 and similarly for the raw control scores, with 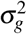 being the expression variance of gene *g* and 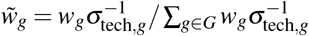 the corresponding weight; this heuristic assumes independence of the genes (or different gene sets have similar levels of gene-gene correlation), and consequently avoids downweighting the raw disease score due to the higher correlation between disease genes (Supp. Fig. 1D, Supp. Note). After adjusting the control gene sets, the gold standard MC p-values, based on comparison to *B* MC samples of raw control scores of the same cell, can be written as^22^

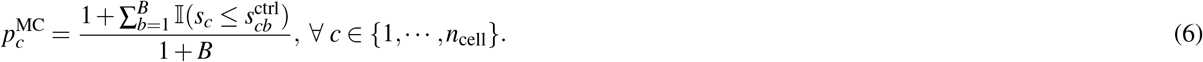

This finite-sample MC p-value is a conservative estimate of the ideal MC p-value obtained via an infinite number of MC samples^22^. However, as Eq. (6) suggests, an MC test with *B* MC samples can only produce an MC p-value no smaller than 1*/*(1 + *B*). Instead of using a large number of MC samples which is computationally intensive, we approximate the ideal MC p-value by pooling the control scores across cells. Specifically, we first align the control score distributions (across the *B* control gene sets, for each cell) by matching their means and variances, followed by re-centering the mean scores of different gene sets (Box 1, steps 3b-3c, Supp. Fig. 1E,F, Supp. Note). This procedure produces a normalized disease score and *B* normalized control scores for each cell. Finally, we compute the `scDRS` p-values based on the empirical distribution of the pooled normalized control scores across all control gene sets and all cells (Box 1, step 3d). The pooling procedure assumes that the raw control score distributions (across the *B* control gene sets, for each cell) are from the same location-scale family (e.g., the family of all normal distributions or that of all student’s t-distributions) such that they can be aligned by matching the first two moments; it is a reasonable assumption when the number of disease genes is neither too small nor too large (e.g., 50 *<* |*G*| *<* 20%*n*_gene_), where the control score distributions are close to normal distributions by the central limit theorem (Supp. Note). As shown in Supp. Fig. 1G-I, the `scDRS` p-values with *B* =1,000 is indeed able to well approximate the MC p-values obtained using a much larger number of MC samples (*B* =20,000).

### Downstream applications and MC test

`scDRS` outputs individual cell-level p-values, (normalized) disease scores, and (normalized) control scores that can be used for a wide range of downstream applications: assessing association between a given cell type and a given disease; assessing heterogeneity in association with a given disease across a given set of cells; and assessing association between a cell-level variable and a given disease across a given set of cells. We use a unified MC test for these 3 analyses based on the disease score and control scores. Specifically, let *t* be the test statistic computed from the disease score of the given set of cells (the 3 analyses differ by the test statistics they use) and let 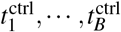 be the same test statistics computed from the *B* sets of control scores of the same set of cells. The MC p-value can be written as

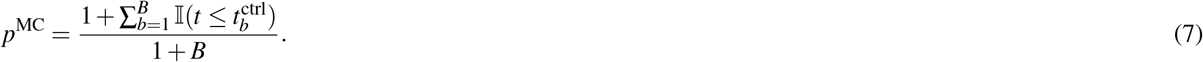

The MC test avoids the assumption that the cells are independent—a strong assumption in scRNA-seq analyses, e.g., when analyzing cells in the same cluster that are dependent due to the clustering process. We can also compute an MC z-score as 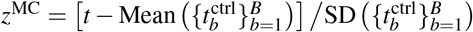 ; this MC z-score is not restricted by the MC limit of 1*/*(1 + *B*) but relies the assumption that the control test statistics 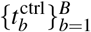 approximately follow a normal distribution. Below, we describe the test statistics used by the 3 analyses listed above. We note that the MC test can in principle be extended to any analysis that computes a test statistic from the disease scores of a set of cells.

### Assessing association between a given cell type and a given disease

We use the top 5% quantile of the disease scores of cells from the given cell type as the test statistic. This test statistic is robust to annotation outliers, e.g., a few misannotated but highly significant cells. One can also use other test statistics such as the top 1% quantile or the maximum.

### Assessing heterogeneity in association with a given disease across a given set of cells

We use Geary’s C^16, 93^ as the test statistic. Geary’s C measures the spatial autocorrelation of the disease score across a set of cells (e.g., cells from the same cell type or cell cluster) with respect to a cell-cell similarity matrix. Given a set of *n* cells, the corresponding disease scores *s*_1_, …, *s*_*n*_, and the cell-cell similarity matrix *W* ∈ ℝ^*n*×*n*^, Geary’s C is calculated as

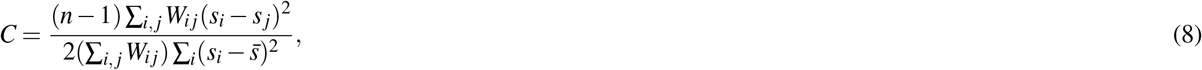

where 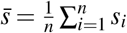. We use the cell-cell connectivity matrix for the similarity matrix like previous works^16^, which corresponds to the “connectivities” output from the scanpy function “scanpy.pp.neighbors”^94^. A value significantly lower than 1 indicates positive spatial autocorrelation, suggesting cells close to each other on the similarity matrix have similar disease scores, forming subclusters of cells with similar levels of disease association. This indicates a high level of disease association heterogeneity across the given set of cells. We use this test to assess within-cell type disease association heterogeneity and within-cluster association disease heterogeneity.

### Assessing association between a cell-level variable and a given disease across a given set of cells

For associating a single cell-level variable with disease, we use the Pearson’s correlation between the cell-level variable and the disease score across the given set of cells as the test statistic. For jointly associating multiple cell-level variables with disease, we use the regression *t*-statistic as the test statistic, obtained from jointly regressing the disease score against the cell-level variables.

### Computational cost

Both the computation time and memory use of `scDRS` scale linearly with the number of cells and the number of control gene sets (default 1,000). We performed benchmark experiments by subsampling cells from the Nathan et al. data^64^ and, as expected, observed a linear relationship between the number of cells and both the computation time and memory usage (Supp. Fig. 3). `scDRS` required 1.6 hours of computation time and 30GB of memory to process 500K cells under the default setting (1,000 control gene sets); it is estimated to take around 3 hours and 60GB of memory to run `scDRS` on a data set with a million cells and a similar level of sparsity. Of note, in this experiment, the memory usage is only 1.5X of the theoretical lower limit, namely 18.9G consisting of 11.4G for loading the data in high precision (64-bit float) and 7.5G for computing the 1,000 sets of raw and normalized control scores for each cell (2× 500,089 ×1,000 ×8B = 7.5G); the memory usage is 3X of the theoretical lower limit for low-precision computation. Therefore, `scDRS` is reasonably efficient in memory usage. Based on this benchmark experiment, we also suggest an empirical formula for estimating the memory usage (in the unit of GB) as 3 ×(low_precision_data_size + *n*_cell_*B* × 8*/*1024^3^).

### Simulations

We performed simulations on a data set with 10,000 cells subsampled from the TMS FACS data. In null simulations, we randomly selected putative disease genes from a set of non-informative genes. We considered four numbers of putative disease genes (100, 500, 1,000, or 2,000) and four types of genes to sample from: (1) the set of all genes, (2) the set of top 25% genes with high mean expression, (3) the set of top 25% genes with high expression variance, (4) the set of top 25% overdispersed genes, where the level of overdispersion is calculated as the difference between the actual variance and the estimated technical variance in the log scale data. For the default version of `scDRS`, we simulated GWAS gene weights by first randomly selecting a disease (out of the 74 diseases/traits) and then randomly permuting the top MAGMA z-scores from the selected disease. We did not simulate gene-specific technical noise-based single-cell weights because these weights were inherent to the single-cell data. For the MC test for cell type-disease association, we used the top 5% quantile as the test statistic and computed the MC p-values for each cell type and each set of random putative disease genes by comparing the test statistic from the disease scores to those computed from the 1,000 sets of control scores (see Monte-Carlo-based downstream analyses above). In causal simulations, we randomly selected 1,000 causal disease genes, randomly selected 500 of the 10,000 cells as causal cells and artificially perturbed their expression levels to be higher (at various effect sizes) across the 1,000 causal disease genes, and randomly selected 1,000 putative disease genes (provided as input to `scDRS` and other methods) with various levels of overlap with the 1,000 causal disease genes. Here, the effect size corresponds to the fold change of expression of the causal genes in the causal cells (multiplicative in the original count space and additive in the log space). We performed three sets of causal simulations: (1) varying effect size from 5% to 50% while fixing 25% overlap, (2) varying level of overlap from 5% to 50% while fixing 25% effect size, (3) assigning the 528 B cells in the subsampled data to be causal (instead of the 500 randomly selected cells; varying effect size while fixing 25% overlap). The FDR and power reported in Fig. 2B and Supp. Fig. 6 are based on applying the B-H procedure^95^ to all cells at nominal FDR=0.1. All experiments were repeated 100 times and confidence intervals were computed based on the normal distribution. We considered three methods for comparison, namely Seurat^15^ (“score_genes” as implemented in scanpy^94^), Vision^16^, and VAM^18^. To our knowledge, VAM is the only published cell-scoring method that provides cell-level association p-values. We chose to include Seurat due to its wide use and standardized its output cell-level scores (mean 0 and SD 1) before computing the cell-level p-values based on the standard normal distribution. We chose to include Vision because its outputs are nominal cell-level z-scores and can be easily converted to p-values; we again added the standardization step because otherwise the results of Vision were highly unstable. We did not include other methods like PAGODA^14^ or AUCell^14^ because it is not straightforward to convert their outputs to cell-level association p-values and also because the z-scoring methods (e.g., Vision) outperformed other methods in a comprehensive evaluation in Frost et al.^18^

### GWAS summary statistic data sets

We analyzed GWAS summary statistics of 74 diseases and complex traits from the UK Biobank^96^ (47 of the 74 diseases/traits with average *N*=415K) and other publicly available sources^32, 97–118^ (27 of the 74 diseases/traits with average *N*=225K); average *N*=346K for all 74 diseases/traits; Supp. Table 1). All diseases and traits were well-powered (heritability z-score*>*5), except celiac disease (Celiac), systemic lupus erythematosus (SLE), multiple sclerosis (MS), subject well being (SWB), and type 1 diabetes (T1D), which were included due to their clinical importance. The major histocompatibility complex (MHC) region was removed from all analyses because of its unusual LD and genetic architecture^119^.

### scRNA-seq data sets

We analyzed 16 scRNA-seq or snRNA-seq data sets (Supp. Table 2). We included 3 atlas-level data sets (TMS FACS, TMS droplet, and TS FACS) to broadly associate diverse cell types and cell populations to disease; these 3 data sets cover different species (mouse and human) and different technologies (FACS and droplet), which allows us to assess the robustness of our results across different species and technologies. We included another 13 data sets that focus on a single tissue and contain finer-grained annotations of cell types and cell states. Notably, several of these data sets contain experimentally determined annotations which allow us to better validate our results, including Cano-Gamez & Soskic et al. data^51^ containing experimentally perturbed CD4^+^ T cell states, Nathan et al. data^64^ containing T cells states determined by profiling surface markers using CITE-seq, Habib & Li et al. data^69^ containing experimentally determined spatial locations for CA1 pyramidal neurons based on ISH of spatial landmark genes, Ayhan et al. data^70^ containing experimentally determined spatial locations for CA1 pyramidal neurons (dorsal and ventral) based on surgical resection, and Richter & Deligiannis et al. data^82^ containing experimentally determined hepatocyte ploidy levels based on Hoechst staining.

### Adjusting for cell type proportions

`scDRS` can additionally take a set of cell type annotations (or any cell group annotations) and adjust for cell type proportions by inversely weighting cells by the number of cells in the corresponding cell type (weights were normalized to have mean 1 and were constrained between 0.1 and 10). This version of scDRS generated highly consistent as the default version in the TMS FACS data (median of 0.97 across 74 traits for the disease score correlation computed across all TMS FACS cells) and was well-calibrated in null simulations (Supp. Fig. 4). We recommend the use of this new option only for extremely unbalanced data sets, for 3 reasons. First, it produced consistent results for relatively balanced data sets such as TMS FACS. Second, it requires cell type annotations where the cell types have a similar level of granularity (e.g., B cells vs. T cells instead of B cells vs. a subtype of CD4^+^ Th17 cells), which is not always available. For example, the TMS cell type annotation contains both high-level cell types like T cells and more fine-grained cell types like Tregs. Third, the cell type annotation can be defined with different levels of granularity, such as broader types like immune cells or very specific types like CD4^+^ Th17 cells, and it is unclear how to choose the right level of granularity for a given data set.

### Comparison with other cell type-level association methods

We briefly discuss the similarities and differences between `scDRS` and 3 cell type-level association methods that also make use of MAGMA: the MAGMA-based method in Skene et al.^26^, Watanabe et al.^7^, and the MAGMA-based method in Bryois et al.^8^ All statements about `scDRS` apply to both individual cell level-analysis and cell type-level analysis. First, all 4 methods focus on specifically-expressed genes in a cell type (or cell) rather than merely highly-expressed genes. Second, all 4 methods produce results that depend on cell types (cells) present in the data set, so it is important to interpret the results with respect to other cell types (cells) in the data set. Third, `scDRS` and the method of Watanabe et al. depend on different scaling factors for size factor normalization (while the other 2 methods do not). However, we determined this step is crucial for removing confounding effects, and `scDRS` is not sensitive to different choices of scaling factors (Results). Fourth, the other 3 methods use linear regression to associate MAGMA z-scores with cell type features across genes (cell type expression level in Watanabe et al.; cell type specificity in Skene et al. and Bryois et al.). `scDRS` can be viewed as a non-parametric alternative to these methods, employing a stratified permutation test that associates MAGMA z-scores for top genes with expression levels for a given cell by permuting genes within each level of expression mean and variance. Thus, unlike the other 3 methods, `scDRS` does not rely on a linearity assumption. Fifth, `scDRS` may be more powerful when there is within-cell type heterogeneity in association to disease. Further details are provided in the Supp. Note.

### Alternative versions of scDRS method

We considered alternative versions of `scDRS`, involving (1) other choices of MAGMA gene window size, (2) other strategies for selecting putative disease genes, (3) other methods for choosing gene weights for the selected putative disease genes, (4) an alternative overdispersion score (instead of weighted average), and (5) other methods for constructing putative disease genes. We considered 3 MAGMA gene window sizes for mapping SNPs to genes: 0 kb, 10 kb (default), and 50 kb. We considered 6 strategies for selecting putative disease genes: top 100, top 500, top 1,000 (default), top 2,000, FWER*<*5%, FDR*<*1% (multiple testing correction performed based on MAGMA p-values for each trait separately; number of top genes constrained between 100 and 2,000 for the latter two methods). We considered 4 methods for choosing gene weights for the selected putative disease genes: no weights, GWAS z-score weights (proportional to MAGMA z-score capped at 10), single-cell VS weights (proportional to reciprocal of technical noise level 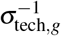 capped at 10), and using both sets of weights (default). We evaluated the performance based on a curated set of 20 traits with expected and unexpected disease-critical cell types; we caution that some cell types labeled as unexpected may still be relevant to disease despite not being implicated in the current literature (Supp. Fig. 12, Supp. Table 17). The default version of `scDRS` substantially outperformed all other versions except the version that uses the top 2,000 genes for the gene selection method. This latter version was not chosen as the default because it was not significantly better than using the top 1,000 genes and `scDRS` was less well-calibrated for gene sets with 2,000 genes.

The overdispersion score is defined as

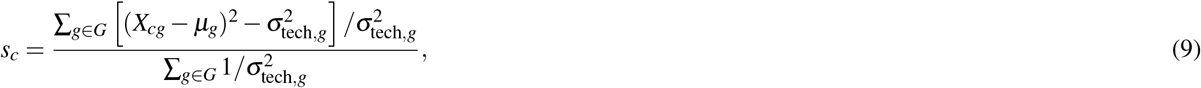

where *μ*_*g*_ and 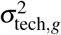 are the average expression and technical noise level of gene *g* respectively. The overdispersion score tests for both overexpression and underexpression of the putative disease genes in the relevant cell population (unlike the weighted average score which only tests for overexpression of the disease genes). We compared the overdispersion score to two versions of `scDRS` that only tested for overexpression: the default version (GWAS+single-cell weights) and the unweighted score. We assessed the performance in terms of number of significant discoveries in TMS FACS across the 74 diseases (details in Supp. Fig. 14).

We also discuss other methods for constructing putative disease genes. While we constructed putative disease genes using GWAS data and mapped SNPs to genes based on genomic locations, it may be possible to obtain a more accurate disease gene set by either incorporating data from other sources such as protein-protein interaction data^90^ or using a more sophisticated SNP-to-gene linking strategy^91^; exploring these approaches is an interesting future direction.

### Analysis of T cells and autoimmune diseases

We collectively analyzed all T cells from the TMS FACS data (4,125 cells labeled as CD4^+^ *α*-*β* T cell, CD8^+^ *α*-*β* T cell, regulatory T cell, mature NK T cell, mature *α*-*β* T cell, or T cell in the TMS data; Supp. Table 5); the more general terms like “T cell” and “mature *α*-*β* T cell” were used for cells whose more specific identities were not clear. We processed the T cells following the standard procedure using scanpy^94^. First, we performed size factor normalization (10,000 counts per cell) and log transformation. Second, we selected highly variable genes and computed the batch-corrected PCA embedding using Harmony^120^, treating each mouse as a batch. Finally, we constructed KNN graphs and clustered the cells using the Leiden algorithm^121^ (resolution=0.7), followed by computing the UMAP embedding. We removed 376 cells either from small clusters (less than 100 cells) or whose identities are ambiguous, resulting in 3,769 cells. We annotated the clusters based on the major TMS cell types in the cluster; the label “mature *α*-*β* T cell” was omitted because a more specific TMS cell type label (e.g., “CD8^+^ *α*-*β* T”) was available in the corresponding cluster. We considered cells from clusters 1-4 as clear CD4^+^ T cells (1,686 cells) and cells from clusters 1, 2, 7-9 as clear CD8^+^ T cells (2,197 cells; the shared clusters 1 and 2 contain a mix of naive CD4^+^ and CD8^+^ T cells). We used diffusion pseudotime (DPT)^52^ to assign effectorness gradient for CD4^+^ and CD8^+^ T cells separately, where we used the leftmost cell in cluster 2 on the UMAP as the root cell (clearly naive T cell).

To robustly annotate disease-associated T cell subpopulations, we performed 2 sets of automatic T cell subtype analyses: classification based on marker gene expression (details in Supp. Fig. 18) and automatic T cell states annotation using projecTILE^42^ (v2.0.2). The two sets of annotations were consistent for distinguishing effector vs. naive T cells and distinguishing CD4^+^ vs. CD8^+^ T cells, suggesting the results were overall consistent. Since the projecTILE reference contained a limited set of T cell subtypes (e.g., no Th2 or Th17 cells), we used the marker gene-based annotation for the main results. For the analysis of individual cells associated with IBD, we considered 4 major clusters of T cells with *>*25 IBD-associated cells (FDR*<*0.1). First, the subpopulation of 123 IBD-associated cells in cluster 3 (which consisted of 629 cells with TMS cell type labels “CD4^+^ *α*-*β* T” or “regulatory T”) were labeled as “Treg” as described in the main paper. Second, the 78 IBD-associated cells in cluster 4 (which consisted of 165 cells with TMS cell type label “CD4^+^ *α*-*β* T”) were labeled as “Th2/Treg-like” as described in the main paper. Their specifically expressed genes significantly overlapped with a *KLRG1*^+^ *AREG*^+^ effector-like Treg program^43^ characterized by high expression levels of *IL1RL1* (*ST2*), *KLRG1*, and *AREG* (*P* =1.3 × 10^−50^, Fisher’s exact test; Supp. Fig. 20D), suggesting these cells had active functions for Treg differentiation, immunosuppression, and tissue repair^43^. Third, the 85 IBD-associated cells in cluster 5 (which consisted of 370 cells with TMS cell type label “T cell”) were labeled as “Th17-like” as described in the main paper. Their specifically expressed genes significantly overlapped with Th17 signatures (*P* =2.0 × 10^−6^, Fisher’s exact test; Supp. Fig. 20C) and a Th17-like Treg program^43^ (*P* =1.9 × 10^−24^, Fisher’s exact test; Supp. Fig. 20D), suggesting Th17 proinflammatory functions. Finally, the 41 IBD-associated cells in cluster 9 (consisting of 499 cells with TMS cell type label “CD8^+^ *α*-*β* T”) were labeled as “CD8^+^ effector-like” as described in the main paper. Their specifically expressed genes significantly overlapped with effector CD8^+^ T cell signatures (*P* =1.6 × 10^−9^, Fisher’s exact test; Supp. Fig. 20C), suggesting cytotoxic T cell functions. For the analysis of individual cells associated with HT, the putative identities of HT-associated cells in clusters 3,4,9 were similar to the putative identities of IBD-associated cells in the corresponding clusters. The 44 HT-associated cells in cluster 10 (consisting of 112 cells with TMS cell type label “T cell”) were labeled as “Proliferative” due to high expression of proliferation markers (Supp. Fig. 18,20B).

We used MSigDB^122, 123^ (v7.1) to curate T cell signature gene sets, including naive CD4, memory CD4, effector CD4, naive CD8, memory CD8, effector CD8, Treg, Th1 (T helper 1), Th2 (T helper 2), and Th17 (T helper 17) signatures. For each T cell signature gene set, we identified a set of relevant MSigDB gene sets (22-34, Supp. Table 9), followed by selecting the top 100 most frequent genes in these MSigDB gene sets as the T cell signature genes; a gene was required to appear at least twice and genes appearing the same number of times were all included, resulting in 62 to 513 genes for the 10 T cell signature gene sets (Supp. Table 10). For gold-standard gene sets used in the analysis of disease gene prioritization, we curated 27 putative drug target gene sets from Open Targets^57^ (mapped to 27 of the 74 diseases/traits considered in the paper; Supp. Table 21); for a given disease, we selected all genes with drug score *>*0 (clinical trial phase 1 and above) and only considered diseases with at least 10 putative drug target genes. We curated 16 Mendelian diseases gene sets from Freund et al.^58^ (mapped to 45 of the 74 diseases/traits considered in the paper; Supp. Table 21). For comparison of two gene sets, the p-value is based on Fisher’s exact test and excess overlap is defined as the ratio between the observed overlap of the two gene sets and the expected overlap (by chance). Of note, for a given query gene set with a fixed size and a fixed level of excess overlap with the reference gene set, the −log_10_ p-value increases with the size of the reference gene set; we report both excess overlap and −log_10_ p-value while using the former as our primary metric, which is more interpretable.

### Analysis of neurons and brain-related diseases/traits

For the TMS FACS data, we focused on the 484 neurons (TMS label “neuron”, excluding cells with TMS label “medium spiny neuron” or “interneuron”). For the Zeisel & Muñoz-Manchado et al. data, we applied `scDRS` to all 3,005 cells and then focused on the 827 CA1 pyramidal neurons (“level1class” label “pyramidal CA1”). For inferring spatial coordinates, we curated differentially expressed genes for each of the 6 spatial regions (dorsal vs. ventral, ventral vs. dorsal, proximal vs. distal, distal vs. proximal, deep vs. superficial, and superficial vs. deep) using the gene expression data from Cembrowski et al.^68^ (GEO GSE67403; gene sets in Supp. Table 10). For each differential gene expression analysis, we selected genes based on FPKM*>*10 for the average expression in the enriched region (e.g., dorsal for the dorsal vs. ventral comparison), *q*-value*<*0.05, and log_2_(fold change) *>*2. We used `scDRS` and these signature gene sets to assign 6 spatial scores for each cell. For the regression analysis, we separately regressed the `scDRS` disease scores for each of the 7 brain-related traits (and height, a negative control trait) on each of the 6 spatial scores. We performed marginal regression instead of joint regression for these spatial scores because the inferred spatial scores for opposite regions on the same axis (e.g., dorsal vs. ventral) were highly collinear (strongly negatively correlated), and the inferred spatial scores for dorsal, proximal, and deep regions (which had strong marginal associations to diseases) had very low pairwise correlations (average |*r*| =0.10; Supp. Fig. 28D), suggesting these associations were independent. We reported correlation p-values (MC test) and variance explained for each of the 6 spatial scores.

### Analysis of hepatocytes and metabolic traits

We considered all hepatocytes in the TMS FACS data (1,162 cells) and reprocessed them following the same procedure as we did for the T cells. We further filtered out low-quality cells (mitochondrial proportion ≥0.3; likely to be apoptotic or lysing cells), resulting in 1,102 hepatocytes (Fig. 5C). We curated signature gene sets for ploidy level, zonation, and putative zonated pathways. We curated 4 sets of polyploidy signatures, including differentially expressed genes (DEGs) for partial hepatectomy (PH) vs. pre-PH^81^ (used for the polyploidy score), Cdk1 knockout (case) vs. control^81^, 4n vs. 2n hepatocytes^82^, large vs. small hepatocytes^81^. We curated 3 sets of diploidy signatures, including DEGs for pre-PH vs. PH^81^, control vs. Cdk1 knockout^81^, and 2n vs. 4n hepatocytes^82^. We curated signature gene sets for pericentral (CV) and periportal (PN) hepatocytes from Halpern et al.^83^. We curated gene sets for putative zonated pathways from MSigDB^122, 123^ (v7.1), including glycolysis (pericentral), bile acid production (pericentral), lipogenesis (pericentral), xenobiotic metabolism (pericentral), beta-oxidation (periportal), cholesterol biosynthesis (periportal), protein secretion (periportal), and gluconeogenesis (periportal). All signature gene sets are reported in Supp. Table 10. For the joint regression analysis of `scDRS` disease score on ploidy and zonation scores, we regressed the polyploidy score out of both the pericentral and periportal score before the joint regression because the ploidy level confounded both zonation scores. We performed joint regression instead of marginal regression here (unlike the regression analysis in the neuron section) because the polyploidy score was positively correlated with the pericentral and periportal scores (unlike the analysis in the neuron section where the 3 sets of scores had low correlations).

## Data availability

We release our data at https://figshare.com/projects/Single-cell_Disease_Relevance_Score_scDRS_/118902 (instructions at https://github.com/martinjzhang/scDRS), including GWAS summary statistics of the 74 diseases/traits, TMS FACS scRNA-seq data, reprocessed TMS FACS data (for T cells and hepatocytes), MAGMA and gold standard gene sets, and `scDRS` results for TMS FACS (disease scores and control scores for the 74 diseases/traits). The 16 scRNA-seq data sets were obtained as follows. The TMS FACS data and TMS droplet data^19^ was downloaded from the official release https://figshare.com/articles/dataset/Processed_files_to_use_with_scanpy_/8273102. The TS FACS data^25^ was downloaded from the official release https://figshare.com/articles/dataset/Tabula_Sapiens_release_1_0/14267219. The Cano-Gamez & Soskic et al. data^51^ was downloaded from https://www.opentargets.org/projects/effectorness. The Nathan et al. data^64^ was downloaded from https://www.ncbi.nlm.nih.gov/geo/query/acc.cgi?acc=GSE158769. The Zeisel & Muñoz-Manchado et al. data^65^ was downloaded from http://linnarssonlab.org/cortex/. The Zeisel et al. data^73^ was downloaded from http://mousebrain.org/downloads.html. The Habib & Li et al. data^69^ and Habib, Avraham-Davidi, & Basu et al. data^75^ were downloaded from https://singlecell.broadinstitute.org/single_cell. The Ayhan et al. data^70^ was downloaded from https://cells.ucsc.edu/human-hippo-axis/. The Yao et al. data^74^ was downloaded from https://assets.nemoarchive.org/dat-jb2f34y. The Zhong et al. data^76^ was downloaded from https://www.ncbi.nlm.nih.gov/geo/query/acc.cgi?acc=GSE119212. The Aizarani et al. data^86^ was downloaded from https://www.ncbi.nlm.nih.gov/geo/query/acc.cgi?acc=GSE124395. Halpern & Shenhav et al. data^83^ was downloaded from https://www.ncbi.nlm.nih.gov/geo/query/acc.cgi?acc=GSE84498. The Richter & Deligiannis et al. data^82^ (annotated count matrix) was obtained via communication with the authors (raw data publicly available via links in the paper). The Taychameekiatchai et al. data^85^ is not publicly available, but was obtained via communication with the authors.

## Code availability

Software implementing `scDRS` and its downstream applications and a web interface for interactively exploring results of `scDRS` are available at https://github.com/martinjzhang/scDRS.

## Acknowledgements

We thank Huwenbo Shi, Katherine Siewert-Rocks, Tiffany Amariuta, Xiaoyu Xu, Benjamin J Strober, and Alexander Gusev for helpful suggestions. This research was funded by NIH grants U01 HG009379, R01 MH101244, R37 MH107649, R01 MH115676, and U01 HG012009.

