## Supplementary Info for "Polygenic enrichment distinguishes disease associations of individual cells in single-cell RNA-seq data"

### scDRS

Zhang & Hou et al.

### Supplementary Information

### Supplementary Note

#### Estimation of gene-specific technical noise variance of gene expression

We estimate the technical variance similar to previous works[1,2]. Specifically, we first compute the mean expression and expression variance for each gene across all cells in the data set in the original non-log-transformed space. Next, we fit a non-linear tread to the log10-scale variance/mean relationship using local polynomial regression (span=0.3, degree=2). The estimated trend models the expected technical variance based on the mean expression; observed variance values above this expected trend reflect biological variance. Given this trend, the proportion of technical variance is computed as the ratio of predicted technical variance to observed variance (in the original non-log-transformed space). Note that it is possible for this ratio to be greater than 1 if the observed variance is less than the expected variance. Finally, the technical variance of the log-transformed data $\sigma^2_{\text{tech},g}$ is computed as the product of the variance of the gene (computed using the log-transformed data) and the proportion of technical variance as estimated above.

#### Distribution of raw disease scores and raw control scores

Here we characterize the distribution of the raw disease scores and raw control scores under a simplified model. Assume the expression levels $\{\mathbf{X}_c \in \mathbb{R}^{n_{\text{gene}}}\}_{c=1}^{n_{\text{cell}}}$ are independently and identically distributed (i.i.d.) across cells with mean $\boldsymbol{\mu} \in \mathbb{R}^{n_{\text{gene}}}$ and covariance $\Sigma \in \mathbb{R}^{n_{\text{gene}} \times n_{\text{gene}}}$,

$$\mathbf{X}_c \overset{\text{i.i.d.}}{\sim} (\boldsymbol{\mu}, \Sigma). \tag{1}$$

Without loss of generality, also assume the gene weights are the same. Then the raw disease score and raw control scores (for a given control gene set $G_b^{\text{ctrl}}$) can be written as

$$s_c = \frac{1}{|G|}\sum_{g\in G} X_{cg}, \quad s_{cb}^{\text{ctrl}} = \frac{1}{|G_b^{\text{ctrl}}|}\sum_{g\in G_b^{\text{ctrl}}} X_{cg}, \tag{2}$$

where $|G| = |G_b^{\text{ctrl}}|$. We next compare the distributions of $s_c$ and $s_{cb}^{\text{ctrl}}$. Since the control genes match the mean expression and expression variance of the disease genes, we have

$$\sum_{g\in G}\mu_g = \sum_{g\in G_b^{\text{ctrl}}}\mu_g, \quad \sum_{g\in G}\Sigma_{gg} = \sum_{g\in G_b^{\text{ctrl}}}\Sigma_{gg}. \tag{3}$$

The first equation gives that $\mathbb{E}[s_c] = \mathbb{E}[s_{cb}^{\text{ctrl}}]$.

For covariance, since the disease genes are more positively correlated due to co-expression in the associated cell population, the covariance matrix of the disease genes has larger off-diagonal elements than that of the control genes. Since the sum of the diagonal elements are equal between the two matrices due to the second part of Eq. (3), we have

$$\sum_{g,g'\in G}\Sigma_{g,g'} > \sum_{g,g'\in G_b^{\text{ctrl}}}\Sigma_{g,g'} \tag{4}$$

Since the variances of the raw disease score and raw control score can be written as

$$\text{Var}[s_c] = \frac{1}{|G|^2}\sum_{g,g'\in G}\Sigma_{g,g'}, \quad \text{Var}[s_{cb}^{\text{ctrl}}] = \frac{1}{|G_b^{\text{ctrl}}|^2}\sum_{g,g'\in G_b^{\text{ctrl}}}\Sigma_{g,g'}, \tag{5}$$

we have $\text{Var}[s_c] > \text{Var}[s_{cb}^{\text{ctrl}}]$. In summary,

$$\mathbb{E}[s_c] = \mathbb{E}[s_{cb}^{\text{ctrl}}], \quad \text{Var}[s_c] > \text{Var}[s_{cb}^{\text{ctrl}}]. \tag{6}$$

Since the raw disease scores have the same mean and a higher variance than the raw control scores, the top values of the raw disease scores are larger than all raw control scores. These top values correspond to the disease-associated cells, which have higher expression of the disease genes. Therefore, the disease-associated cells have larger raw disease scores than all raw control scores.

We are primarily interested in the case where disease genes have correlated expression patterns due to co-expression in a disease-relevant cell population (first case). However, a caveat is that disease genes may instead have correlated expression patterns due to a subset of disease genes being physically proximal in the genome (second case; this is not the case that `scDRS` aims to detect). The major difference between the two cases is that in the first case, disease genes have broadly correlated expression patterns (due to co-expression in the relevant cell population), while in the second case, the disease genes are expected to have correlated expression patterns only in small groups of genes (that lie in the same genomic region). `scDRS` does not rigorously control for this second case because of the difficulty of separating the two cases across a continuous spectrum of levels of correlations. However, the level of correlations due to the second case is expected to be lower and such correlations are not expected to concentrate in specific cell populations. So `scDRS` is expected to produce far fewer significant associations, if any, in the second case where the signals are weaker and do not concentrate in specific cell populations. Indeed, we performed additional null simulations using genomic location-matched null gene sets and determined that `scDRS` was well-calibrated (Supp. Fig. 5).

#### Normalization of disease scores and control scores

**First gene set alignment.** This step aims to correct for the mismatch that the control genes do not have exactly the same mean expression and expression variance as the disease genes. The variance level is estimated based on a heuristic that assumes the genes in the given gene set are independent. Specifically, if $Y = \sum_{i=1}^{n} w_i X_i$ for weights $w_1, \cdots, w_n \in \mathbb{R}$ and independent random variables $X_1, \cdots, X_n$, then $\text{Var}[Y] = \sum_{i=1}^{n} w_i^2 \text{Var}[X_i]$.

**Cell-wise standardization.** Consider the conditional distribution of the raw control score $s_{cb}^{\text{ctrl}}$ given the expression matrix $\mathbf{X}$, where the randomness only comes from MC sampling of the control gene set $G_b^{\text{ctrl}}$ but not from the data generation process of $\mathbf{X}$. Again without loss of generality, assume all genes have the same mean and variance, and the gene weights are the same. Then the raw control score can be written as

$$s_{cb}^{\text{ctrl}} = \frac{1}{|G_b^{\text{ctrl}}|} \sum_{g \in G_b^{\text{ctrl}}} X_{cg}. \tag{7}$$

Since all genes have the same mean and variance, $G_b^{\text{ctrl}}$ corresponds to randomly and uniformly sampling $|G|$ genes from the set of all genes $\{1, \cdots, n_{\text{gene}}\}$ without replacement. When $|G| \ll n_{\text{gene}}$, sampling without replacement can be well approximated by sampling with replacement ($|G| < 20\% n_{\text{gene}}$ is usually a good heurestic[3]). In other words,

$$s_{cb}^{\text{ctrl}} \overset{d}{\approx} \frac{1}{|G|} \sum_{g=1}^{|G|} Y_{cg}, \tag{8}$$

where $Y_{cg}$'s are i.i.d. random variables uniformly sampled from $\{X_{c1}, \cdots, X_{cn_{\text{gene}}}\}$ (meaning sampling with replacement) and $\overset{d}{\approx}$ means approximately equal in distribution. Furthermore, when $|G|$ is large (e.g., >50), by the central limit theorem, being an average of $|G|$ i.i.d. random variables, $s_{cb}^{\text{ctrl}}$ is close to a normal distribution, which depends only on its first two moments. Therefore, it is sufficient to only match the mean and variance of the control score distributions of different cells.

As a remark, the raw control score distribution considered here is defined with respect to randomly sampling control gene sets for a given cell, where the expression matrix $\mathbf{X}$ is fixed. It is different from the distribution of scores computed from a given gene set $S$ (disease gene set or control gene set) across cells, where the randomness can be viewed as coming from the data generation process of the expression matrix $\mathbf{X}$. In this latter case, since the gene expression levels are correlated, the score of a given cell $s = \frac{1}{|S|} \sum_{g \in S} X_{cg}$ can not be viewed as an average of i.i.d. random variables (because $X_{cg}$'s may be dependant) and can be very different from the normal distribution especially when the gene set size $|S|$ is small.

**Second gene set alignment.** This step aims to correct for the differences of the mean values of scores from different gene sets introduced by cell-wise standardization. The differences are relatively small (Supp. Fig. 1F). Hence, this step is less important. For this reason and also because it is hard to find a good heuristic for estimating the variance levels of scores from different gene sets without downweighting the disease scores due to the higher correlation between disease genes, we do not correct for the difference of the variance of scores from different gene sets.

#### Secondary analyses for simulations assessing calibration and power

We performed 4 secondary analyses pertaining to null simulations. First, we considered other numbers of putative disease genes (100, 500, or 2,000, instead of 1,000). We determined that `scDRS` remained well-calibrated, VAM continued to suffer from severely inflated type I error, and Seurat and Vision suffered increased type I error at 500 genes and severely inflated type I error at 100 genes (Supp. Fig. 4A-D). Second, we considered biased sets of putative disease genes (randomly selected from genes with high mean expression, genes with high expression variance, or overdispersed genes (genes with high expression variance but normal levels of technical noise[4])). We determined that `scDRS` remained reasonably well-calibrated, VAM continued to suffer from severely inflated type I error, and Seurat and Vision were conservative for high-expression genes and high-variance genes but suffered inflated type I error for overdispersed genes (Supp. Fig. 4E-P). Third, we assessed calibration of our MC test for *cell type*-disease association based on the output of scDRS, using the same subsampled data (and 1,000 putative disease genes). We confirmed that this test was well-calibrated (Supp. Table 12). Fourth, we considered null gene sets matching the genomic locations of real GWAS putative disease gene sets, and determined that scDRS was well-calibrated in this simulation (Supp. Fig. 5).

We performed 4 secondary analyses pertaining to causal simulations. First, we considered other levels of overlap between the 1,000 causal genes and 1,000 putative disease genes (from 5% to 50%, instead of 25%). We determined that scDRS continued to attain higher power than Seurat and Vision (Supp. Fig. 6B). Second, we considered causal simulations in which we selected all 528 B cells in the subsampled data as causal cells, instead of randomly selecting 500 causal cells. We observed a similar improvement in power of scDRS over Seurat and Vision (Supp. Fig. 6C). Third, we computed the actual FDR for each method in each of the above causal simulations. We determined that scDRS attained well-calibrated FDR across all parameter settings, whereas Seurat and Vision suffered from inflated type I error at smaller effect sizes ($\leq$1.1 times higher expression for causal cells) and lower levels of overlap ($\leq$15%) (Supp. Fig. 6D-F). Fourth, since Seurat and Vision were not initially designed to produce calibrated p-values, we also evaluated each method's area under the receiver operating characteristic curve (AUC) in distinguishing causal from non-causal cells. We determined that scDRS attained more accurate classification than Seurat and Vision under this metric (Supp. Fig. 6G-I).

#### Secondary analyses for results across 120 TMS cell types for 74 diseases and complex traits

We discuss the 2 exceptions to the block-diagonal pattern, involving 4 diseases/traits (Fig. 3). First, ventricular myocytes (in addition to immune cell types) were associated with lymphocyte count. This association is consistent with the prognostic value of relative lymphocyte concentration in patients with symptomatic heart failure[5], but to our knowledge has not been reported in previous genetic studies. Second, pancreatic beta cells (in addition to brain cell types) were associated with SCZ, body mass index (BMI), and smoking status; risk variants for SCZ and BMI are reported to be enriched in pancreatic islet-specific epigenomic regulatory elements[6,7].

We performed 4 secondary analyses to assess robustness of these results. First, we performed the same analyses on a mouse cell atlas assayed with a different technology (TMS droplet) and a human cell atlas assayed using the same technology (TS FACS) to provide comparisons of the results across technologies and across species. Results are reported in Supp. Fig. 10. We determined that the associations are highly consistent across technologies ($r$ =0.91 for association $-\log_{10}$ p-value across cell type-disease pairs; $P$ =$2.8\times10^{-24}$, Fisher's exact test) and reasonably consistent across species ($r$ =0.63 for association $-\log_{10}$ p-value; $P$ =$1.3\times10^{-7}$, Fisher's exact test). Second, we analyzed the same 120 TMS FACS cell types and 74 diseases using LDSC-SEG[8] and 3 methods in Bryois et al.[9] (LDSC-specificity, MAGMA-specificity, and the combined method) for comparison purposes (Methods). Results are reported in Supp. Fig. 11. We determined that the cell type-disease associations identified by scDRS are highly consistent with these 4 methods ($r$ =0.67, 0.69, 0.64, 0.7 for association $-\log_{10}$ p-value between scDRS and the 4 comparison methods respectively; $P < 1\times10^{-300}$ for all 4 comparisons, Fisher's exact test). Third, since scDRS computes gene-level statistics across all cells in the data set, the results may be biased towards major cell types with many cells. We assessed the impact of this potential bias by implementing a new version of scDRS that adjusts for cell type proportions so that the results will only depend on the set of cell types in the data set (but not the number of cells of each cell type), analogous to other disease-cell type association methods[8–10]; we determined this version of scDRS was highly consistent with the default version on TMS FACS (median scDRS disease score correlation of 0.97 across 74 diseases/traits) and was well-calibrated in the null simulations (Supp. Fig. 4; Methods). Fourth, we determined that results are consistent between using different scaling factors for the size-factor normalization of the single-cell data (median scDRS disease score correlation of 0.90 across 74 diseases/traits between scaling to the default 10,000 vs. 1 million reads per cell).

We performed 2 secondary analyses to assess alternative versions of scDRS. First, we investigated other methods for selecting putative disease genes (top 100, top 500, top 2,000, FWER<5%, or FDR<1%, instead of top 1,000), other gene weights (no weights, GWAS z-score weights, or single-cell VS weights, instead of using both sets of weights), and other MAGMA window sizes for mapping SNPs to genes (0 kb or 50 kb, instead of 10 kb). We evaluated the performance based on a curated set of 20 traits with expected and unexpected disease-critical cell types (Supp. Table 17; Methods). We determined that our default version of scDRS (top 1,000 genes, GWAS z-score + single-cell VS weights, 10-kb MAGMA window) significantly outperformed all other methods (except the method using the top 2,000 genes, which was not significantly better than the default), but was highly consistent with other versions (median scDRS score correlations 0.49-0.98; Supp. Fig. 12,13, Supp. Table 18). Second, we considered an overdispersion score capturing both overexpression and underexpression of putative disease genes in the relevant cell population (whereas the default weighted score only captures overexpression; Methods). We determined our default weighted score achieved substantially higher power than the overdispersion score, suggesting that most putative disease genes are overexpressed in the relevant cell population (Supp. Fig. 14).

#### Comparison with other cell type-level association methods

We center our discussion around 3 cell type-level association methods that also make use of MAGMA: the MAGMA-based method in Skene et al.[11], Watanabe et al.[10], and the MAGMA-based method in Bryois et al.[9] All statements about scDRS apply to both individual cell level-analysis and cell type-level analysis.

We first discuss the points previously raised in these 3 works. First, scDRS assesses the excess expression of putative disease genes in a given cell relative to control genes with similar mean expression levels across all cells in the data set. So it is in line with the other 3 methods that focus on specifically-expressed genes in a cell type relative to other cell types rather than just highly-expressed genes in the given cell type. Second, the results of the other 3 methods depend on the cell types in the data set while the scDRS result depends not only on the cell types but also individual cells in the data set (therefore the scDRS result also depends on cell type proportions). The additional dependency on cell type proportions is not a major issue for relatively balanced data sets such as TMS FACS (as shown in the Results across 120 TMS cell types ... subsection in the Results section). Also, we provide an option to explicitly adjust for cell type proportions (Discussion). Third, a concern raised by Bryois et al. about Watanabe et al. is that the latter method is sensitive to different scaling factors for size factor normalization. This concern applies to scDRS but not the methods in Skene et al. and Bryois et al. We choose to retain these preprocessing steps (size factor normalization and log transformation) because they are essential for correcting for the confounding effect of sequencing depth and stabilizing the count data and are recommended best practices[1,12]. In addition, we have determined that the results of scDRS are consistent between using different scaling factors (Results). Fourth, a concern raised by Watanabe et al. about Skene et al. is that the latter method does not condition on the average expression level of a gene across cell types when regressing the MAGMA z-scores against cell type specificity across genes, and is thus "vulnerable to confounding by a general effect of gene expression"[10]. This concern also applies to Bryois et al. scDRS is not susceptible to this concern, because a general effect of gene expression on trait association would be present in both the putative disease genes and the matched control genes (which have the same mean expression), and would thus cancel each other out.

Overall, the other 3 methods use linear regression to associate MAGMA z-scores with cell type features across genes (cell type expression level in Watanabe et al.; cell type specificity in Skene et al. and Bryois et al.). scDRS can be viewed as a non-parametric alternative to these methods, employing a stratified permutation test that associates MAGMA z-scores for top genes with expression levels for a given cell by permuting genes within levels of expression mean and variance. Thus, unlike the other 3 methods, scDRS does not rely on a linearity assumption and does not need to correct for gene-level technical covariates (such as gene length), because these covariates are expected to affect gene expression levels uniformly across cells and are unlikely to have additional cell-specific effects after conditioning on the expression mean and variance across cells.

scDRS has 2 potential advantages over the other 3 methods in terms of cell type-level analysis. First, scDRS does not assume a linear relationship between MAGMA z-scores and cell type features across genes (see above). Second, scDRS may be more robust to within-cell type heterogeneity. For example, if only 20% of cells in a heterogeneous cell type are disease-associated, the cell type features in the other 3 methods will be dominated by the non-associated 80% of cells and may not be able to capture the associated subpopulation. In comparison, scDRS tests if the top 5% most associated cells in the cell type are significantly associated with the disease, and may therefore be able to capture the associated subpopulation.

#### Secondary analyses for heterogeneous subpopulations of T cells associated with autoimmune disease

We performed 4 secondary analyses. First, we assessed cell type-disease associations using scDRS for two human scRNA-seq data sets (Cano-Gamez & Soskic et al.[13] and Nathan et al.[14]; Supp. Table 2) and each of the 10 autoimmune diseases (and height, a negative control trait); we focused on cell type-disease associations because these data sets contain well-annotated T cell subtypes and states. Results are reported in Supp. Table 23. In the Cano-Gamez & Soskic et al. data, natural Tregs and cytokine-induced Th17/Treg cells were significantly associated with IBD (FDR<0.05, MC test). In the Nathan et al. data, *RORC*$^+$ Tregs, Th17/Th1 cells, CD161$^+$ Th2 cells, and activated CD4$^+$ T cells were weakly associated with IBD (FDR<0.2, MC test). These findings are consistent with our discoveries in TMS FACS linking effector T cells, particularly Tregs and Th17 cells, to IBD. In addition, as a negative control, no cell type was significantly associated with height in these two data sets. Second, we compared scDRS to cluster-level analyses using LDSC-SEG at various clustering resolutions. Results are reported in Supp. Fig. 24. We determined that both methods produced similar results at the cluster level, but the cluster-level analyses failed to recapitulate the individual cell-disease associations detected in the scDRS individual cell-level analysis (even when clustering at a very high resolution). Third, we investigated alternative disease gene prioritization methods, including prioritizing genes based on specific expression in the disease-critical T cell population (differentially expressed genes for comparing T cells vs. other cells in the TMS FACS data) and based on correlating the expression level of a given gene with scDRS disease scores across T cells, CD4$^+$ T cells, or CD8$^+$ T cells (instead of all TMS FACS cells). We determined that our primary approach provided a more accurate prioritization of gold-standard disease-relevant genes (Supp. Fig. 25A-J). Fourth, we extended our prioritization of disease-relevant genes to all 74 diseases/traits. We compared the prioritized genes with drug target genes for 27 diseases and Mendelian disease genes for 45 diseases (Supp. Table 21). We determined that our approach attained a similar improvement over MAGMA across this broader set of diseases/traits (Supp. Fig. 25K-N).

#### Secondary analyses for heterogeneous subpopulations of hepatocytes associated with metabolic traits

For inferred polyploidy scores, we verified that the inferred high-ploidy hepatocytes had higher expression levels of the *Xist* (X-inactive specific transcript) non-coding RNA gene (for hepatocytes in female mice) and higher numbers of expressed genes (Supp. Fig. 32H,I), two distinguishing features of high-ploidy hepatocytes[15,16]. We further verified that the inferred polyploidy score obtained by applying this procedure to independent data[16] with experimentally determined ploidy level annotation were significantly correlated with the experimentally determined annotation ($r$ =0.28, $P$ <0.001, MC test), and that the inferred zonation scores obtained by applying this procedure to independent data[17] with experimentally determined zonation annotations were significantly correlated with the experimentally determined annotations ($r$ =0.42, $P$ <0.001 for pericentral score, $r$ =0.45, $P$ <0.001 for periportal score, MC test).

We performed 4 secondary analyses. First, we reapplied `scDRS` to 4 additional mouse single-cell data sets[16–19] and 1 human single-cell data set[20] (Supp. Table 2). Results are reported in Supp. Fig. 34A,B. The results suggest consistent association of the polyploidy score and both the pericentral and periportal scores with the 9 metabolic traits. Second, given that `scDRS` associated both pericentral and periportal hepatocytes to metabolic traits, we assessed whether `scDRS` is able to detect pericentral-specific and periportal-specific effects. We analyzed all 6 hepatocyte scRNA-seq data sets using 8 metabolic pathway gene sets[21,22] (instead of MAGMA genes from GWAS; Supp. Table 10; Methods) whose zonation patterns are well-understood (4 pericentral-specific pathways and 4 periportal-specific pathways[23]). Results are reported in Supp. Fig. 34C,D. We determined that pericentral-specific pathways generally exhibited pericentral-specific effects, and periportal-specific pathways generally exhibited periportal-specific effects. Third, we assessed the robustness of our polyploidy score by inferring the ploidy level of hepatocytes using 3 additional sets of polyploidy signatures and 3 additional sets of diploidy signatures[24] (expected to be negatively correlated with the polyploidy score; Methods) for each of the 6 data sets. Results are reported in Supp. Table 27. We determined that the polyploidy score is strongly positively correlated with scores obtained using the additional polyploidy signatures ($P$ <0.005 for 17/18 correlations, MC test) and strongly negatively correlated with scores obtained using the additional diploidy signatures ($P$ <0.005 for 10/18 correlations, MC test). Fourth, the association between hepatocyte ploidy level and metabolic traits may imply that there are metabolic trait GWAS variants associated with ploidy (ploidyQTL). We observed significant enrichment for the GWAS gene sets for these metabolic traits for genes from the polyploidy signature gene set[24] (average odds ratio 1.46 with SE=0.09 across the 9 metabolic traits), suggesting that GWAS SNPs for these metabolic traits may be enriched for ploidyQTL. We were unable to assess this directly because ploidyQTL data are limited, as genetic studies of ploidy level have largely focused on organisms other than humans[25], perhaps because most human cells are diploid.

#### Related works

**Previous methods for identifying disease-critical tissues and cell types.** Many types of data that assay gene regulation have been integrated with GWAS data to identify disease-relevant tissues and cell types, including chromatin and histone modifications[26–35] and gene expression measurements[8–10,36–42]. Studies using gene expression data have generally either used tissue-level data derived from DNA microarrays / bulk RNA-Seq[8,36–39], or focused on predefined cell types (usually classical cell types based on known marker genes) in scRNA-seq data by aggregating cells from the same cell type[9,10,40]; these cell type annotations may be hard to obtain especially for less well-studied cell populations or subtle cell states within a cell type. One exception is Jagadeesh and Dey et al.[42], who associated intra- and inter-cell type cellular processes identified in scRNA-seq data to disease.

**Previous methods for scoring individual cells.** Previous works analyzing scRNA-seq data alone have used individual cell-level scores to characterize cellular heterogeneity and subtle cell states within classically defined cell types[2,43–48], where the cell scores were computed based on the expression of a predefined set of genes such as cell cycle signature genes or biological pathways. These works did not integrate GWAS data. In addition, they generally did not provide individual cell-level p-values for associating individual cells to the gene set (VAM[2] is the only exception, but we show that VAM suffers severely inflated type I error; Fig. 2A). We further note that two studies have associated individual cells in scATAC-seq data to disease[34,35]. However, scATAC-seq and scRNA-seq data have different data structures and require different treatments (e.g., matching nucleotide GC content and fragment accessibility for scATAC-seq data[34] vs. matching gene expression mean and variance in our paper). To our knowledge, no previous study has associated individual cells in scRNA-seq data to disease.

#### Additional limitations of `scDRS`

We note additional limitations and future directions of our work. First, the relevant cell-level variables that we identified (e.g., T cell effectorness gradients for autoimmune diseases) only partially explain the heterogeneity across individual cells in their association to disease; there are likely more cell-level variables driving this heterogeneity that remain to be identified.

Second, we primarily used mouse RNA-seq data (TMS FACS) to study human diseases and complex traits, but there are biological differences between human and mouse. Arguments in favor of using mouse RNA-seq data to study human diseases and complex traits include (1) it is easier to obtain high-quality atlas-level scRNA-seq data from mice, (2) our key findings were replicated in human data, (3) we evaluated only protein-coding genes with 1:1 orthologs between mice and humans, which are highly conserved, (4) we used a large number of genes to associate cells to diseases (1,000 MAGMA putative disease genes), minimizing potential bias due to individual genes differentially expressed across species (see Bryois et al.[9] and other studies[8,10,11,40] for additional discussion). However, it is possible that some cell types are less conserved across species[9,49] (e.g., our results for CA1 pyramidal neurons along the long and radial axes (Supp. Fig. 30) seem to indicate different disease association patterns between human and mouse), motivating follow-up analyses involving human scRNA-seq data (including those that we have performed here). Third, `scDRS` detects overexpression of putative disease genes (analogous to previous works[8–10]), but is not designed to detect underexpression. Our initial implementation of an overdispersion score was less well-powered than `scDRS` in analyses of real disease/traits (Supp. Fig. 14), but further efforts to combine directional and overdispersion scores may be warranted[50]. Fourth, `scDRS` results for a given cell depends on the other cells in the data set through both the estimation of technical noise levels and the selection of matched control genes; however, both steps depend only on gene-specific quantities averaged across all cells (gene-specific expression mean and expression variance) and are thus robust to inclusion or exclusion of a small set of cells (or a large random subset of cells). Fifth, the fact that `scDRS` assesses the statistical significance of an individual cell's association to disease by implicitly comparing it to other cells via matched control genes may reduce power if most cells in the data are truly causal. For example, association with IBD in a data set containing only Tregs (one of the causal cell types for IBD) will likely yield largely non-significant results. This limitation did not impact our main analyses, because the TMS data includes a broad set of cell types; in more specialized data sets (which may be preferred in some settings due to the more comprehensive profiling of the focal cell population), this limitation can potentially be addressed by selecting matched control genes based on a broad cell atlas (e.g., the TMS or TS data). Sixth, we have only analyzed scRNA-seq data from control samples. Extending `scDRS` to analyze scRNA-seq data from case-control samples or experimentally perturbed samples[51], perhaps by applying `scDRS` and comparing disease scores of cells from different conditions, may provide further insights about disease.

64. Fatma Ayhan, Ashwinikumar Kulkarni, Stefano Berto, Karthigayini Sivaprakasam, Connor Douglas, Bradley C Lega, and Genevieve Konopka. Resolving cellular and molecular diversity along the hippocampal anterior-to-posterior axis in humans. *Neuron*, 2021.

**78.** Elena Gianchecchi and Alessandra Fierabracci. Inhibitory receptors and pathways of lymphocytes: the role of pd-1 in treg development and their involvement in autoimmunity onset and cancer progression. *Frontiers in immunology*, page 2374, 2018.

### Supplementary Tables

See Supplementary Excel file

**Supplementary Table 1. GWAS diseases and complex trait data sets.** We report the name, identifier, code, category, reference, sample size, number of variants, estimated heritability using LD Score regression (LDSC)[52–54], and z-score for non-zero heritability for the 74 diseases/traits analyzed in the paper. We also report polygenicity for a subset of 21 diseases/traits ($\log_{10} M_e$ for common SNPs; Table 1 in O'Connor et al.[55]). The disease called "Auto Immune Traits" (in UK Biobank) is based on the following codes and disease names that characterize autoimmune physiopathogenic etiology: 1222 (t1d; type 1 diabetes); 1256 (guillainBarre); 1260 (myasthenia); 1261 (ms); 1372 (vasculitis); 1378 (granulomatosis with polyangiitis, previously known as Wegener's granulomatosis); 1381 (sle); 1382 (Sjogren); 1384 (sysSclerosis); 1437 (myasthenia); 1456 (celiac); 1464 (ra); 1522 (grave); 1661 (vitiligo)[56,57]. Note that myasthenia gravis appears twice (under codes 1260 and 1437)[58] and both codes were used.

| **Data set** | **Species** | $N_{\text{cell}}$ | $N_{\text{tissue}}$ | $N_{\text{cell type}}$ | **Description** |
|---|---|---|---|---|---|
| TMS FACS[18] | Mus musculus | 110,096 | 23 | 120 | Mouse cell atlas (FACS + Smart-seq2) |
| TMS droplet[18] | Mus musculus | 245,389 | 16 | 123 | Mouse cell atlas (10x microfluidic droplets) |
| TS FACS[59] | Homo sapiens | 26,813 | 24 | 134 | Human cell atlas (FACS + Smart-seq2) |
| Cano-Gamez & Soskic et al.[13] | Homo sapiens | 43,112 | 1 | 22 | Subtypes of naive, memory, and activated $CD4^{+}$ T cells from the blood |
| Nathan et al.[14] | Homo sapiens | 500,089 | 1 | 29 | T cells from the blood |
| Zeisel & Muñoz-Manchado et al.[60] | Mus musculus | 3,005 | 1 | 9 | Cortex and hippocampus; 827 CA1 pyramidal cells |
| Zeisel et al.[61] | Mus musculus | 160,797 | 1 | 265 | Whole nervous system; 304 CA1 pyramidal cells |
| Habib & Li et al.[62] | Mus musculus | 1,367 | 1 | 7 | Hippocampal regions from adult mice; 155 CA1 pyramidal cells; snRNA-seq |
| Habib, Avraham-Davidi, & Basu et al.[63] | Homo sapiens | 14,963 | 1 | 21 | Archived brain sample; 421 CA1 pyramidal cells; snRNA-seq |
| Ayhan et al.[64] | Homo sapiens | 129,908 | 1 | 24 | Surgically resected anterior and posterior hippocampus from epilepsy patients; 5,454 CA1 pyramidal cells; snRNA-seq |
| Yao et al.[65] | Mus musculus | 74,974 | 1 | 388 | Cortex and hippocampus; 1,701 CA1 pyramidal cells using SMART-Seq v4 technology |
| Zhong et al.[66] | Homo sapiens | 30,416 | 1 | 11 | Hippocampus at gestational weeks 16–27; 5,972 CA1 pyramidal cells |
| Aizarani et al.[20] | Homo sapiens | 10,372 | 1 | 11 | Hepatocytes, endothelial cells, and other common cell types from the liver |
| Halpern & Shenhav et al.[17] | Mus musculus | 1,415 | 1 | 1 | Hepatocytes |
| Richter & Deligiannis et al.[16] | Mus musculus | 1,649 | 1 | 1 | Sorted 2n and 4n hepatocytes (Hoechst dye + FACS); snRNA-seq |
| Taychameekiatchai et al.[19] | Mus musculus | 19,254 | 1 | 15 | Hepatocytes, endothelial cells, and other common cell types from the liver |

**Supplementary Table 2. ScRNA-seq and snRNA-seq data sets.** We report the reference, species, number of cells, number of tissues, number of cell types, and a short description for each scRNA-seq/snRNA-seq data set analyzed in the paper. Data sets without "snRNA-seq" in the description are scRNA-seq data sets. The 16 data sets contain more than 1.3 million cells from 31 tissues and organs, including aorta, brown adipose tissue (BAT), bladder, blood, bone marrow, brain myeloid, brain non-myeloid, diaphragm, eye, gonadal adipose tissue (GAT), heart, kidney, large intestine, limb muscle, liver, lung, lymph node, mesenteric adipose tissue (MAT), mammary gland, pancreas, prostate, subcutaneous adipose tissue (SCAT), salivary gland, skin, small intestine, spleen, thymus, tongue, trachea, uterus, vasculature. For clarification, Zeisel & Muñoz-Manchado et al. refers to the data from Zeisel & Muñoz-Manchado et al. 2015 *Science*[60] and Zeisel et al. refers to the data from Zeisel et al. 2018 *Cell*[61].

See Supplementary Excel file

**Supplementary Table 3. Gene-level statistics.** We report mean expression, expression variance, technical variance, and proportion of zero counts for each gene of the normalized log-scale data for the 16 data sets.

See Supplementary Excel file

**Supplementary Table 4. Correlation of technical variance between data sets.** We report the correlation of technical variance, computed across genes, for each pair of the 16 data sets.

See Supplementary Excel file

**Supplementary Table 5. Cell types in the TMS FACS data.** We report 120 cell types in the TMS FACS data, the corresponding number of cells, and the corresponding tissue composition (for tissues consisting $>1\%$ of cells from the cell type).

See Supplementary Excel file

**Supplementary Table 6. Cell types in the TMS droplet data.** We report 123 cell types in the TMS droplet data, the corresponding number of cells, and the corresponding tissue composition (for tissues consisting $>1\%$ of cells from the cell type).

See Supplementary Excel file

**Supplementary Table 7. Cell types in the TS FACS data.** We report 132 cell types in the TS FACS data, the corresponding number of cells, and the corresponding tissue composition (for tissues consisting $>1\%$ of cells from the cell type).

See Supplementary Excel file

**Supplementary Table 8. MAGMA gene sets.** We report MAGMA gene sets and corresponding GWAS MAGMA z-score gene weights for the 74 diseases and traits.

See Supplementary Excel file

**Supplementary Table 9. MSigDB terms for curating signature gene sets.** We report the MSigDB terms used to curate the signature gene sets in the paper.

See Supplementary Excel file

**Supplementary Table 10. Signature gene sets.** We report the signature gene sets used in the paper.

See Supplementary Excel file

**Supplementary Table 11. Numerical results for null simulations in Fig. 2A.** We report the mean and SE of p-value quantiles for different cell-scoring methods over 100 repetitions.

| Nominal FDR level | 0.05 | 0.1 | 0.2 |
|---|---|---|---|
| Actual FDR level | 0.00±0.00 | 0.01±0.02 | 0.14±0.07 |

**Supplementary Table 12. Results for null simulations for testing cell type-disease association.** We assessed calibration of the MC test for cell type-disease association based on the output of `scDRS`. We used the same subsampled data (10,000 cells from TMS FACS) and 1,000 randomly-selected disease genes. We report the actual FDR for multiple testing across all 118 cell types in the subsampled data at various nominal FDR levels. 95% confidence intervals were provided based on the 100 repetitions.

See Supplementary Excel file

**Supplementary Table 13. Numerical results for causal simulations in Fig. 2B.** We report the mean and SE of power at various effect sizes for different cell-scoring methods over 100 repetitions.

See Supplementary Excel file

**Supplementary Table 14. Numerical results for cell type-level analyses for the TMS FACS data, including results in Fig. 3.** For each pair of cell type and disease/trait, we report the proportion of significantly associated cells (FDR<0.1), FDR for cell type-disease association, and FDR for within-cell type disease association heterogeneity.

See Supplementary Excel file

**Supplementary Table 15. Numerical results for cell type-level analyses for the TMS droplet data.** For each pair of cell type and disease/trait, we report the proportion of significantly associated cells (FDR<0.1), FDR for cell type-disease association, and FDR for within-cell type disease association heterogeneity.

See Supplementary Excel file

**Supplementary Table 16. Numerical results for cell type-level analyses for the TS FACS data.** For each pair of cell type and disease/trait, we report the proportion of significantly associated cells (FDR<0.1), FDR for cell type-disease association, and FDR for within-cell type disease association heterogeneity.

See Supplementary Excel file

**Supplementary Table 17. Control traits and cell types for different versions of `scDRS`.** We report the 20 traits and the corresponding expected and unexpected control cell types[8,23,34,35,67,68] in the TMS FACS data for evaluating the performance of `scDRS` under different parameter settings.

See Supplementary Excel file

**Supplementary Table 18. Numerical results for optimizing parameters of `scDRS`.** Each row represents a version of `scDRS`. We report the mean and SE of the normalized t-statistics for each version of `scDRS`, and the mean, SE, and p-value of the difference of the normalized t-statistics between each version and the default version (top 1,000 genes, GWAS Z + single-cell VS weights, 10-kb window; details in Supp. Fig. 12). As a reference, we also report the performance of 3 oracle methods (not available in practice due the use of oracle information): 1.41 (SE 0.033) for the method that uses the best out of the 6 gene selection methods for each trait while fixing the other parameters as the default; 1.26 (SE 0.029) for the method that uses the best out of the 4 gene weighting methods for each trait while fixing the other parameters as the default; 1.43 (SE 0.032) for the method that uses the best combination out of the 6 gene selection methods and 4 gene weighting methods for each trait while fixing the other parameters as the default.

| Results for 26 main traits | $N_{\text{trait}}$ | Gene set overlap (SE) | Score corr. (SE) | Score corr. excl. overlapping genes (SE) |
|---|---|---|---|---|
| Immune | 10 | 230.7 (10.0) | 0.507 (0.007) | 0.162 (0.009) |
| Brain | 7 | 199.3 (4.4) | 0.440 (0.012) | 0.169 (0.012) |
| Metabolic | 9 | 275.4 (9.6) | 0.355 (0.011) | 0.016 (0.004) |
| Inter-category | 26 | 110.9 (1.1) | 0.018 (0.003) | -0.095 (0.003) |
| Results for all 74 traits | | | | |
| Brain | 21 | 154.5 (3.8) | 0.328 (0.009) | - |
| Heart | 6 | 295.8 (56.5) | 0.408 (0.051) | - |
| Blood/immune | 21 | 167.0 (3.0) | 0.336 (0.009) | - |
| Metabolic | 13 | 230.3 (6.4) | 0.270 (0.009) | - |
| Other | 13 | 120.2 (4.1) | 0.183 (0.017) | - |
| Inter-category | 74 | 105.7 (0.5) | 0.061 (0.001) | - |

**Supplementary Table 19. Gene set overlap and scDRS score correlation across TMS FACS cells between traits.** We report the number of traits, gene overlap, scDRS score correlation, and correlation of scDRS scores computed using only non-overlapping genes between a pair of traits, for the 26 immune, brain, and metabolic traits considered in the individual-cell level analyses (upper). We also report the number of traits, gene overlap, and scDRS score correlation for all 74 traits grouped by categories (lower). SEs were computed via jackknifing the traits.

| Trait | CD4.P | CD4.Var | CD4.P.joint | CD8.P | CD8.Var | CD8.P.joint |
|---|---|---|---|---|---|---|
| IBD | **0.001** | 0.282 | **0.005** | 0.021 | 0.083 | 0.021 |
| CD | **0.001** | 0.202 | **0.004** | 0.026 | 0.087 | 0.006 |
| UC | **0.004** | 0.159 | 0.067 | 0.441 | -0.000 | 0.548 |
| RA | 0.118 | 0.024 | 0.268 | 0.261 | 0.006 | 0.569 |
| MS | 0.223 | 0.009 | 0.238 | 0.129 | 0.020 | 0.418 |
| AIT | **0.002** | 0.188 | 0.007 | 0.038 | 0.064 | 0.194 |
| HT | **0.003** | 0.153 | 0.030 | 0.100 | 0.028 | 0.267 |
| Eczema | 0.011 | 0.087 | 0.250 | 0.819 | 0.016 | 0.888 |
| ASM | 0.035 | 0.059 | 0.387 | 0.325 | 0.001 | 0.580 |
| RR-ENT | 0.060 | 0.057 | 0.245 | 0.269 | 0.004 | 0.356 |
| Height | 0.282 | 0.004 | 0.739 | 0.825 | 0.009 | 0.629 |

**Supplementary Table 20. Numerical results for correlations between scDRS disease scores and T cell effectorness gradients in Fig. E.** We first regressed the scDRS disease score against the CD4 (resp., CD8) effectorness gradient for each of the 10 autoimmune diseases and the negative control trait height. We report p-values for significant positive correlation between the scDRS disease score and the effectorness gradients (“CD4.P”/“CD8.P”; MC test) and variance explained (“CD4.Var”/“CD8.Var”). We then jointly regressed the scDRS disease score against the CD4 (resp., CD8) effectorness gradient and the cluster labels (encoded as dummy variables). We report p-values for significant positive correlation between the scDRS disease score and the effectorness gradients (“CD4.P.joint”/“CD8.P.joint”; MC test). P-values smaller than 0.005 were highlighted in bold font.

See Supplementary Excel file

**Supplementary Table 21. Gold standard gene sets.** We report matched Experimental Factor Ontology (EFO), corresponding EFO label, corresponding putative drug target gene set, number of putative drug target genes, matched Mendelian disorder, corresponding Mendelian disease gene set, and number of Mendelian disease genes for 27 diseases with putative drug target gene sets (from Open Targets) and 45 diseases/traits with Mendelian gene sets (from Freund et al.[69]); a disease/trait may have both drug target and Mendelian disease gene set. Specifically, for the 10 autoimmune diseases in Fig. 4F, the immune dysregulation Medelian disease gene set was used for HT and RR-ENT while the corresponding drug target gene sets were used for the other 8 autoimmune diseases.

See Supplementary Excel file

**Supplementary Table 22. Numerical results for comparison to gold standard gene sets in Fig. 4F.** We report excess overlap and $-\log_{10}$ p-value for comparison to drug target and Mendelian disease gene sets respectively for each disease gene prioritization method (`scDRS`/MAGMA) and each of the 27 GWAS diseases with drug target gene sets and 45 diseases/traits with Mendelian disease gene sets.

| Cano-Gamez et al. | IBD | CD | UC | RA | MS | AIT | HT | Eczema | ASM | RR-ENT | Height |
|---|---|---|---|---|---|---|---|---|---|---|---|
| TCM1 (Th17/iTreg) | .099 | .081 | ns | ns | .110 | .127 | .179 | ns | ns | ns | ns |
| TCM2 (Th0) | ns | .081 | ns | ns | ns | ns | ns | ns | ns | ns | ns |
| TEM (Th0) | ns | ns | ns | ns | ns | ns | .092 | ns | ns | ns | ns |
| TEM (Th17/iTreg) | .015 | .088 | ns | ns | ns | .073 | .011 | .081 | ns | ns | ns |
| TEMRA (Th0) | ns | .124 | ns | ns | ns | ns | ns | ns | ns | ns | ns |
| TN (Th0) | ns | .081 | ns | ns | ns | .127 | .092 | ns | ns | ns | ns |
| TN (Th17/iTreg) | .015 | .022 | .022 | .077 | ns | .022 | .029 | .011 | .044 | .022 | ns |
| TN (Th2) | ns | .081 | ns | ns | ns | ns | ns | ns | ns | ns | ns |
| TN (iTreg) | ns | ns | ns | ns | ns | ns | .179 | ns | ns | .165 | ns |
| nTreg (Th0) | .015 | .022 | .033 | .022 | .066 | .022 | .011 | .011 | .187 | .183 | ns |
| Nathan et al. | | | | | | | | | | | |
| $CD4^+$ $CD161^+$ Th2 | .167 | ns | ns | ns | ns | ns | ns | .035 | .122 | .058 | ns |
| $CD4^+$ *RORC*$^+$ Treg | .167 | ns | .087 | .014 | ns | .014 | .014 | .022 | .029 | .029 | ns |
| $CD4^+$ Th2 | ns | ns | ns | ns | ns | ns | ns | .022 | .140 | .124 | ns |
| $CD4^+$ Th17 | ns | ns | ns | ns | ns | ns | .156 | .022 | .043 | .041 | ns |
| $CD4^+$ Th17/1 | .167 | .145 | ns | ns | ns | ns | ns | .022 | .043 | .041 | ns |
| $CD4^+$ Treg | ns | ns | ns | .014 | ns | .014 | .014 | ns | ns | ns | ns |
| $CD4^+$ activated | .167 | ns | .159 | .068 | ns | .159 | .145 | .077 | .043 | .041 | ns |
| $CD4^+$ lncRNA | ns | ns | ns | ns | ns | ns | ns | .112 | .161 | .041 | ns |

**Supplementary Table 23. Cell type-disease associations for T cell subtypes in Cano-Gamez & Soskic et al. and Nathan et al. data sets.** We report FDR for significant cell type-disease associations (FDR<0.2) in the Cano-Gamez & Soskic et al. and Nathan et al. data sets (Supp. Table 2); the more lenient threshold of 0.2 was used to also include borderline associations. We performed cell type-disease associations between each cell type in the two data sets and each of the 10 autoimmune diseases (and height, a negative control trait) using the `scDRS`-based MC test. We applied FDR correction for each data set and each disease separately across all cell types in the data set (22 cell types in Cano-Gamez & Soskic et al. and 29 cell types in Nathan et al.). For cell types in Cano-Gamez & Soskic et al., "TCM1 (Th17/iTreg)" means cells induced from central memory T cells (TCM) via TCR/CD28-activation and the Th17/iTreg cytokine condition, and similar for others ("TN" for naive T cells, "TEM" for effector memory T cells, "TEMRA" for effector memory T cells reexpressing CD45RA); "nTreg (Th0)" means natural Tregs induced via only TCR/CD28-activation (without cytokines).

See Supplementary Excel file

**Supplementary Table 24. Results for cell type-level analyses for the Zeisel & Muñoz-Manchado et al. data.** For each pair of cell type and disease/trait, we report the proportion of significantly associated cells (FDR<0.1), FDR for cell type-disease association, and FDR for within-cell type disease association heterogeneity.

| | Dorsal.P | Dorsal.Var | Ventral.P | Ventral.Var | Proximal.P | Proximal.Var | Distal.P | Distal.Var | Deep.P | Deep.Var | Superficial.P | Superficial.Var |
|---|---|---|---|---|---|---|---|---|---|---|---|---|
| MDD | **0.001** | 0.260 | 0.75 | 0.005 | **0.002** | 0.152 | 0.2 | 0.004 | **0.001** | 0.208 | 0.8 | 0.004 |
| SCZ | **0.001** | 0.184 | 0.68 | 0.003 | **0.001** | 0.196 | 0.81 | 0.004 | **0.001** | 0.119 | 0.83 | 0.004 |
| BP | **0.001** | 0.186 | 0.63 | 0.001 | **0.001** | 0.227 | 0.52 | 0.000 | **0.001** | 0.125 | 0.66 | 0.001 |
| NRT | **0.002** | 0.137 | 0.28 | 0.004 | **0.001** | 0.217 | 0.64 | 0.001 | **0.001** | 0.150 | 0.52 | 0.000 |
| Smoking | 0.007 | 0.110 | 0.17 | 0.014 | **0.001** | 0.177 | 0.55 | 0.000 | **0.001** | 0.142 | 0.26 | 0.002 |
| ECOL | **0.001** | 0.170 | 0.41 | 0.000 | **0.001** | 0.225 | 0.4 | 0.000 | **0.001** | 0.169 | 0.66 | 0.001 |
| BMI | 0.019 | 0.064 | 0.073 | 0.026 | **0.001** | 0.292 | 0.63 | 0.001 | **0.001** | 0.143 | 0.41 | 0.000 |
| Height | 0.37 | 0.002 | 0.54 | 0.000 | 0.006 | 0.066 | 0.71 | 0.001 | 0.2 | 0.007 | 0.98 | 0.019 |

**Supplementary Table 25. Numerical results for correlations between `scDRS` disease scores and inferred spatial coordinates in Fig. 5B.** We separately regressed the `scDRS` scores of each of the 7 brain traits (and height, a negative control trait) against each of the 6 inferred spatial coordinates across the 827 CA1 pyramidal neurons. We report p-values for significant positive correlation between the `scDRS` disease score and the inferred spatial coordinates (MC test) and variance explained. P-values smaller than 0.005 were highlighted in bold font.

| | Polyploidy.P | Pericentral.P | Periportal.P | Variance explained |
|---|---|---|---|---|
| TG | **0.001** | 0.730 | 0.009 | 0.568 |
| HDL | 0.006 | 0.620 | 0.044 | 0.422 |
| LDL | 0.007 | 0.350 | 0.006 | 0.443 |
| TC | **0.002** | 0.434 | **0.005** | 0.461 |
| TST | **0.001** | 0.781 | **0.005** | 0.611 |
| ALT | **0.001** | 0.515 | 0.006 | 0.615 |
| ALP | **0.001** | 0.480 | **0.001** | 0.599 |
| SHBG | **0.001** | 0.616 | 0.017 | 0.601 |
| TBIL | **0.001** | 0.751 | 0.241 | 0.527 |
| Height | 0.634 | 0.939 | 0.241 | 0.115 |

**Supplementary Table 26. Numerical results for correlations between `scDRS` disease scores and inferred ploidy and zonation scores in Fig. 5D.** We jointly regressed the `scDRS` scores of each of the 9 metabolic traits (and height, a negative control trait) on the polyploidy score, pericentral score, and periportal score. We report p-values for significant positive correlation between the `scDRS` disease score and the polyploidy score, pericentral score, and periportal score (MC test) and variance explained. P-values smaller than 0.005 were highlighted in bold font.

| | TMS FACS | TMS Droplet | Aizarani | Halpern | Richter | Taychameekiatchai |
|---|---|---|---|---|---|---|
| 4n hepatocyte (vs. 2n) | **0.001** | **0.001** | **0.001** | **0.001** | **0.001** | **0.001** |
| polyploid (Cdk1 ko) | **0.001** | **0.001** | **0.001** | 0.038 | **0.001** | **0.001** |
| large hepatocyte (vs. small) | **0.001** | **0.001** | **0.001** | **0.001** | **0.001** | **0.001** |
| 2n hepatocyte (vs. 4n) | 0.470 | 0.009 | 0.542 | **0.003** | **0.001** | 0.014 |
| diploid (Cdk1 ko) | **0.001** | **0.001** | 0.321 | 0.028 | **0.001** | 0.148 |
| diploid (PH) | **0.001** | **0.001** | 0.156 | **0.001** | **0.002** | **0.001** |

**Supplementary Table 27. Correlation between the `scDRS` polyploidy score and `scDRS` score for other ploidy signatures.** We correlated our polyploidy score (based on DEGs for PH vs. pre-PH) with other ployploidy and diploidy signatures for the 6 data sets. We report p-values for significant positive correlation for the polyploidy signatures (first three rows) and significant negative correlation for the diploidy signatures (last three rows). P-values smaller than 0.005 were highlighted in bold font.

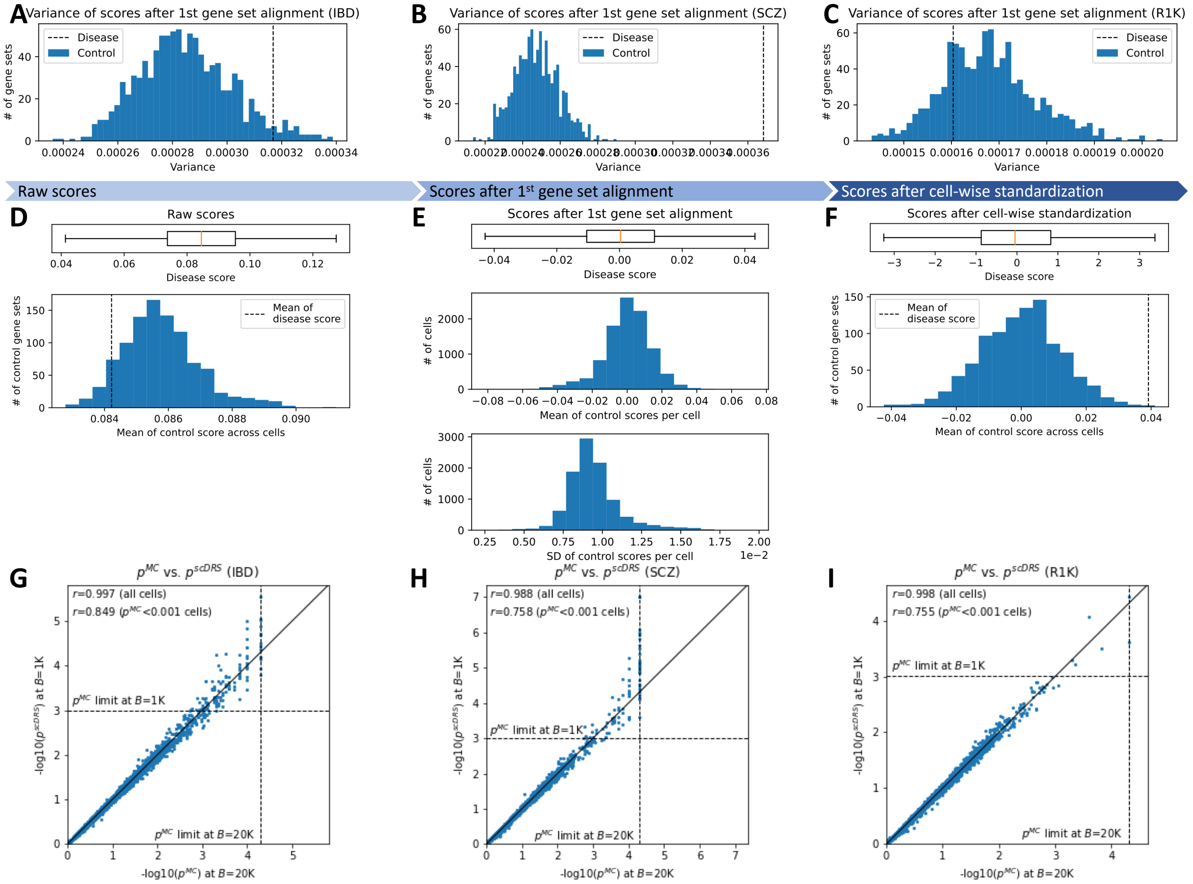


**Supplementary Figure 1. Details of `scDRS` method.** Results are based 10,000 cells subsampled from TMS FACS and 3 sets of putative disease genes: IBD, SCZ, and R1K (1,000 random genes); we used binary gene sets for simplicity. **(A-C)** Variance of raw disease scores (dashed line) and raw control scores (histogram) computed across cells for different gene sets. Informative disease gene sets (IBD and SCZ) have higher disease score variances while the uninformative gene set (R1K) has comparable values for the disease score variance and control score variances. We considered raw scores after the first gene set alignment to remove potential mismatch of expression mean and variance across gene sets. **(D)** Comparison between raw disease scores and raw control scores (before Box 1, step 3a). The upper panel shows the distribution of disease scores and the lower panel shows the histogram of the mean of control scores (computed for each control gene set across cells). There is a moderate level of mismatch of mean expression across control gene sets (ratio between the SD of mean control scores across control gene sets and SD of disease scores across cells is 6.8%); this mismatch is corrected by the first gene set alignment (Box 1, step 3a). **(E)** Comparison between disease scores and control scores after the first gene set alignment (before Box 1, step 3b). The upper panel shows the distribution of disease scores and the middle (resp. lower) panel shows the histogram of the per-cell mean (resp. SD) of control scores (computed for each cell across control gene sets). There is a high level of mismatch of the control score distribution (mean and SD) across cells; this mismatch is corrected by the cell-wise standardization (Box 1, step 3b). **(F)** Comparison between disease scores and control scores after cell-wise standardization (before Box 1, step 3c). The upper panel shows the distribution of disease scores and the lower panel shows the histogram of the mean of control scores (computed for each control gene set across cells). There is a mild level of mismatch of mean expression across control gene sets (ratio between the SD of mean control scores across control gene sets and SD of disease scores across cells is 1%); this mismatch is corrected by the second gene set alignment (Box 1, step 3c). Panels D-F are based on the IBD results. **(G-I)** Comparison between MC p-values with $B=$20,000 and `scDRS` p-values with $B=$1,000 for IBD, SCZ, and R1K. Each dot denotes a cell and the $p^{\mathrm{MC}}$ limit 1/(1+B) is the smallest MC p-value that an MC test with $B$ MC samples can achieve. The `scDRS` p-values are highly correlated with the MC p-values obtained with $B=$20,000, both across all cells and across cells with $p^{\mathrm{MC}}<0.001$ ($p^{\mathrm{MC}}$ limit at $B=1{,}000$; corresponding to cells whose ideal MC p-values (with $B=\infty$) are small and require `scDRS` to extrapolate beyond the $p^{\mathrm{MC}}$ limit at the given number of MC samples), suggesting that `scDRS` can reliably approximate the ideal p-values obtained with an infinite number of MC samples. The dots on the vertical dashed line correspond to those whose ideal MC p-values are smaller than the $p^{\mathrm{MC}}$ limit at $B=$20,000; it is not surprising that the `scDRS` p-values, approximating the ideal MC p-values, are on average smaller than the MC p-values for these cells.

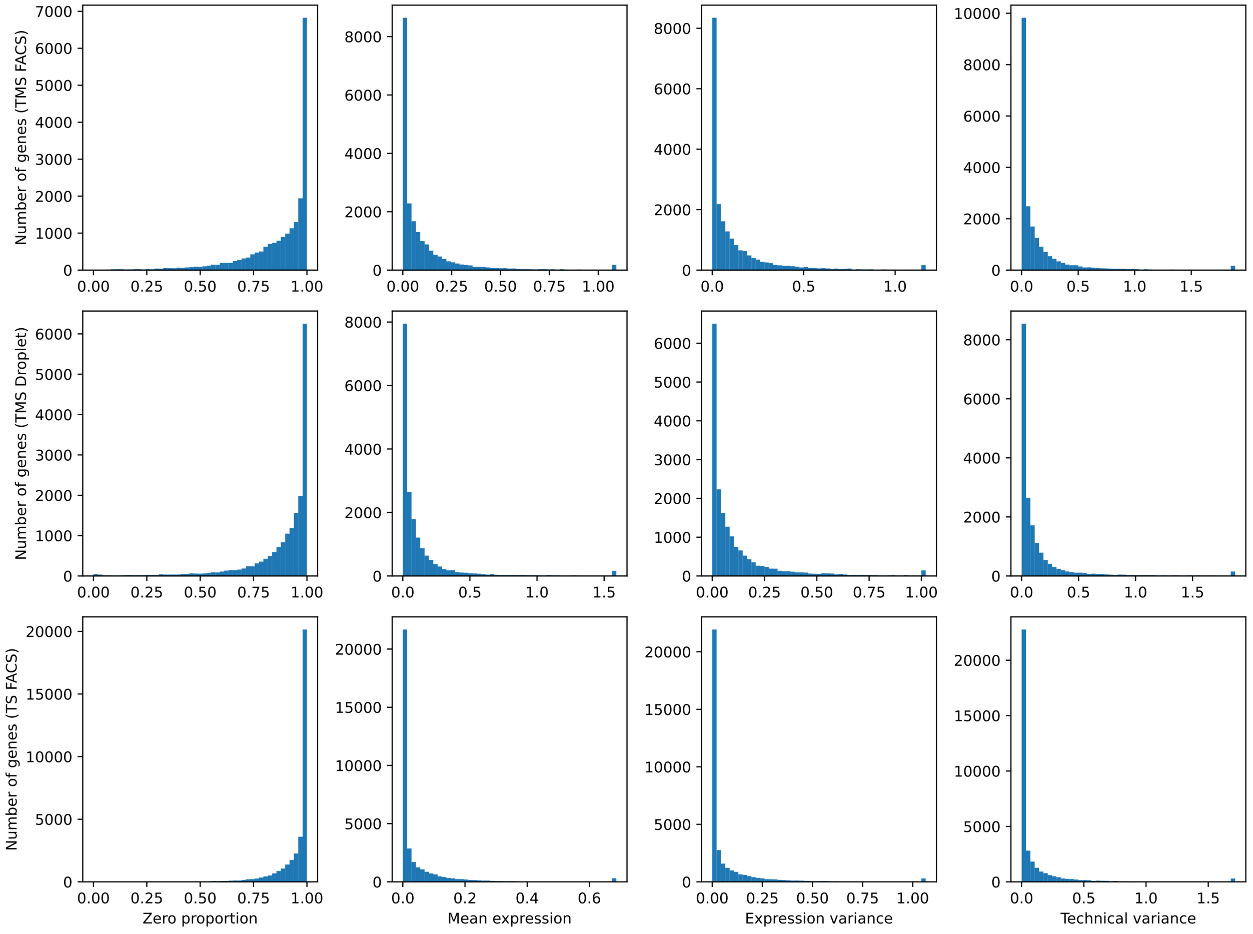


**Supplementary Figure 2. Distribution of zero proportion, mean expression, expression variance, and technical variance across genes for the TMS FACS, TMS Droplet, and TS FACS data.**

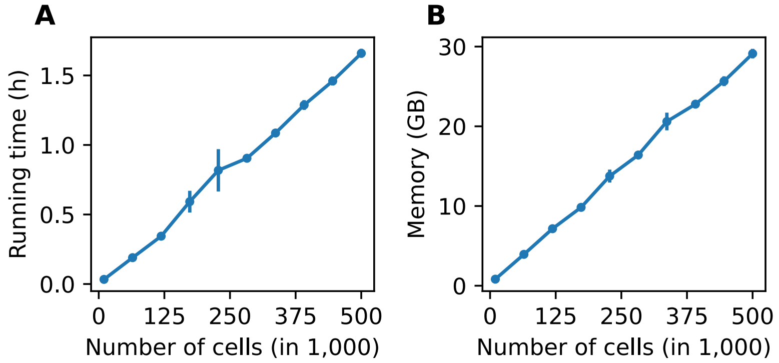


**Supplementary Figure 3. Computational cost of `scDRS`.** Running time (A) and memory usage (B) as a function of the number of cells. We created subsampled data sets from the Nathan et al. data set[14] (500,089 cells and 17,256 genes), and ran `scDRS` under the default setting (1,000 control gene sets) for each data set with a disease gene set of 1,000 genes. Each setting was repeated 10 times and all experiments were performed using one core of Intel Xeon Platinum 8268 CPU @ 2.90GHz processor. 95% confidence intervals were provided based on the 10 repetitions.

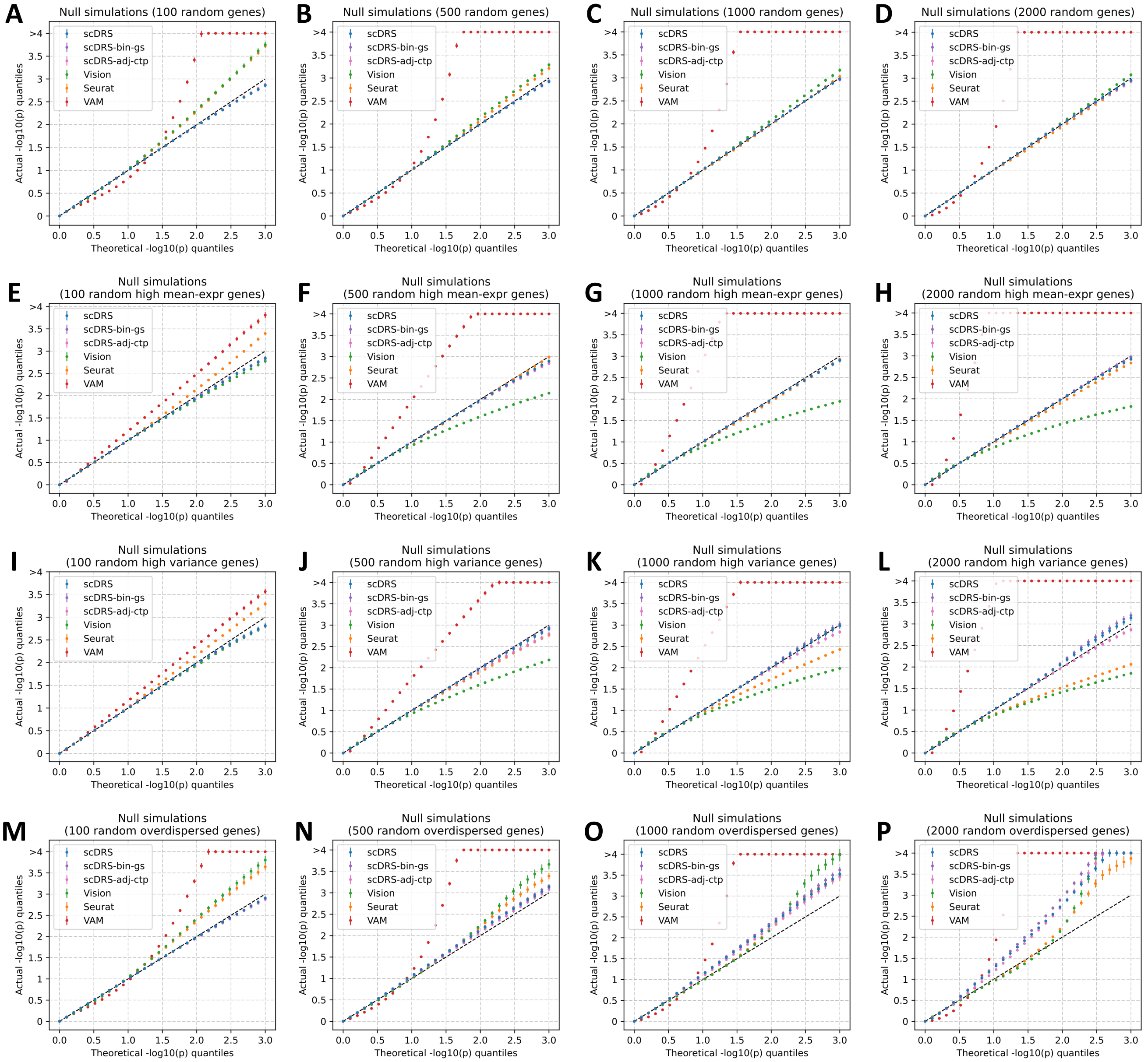


**Supplementary Figure 4. Additional null simulations.** We performed null simulations for various numbers of putative disease genes (100, 500, 1,000, and 2,000 for the four columns respectively) and various types of genes to randomly sample from: all genes (first row), and top 25% genes with high expression (second row), top 25% genes with high expression variance (third row), top 25% overdispersed genes (fourth row). We considered two additional versions of `scDRS`: scDRS-bin-gs (binary gene sets instead of MAGMA z-score gene weights) and scDRS-adj-ctp (adjusting for cell type proportion). For scDRS-adj-ctp, we simulated random biased gene sets (high-mean/high-variance/overdispersed) based on the balanced data (inversely weighting cells by cell type proportion) to better match the model assumption, namely testing for excess expression relative to cells in the balanced data. In each panel, the x-axis denotes theoretical $-\log_{10}$ p-value quantiles and the y-axis denotes actual $-\log_{10}$ p-value quantiles for different methods. The 3 versions of `scDRS` produced well-calibrated p-values in most settings and suffered slightly inflated type I error in panels O and P, possibly because it is hard to match a large number of overdispersed putative disease genes using the remaining set of genes. In comparison, all other methods are less well-calibrated and are particularly problematic when the numbers of putative disease genes are small. All experiments were repeated 100 times and 95% confidence intervals were provided.

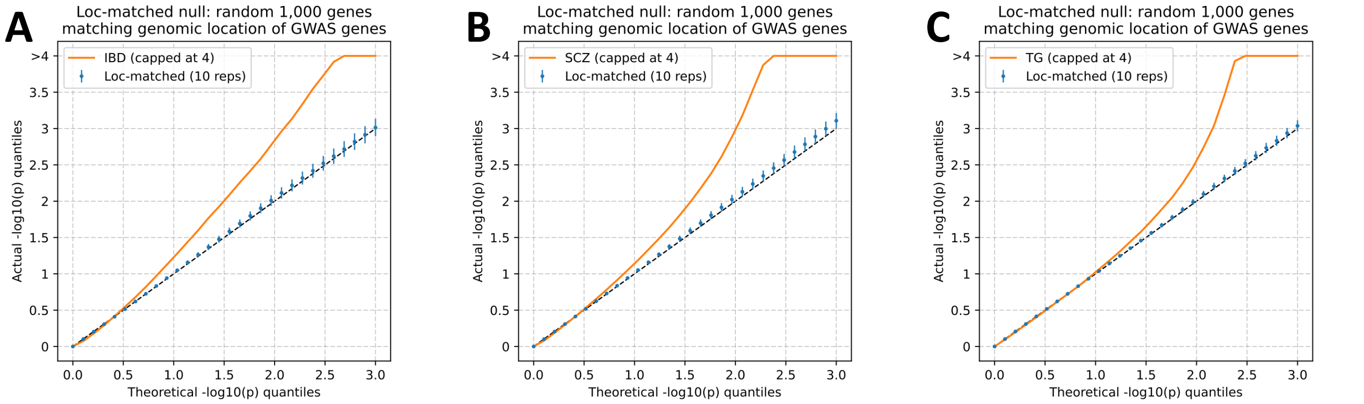


**Supplementary Figure 5. Null simulations with genomic location-matched null gene sets.** We generated location-matched null gene sets from GWAS putative disease gene sets as follows. First, we divided all genes on each chromosome into 20 equal-sized bins based on genomic location. Next, for each chromosome-location bin, we randomly sampled an equal number of location-matched null genes (without replacement) as the putative disease genes in the same bin. For simplicity, we used binary gene sets (without GWAS gene weights) and computed `scDRS` p-values across the full TMS FACS data. The figure shows Q-Q plots of the `scDRS` p-values for both the GWAS putative disease gene set and the location-matched null gene sets for 3 diseases/traits: IBD, SCZ, and TG. 95% confidence intervals were provided based on 10 repetitions of the location-matched null gene sets.

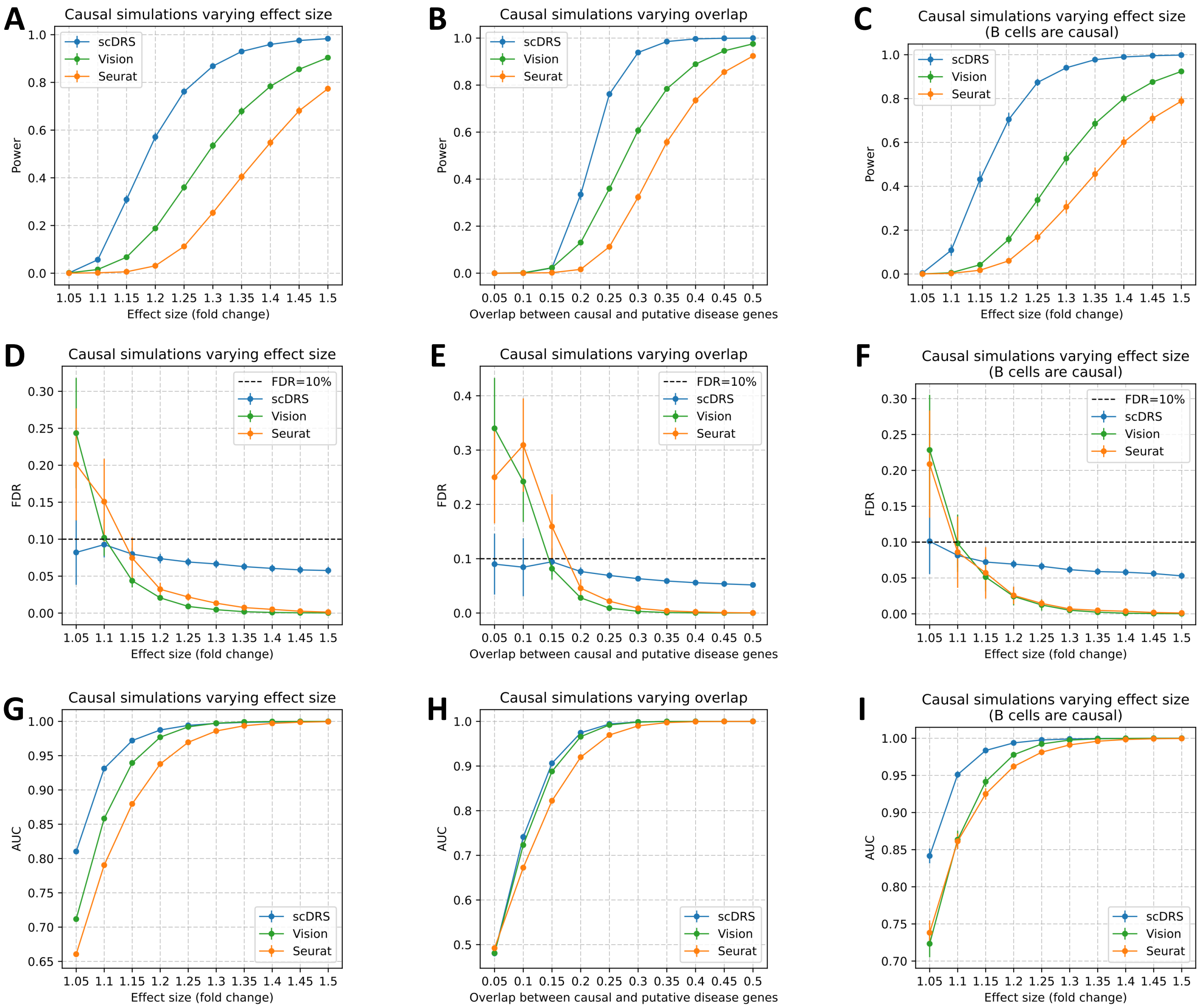


**Supplementary Figure 6.** **Additional causal simulations.** We performed three sets of causal simulations: (1) varying effect size from 5% to 50% while fixing 25% overlap (first column), (2) varying level of overlap from 5% to 50% while fixing 25% effect size (second column), (3) assigning the 528 B cells in the subsampled data to be causal (instead of the 500 randomly selected cells; varying effect size while fixing 25% overlap; third column). We report the power (first row), FDR (second row), and AUC for classifying causal from non-causal cells based on the p-values (third row). `scDRS` outperformed other methods under all metrics. All experiments were repeated 100 times and 95% confidence intervals were provided.

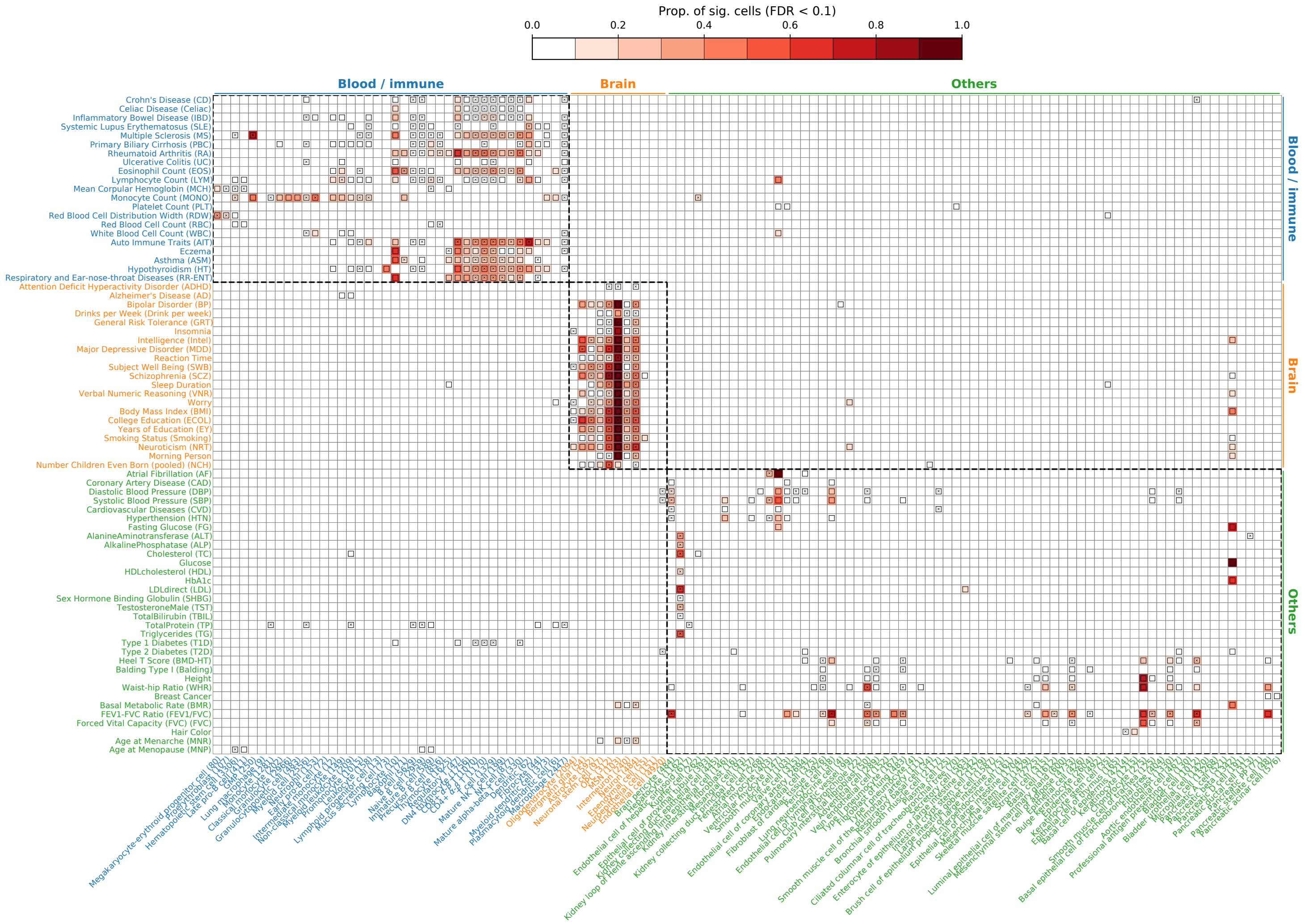


**Supplementary Figure 7. Complete results for disease associations at the cell type level for 74 diseases/traits and 120 cell types in the TMS FACS data in Fig. 3.** Each row represents a disease/trait and each column represents a cell type (with number of cells indicated in parentheses). Heatmap colors for each cell type-disease pair denote the proportion of significantly associated cells (FDR<0.1 across all cells for a given disease). Squares denote significant cell type-disease associations (FDR<0.05 across all pairs of the 120 cell types and 74 diseases/traits; 597 significant pairs; p-values via MC test; Methods). Cross symbols denote significant heterogeneity in association with disease across individual cells within a given cell type (FDR<0.05 across all pairs; 273 significant pairs; p-values via MC test; Methods). Heatmap colors and cross symbols are omitted for cell type-disease pairs with non-significant cell type-disease associations. Within the blood/immune block (40 cell types and 21 diseases/traits), 136 of 264 cell type-disease pairs with significant association also had significant heterogeneity. Within the brain block (11 cell types and 21 diseases/traits), 64 of 133 cell type-disease pairs with significant association also had significant heterogeneity. Within the other block (69 cell types and 32 diseases/traits), 54 of 146 cell type-disease pairs with significant association also had significant heterogeneity. We briefly discuss the results for FEV1/FVC. We identified 20 cell types associated with FEV1/FVC (FDR<0.05), including 5 lung cell types and 15 cell types from other tissues. They can be categorized into 5 sets of associations: (1) type II pneumocyte (2) skin-related cells (3) smooth muscle cells (4) fibroblast-and-MSC-like cells (5) pericyte-like cells. The first 4 sets of associations are consistent with a previous work[70]. The 5th set of pericyte associations is also plausible because pericytes are known to regulate lung morphogenesis[71]. We note that the cell type associations from the lung are more likely to be causal and those from the other tissues are more likely tagging the causal cell types due to shared expression. Numerical results are reported in Supp. Table 14.

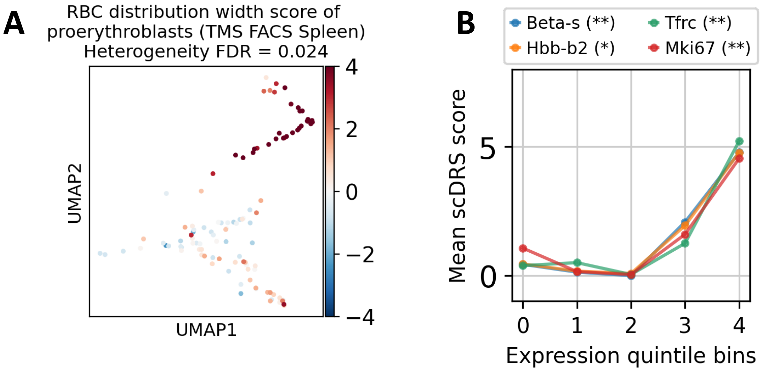


**Supplementary Figure 8.** **Heterogeneous subpopulations of proerythroblasts associated with red blood cell distribution width (RDW). (A)** Significant heterogeneity (FDR=0.024) of proerythroblasts (in the spleen) in association with RDW. **(B)** Expression levels of proerythroblast marker genes are significantly positively correlated with the `scDRS` disease score. The x-axis denotes marker gene expression quintile bins and the y-axis denotes average `scDRS` disease score for each bin. * denotes $P < 0.05$ and ** denotes $P < 0.005$. The heterogeneous association levels of proerythroblasts with RDW may correspond to the different differentiation stages of proerythroblasts[72]. Of note, the `scDRS` disease score was not correlated with age ($P = 0.12$) or sex ($P = 0.39$).

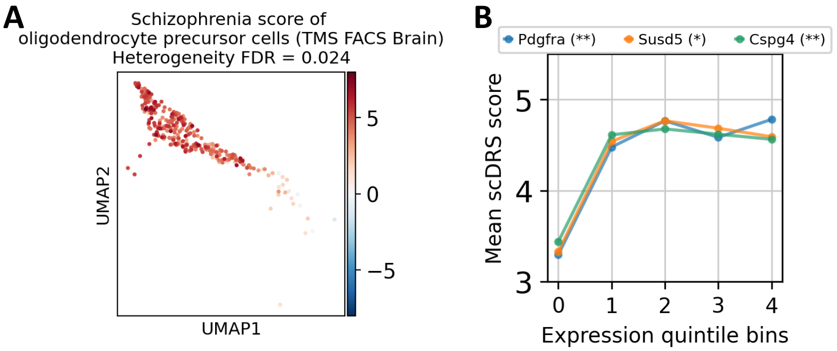


**Supplementary Figure 9.** **Heterogeneous subpopulations of oligodendrocyte precursor cells associated with schizophrenia (SCZ). (A)** Significant heterogeneity (FDR=0.024) of oligodendrocyte precursor cells (in the brain non-myeloid) in association with SCZ. **(B)** Expression levels of oligodendrocyte precursor cell marker genes are significantly positively correlated with the `scDRS` disease score. The x-axis denotes marker gene expression quintile bins and the y-axis denotes average `scDRS` disease score for each bin. * denotes $P < 0.05$ and ** denotes $P < 0.005$. The heterogeneous association levels of oligodendrocyte precursor cells with SCZ may correspond to the different developmental stages of oligodendrocyte precursor cells[73]. Of note, the `scDRS` disease score was slightly higher in male than female ($P = 0.010$), and was not correlated with age ($P = 0.080$).

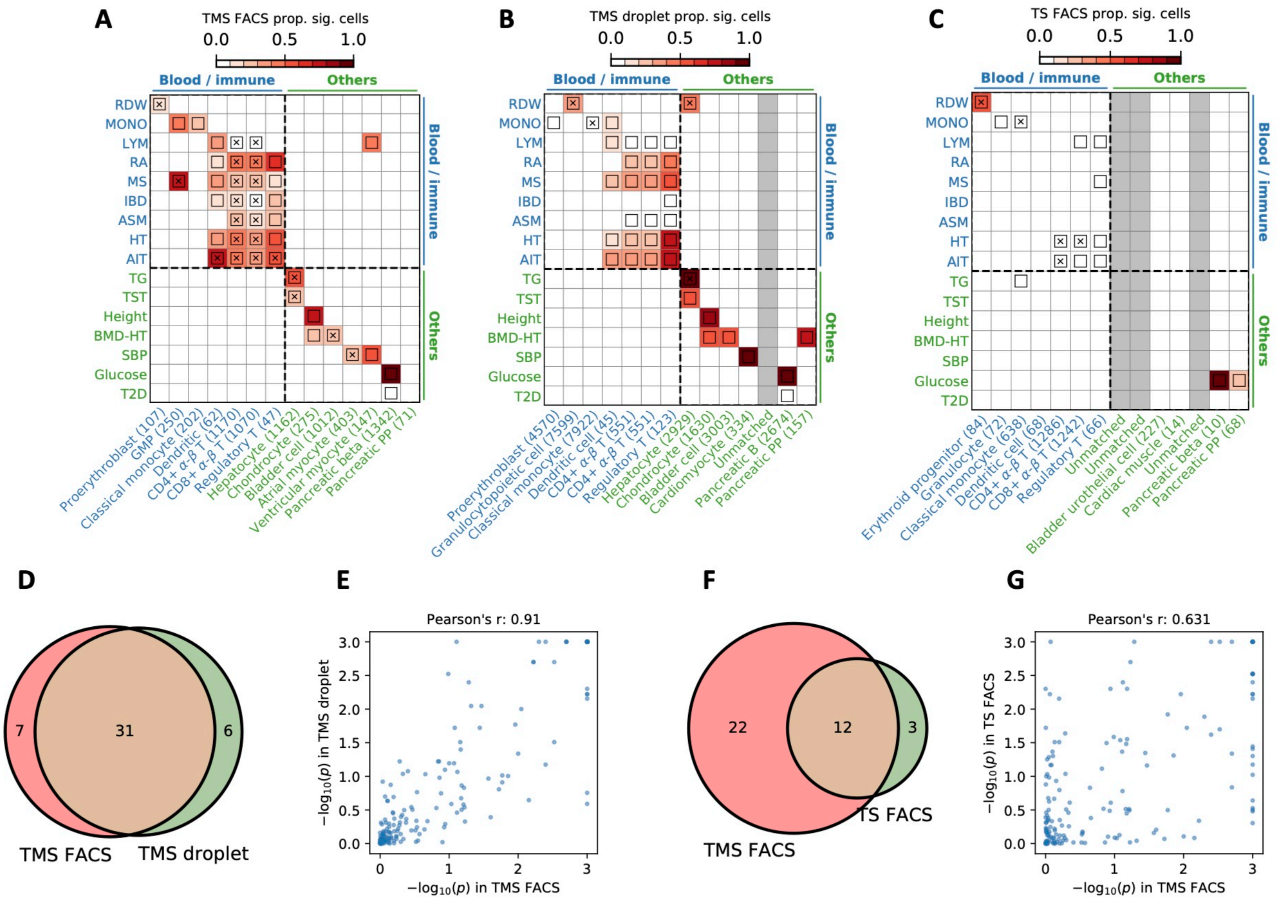


**Supplementary Figure 10. Comparison of cell type-level disease association results between TMS FACS and TMS droplet (different technology), TS FACS (different species).** **(A-C)** Results for disease association at the cell type-level for TMS FACS, TMS droplet, and TS FACS for diseases and cell types in the blood/immune block (upper left) and the other cell types/diseases block (lower right) in Fig. 3 (TMS droplet and TS FACS do not contain brain data; Supp. Table 6,7). The plotting style is same as Fig. 3. Heatmap colors for each cell type-disease pair denote the proportion of significantly associated cells (FDR<0.1); squares denote significant cell type-disease associations (FDR<0.05); and cross symbols denote significant heterogeneity in association with disease across individual cells within a given cell type (FDR<0.05). Heatmap colors (>10% of cells associated) and cross symbols are omitted for cell type-disease pairs with non-significant cell type-disease associations via MC test. We matched each TMS FACS cell type using the closest cell type in the TMS droplet and TS FACS data; unmatched cell types were colored in grey. **(D)** Overlap of significant cell type-disease associations between TMS FACS and TMS droplet ($P = 2.8 \times 10^{-24}$, Fisher's exact test). **(E)** Correlation of $-\log_{10}$ p-values for cell type-disease associations between TMS FACS and TMS droplet. **(F)** Overlap of significant cell type-disease associations between TMS FACS and TS FACS ($P = 1.3 \times 10^{-7}$, Fisher's exact test). **(G)** Correlation of $-\log_{10}$ p-values for cell type-disease associations between TMS FACS and TS FACS. We determined that the results are highly consistent between TMS FACS and TMS droplet, and are reasonably consistent between TMS FACS and TS FACS. Our method is underpowered in the TS FACS data, possibly due to the smaller sample size (27K cells in TS FACS vs. 110K cells in TMS FACS). The current TS FACS data corresponds to the initial data release and there will likely be more cells in future releases[59].

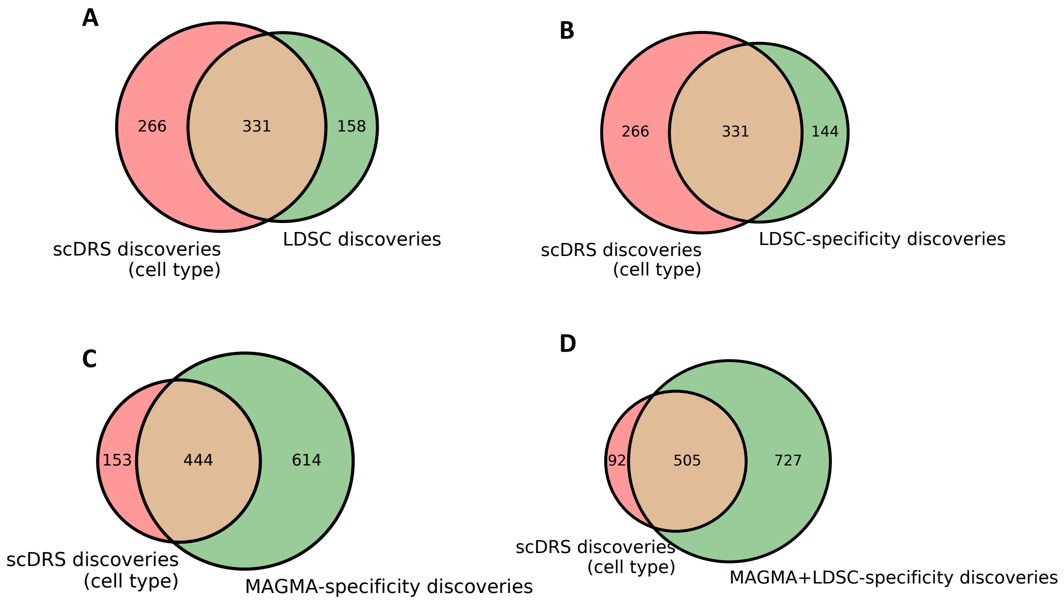


**Supplementary Figure 11. Comparison of cell type-disease associations to alternative methods.** We compared `scDRS` to 4 alternative methods for cell type-level analyses across the same 120 TMS FACS cell types and 74 diseases: LDSC-SEG[8], LDSC-specificity[9], MAGMA-specificity[9], and the combined method[9] (mean -log10 between LDSC-specificity and MAGMA-specificity). For `scDRS`, cell type-diseases associations were computed using the MC test. For LDSC-SEG, we identified specifically-expressed genes for each of the 120 cell types using one-versus-rest differential expression analysis ("rank_genes_groups" with option "t-test_overestim_var" in scanpy[74]; top 1,000 genes to be consistent with `scDRS`). We used 100-kb windows around the gene body to map genes to variants (default setting in LDSC-SEG) and applied S-LDSC[53] conditional on the 52 baseline annotations (baseline v1.2) to identify disease-relevant cell types. For both LDSC-specificity and MAGMA-specificity, we used the top 10% high-specificity genes as defined in Bryois et al.[9] (instead of differentially-expressed genes used in LDSC-SEG). We computed FDR across all pairs of cell types and diseases/traits and used a significance threshold of 0.05. **(A)** Venn diagram of `scDRS` and LDSC-SEG discoveries ($P = 2.9 \times 10^{-306}$, Fisher's exact test). **(B)** Venn diagram of `scDRS` and LDSC-specificity discoveries ($P = 4.1 \times 10^{-313}$, Fisher's exact test). **(C)** Venn diagram of `scDRS` and MAGMA-specificity discoveries ($P = 1.3 \times 10^{-312}$, Fisher's exact test). **(D)** Venn diagram of `scDRS` and the combined method discoveries ($P < 4.1 \times 10^{-313}$, Fisher's exact test). We note that `scDRS` identified some biologically plausible associations that were missed by other methods, including pancreatic PP cells and BMD-HT[75] (`scDRS` FDR=0.063 vs. all other methods FDR>0.95), GMPs and MS[76,77] (FDR=0.020 vs. FDR>0.30).

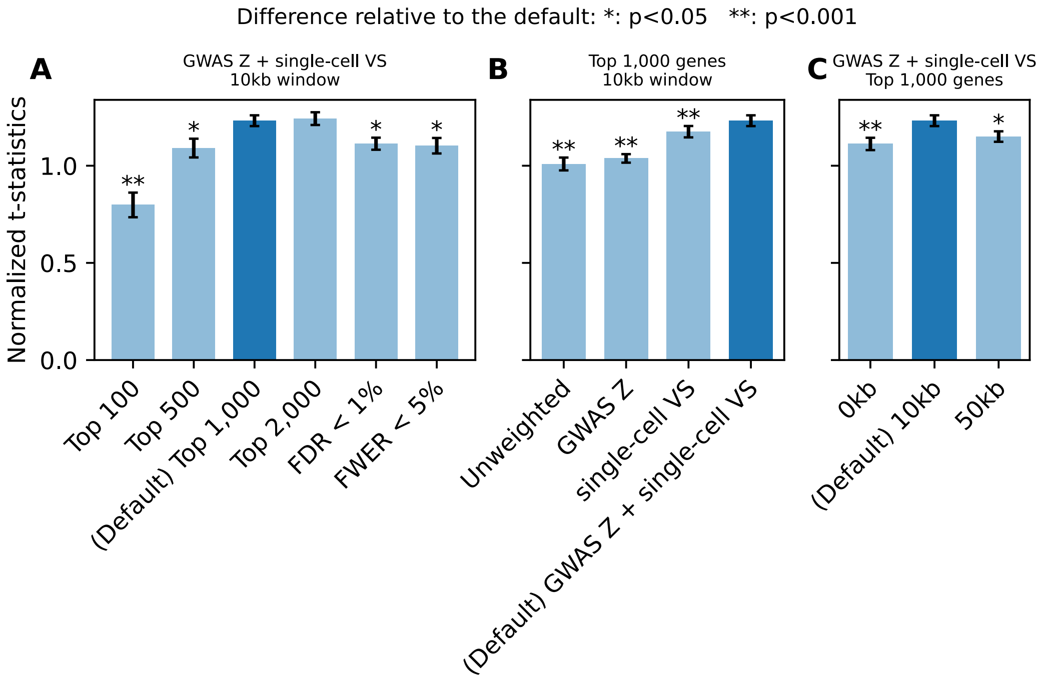


**Supplementary Figure 12.** **Optimizing parameters of scDRS based on expected and unexpected control cell types across 20 traits.** We considered different versions of `scDRS` by varying (1) method for selecting putative disease genes (2) method for selecting weights for the disease genes (3) MAGMA window size. We considered 6 methods for selecting putative disease genes: top 100, top 500, top 1,000 (default), top 2,000, FDR<1%, and FWER<5% genes (with the number of top genes constrained between 100 and 2,000 for the latter 2 methods). We considered 4 methods for selecting gene weights: unweighted, GWAS MAGMA z-score weights (capped at 10), single-cell variance-stabilization weights, and using both sets of weights (default). We considered 3 MAGMA gene window sizes: 0 kb, 10 kb (default), and 50 kb. We applied each version of `scDRS` to subsampled TMS FACS data sets (20 repetitions with 10K cells each) and a curated set of 20 traits with expected and unexpected disease-critical cell types (Supp. Table 17). For a given `scDRS` version and a given trait (with an expected and an unexpected control cell type), we computed the t-statistic between cells from the expected and unexpected cell types, and then divided it by the average t-statistics of results of the given trait from all data sets and all versions of `scDRS` to correct for trait-specific baseline. We evaluated each version by first computing the mean and SE of the normalized t-statistics for a given trait across the 20 data sets and then combining the estimates across the 20 traits via random-effect meta-analysis. We compared the performance of a pair of versions of `scDRS` by applying the same procedure to the difference of the normalized t-statistics between the two versions. **(A)** Varying gene selection methods while fixing other parameters as the default. **(B)** Varying gene weighting methods while fixing other parameters as the default. **(C)** Varying MAGMA gene window size while fixing other parameters as the default. The default version was denoted in dark blue and 95% confidence intervals were provided, using methods as described above. Significant differences relative to the default configuration were marked by asterisks: * denotes $P < 0.05$ and ** denotes $P < 0.001$, using methods as described above. Numerical results are reported in Supp. Table 18.

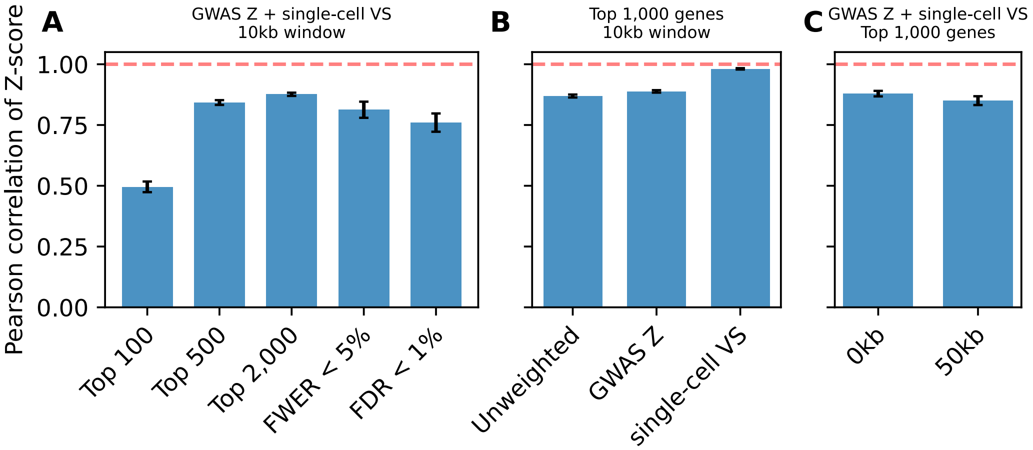


**Supplementary Figure 13. Consistency of `scDRS` results across different versions.** Average Pearson's correlation of `scDRS` disease score across the TMS FACS cells between the default version (top 1,000 genes, GWAS Z + single-cell VS weights, 10-kb window) and other versions. 95% confidence intervals were provided and were computed across the 74 diseases/traits . **(A)** Correlations with other versions of `scDRS` with different gene selection methods while keeping other parameters as the default. **(B)** Correlations with other versions of `scDRS` with different gene weighting methods while keeping other parameters as the default. **(C)** Correlations with other versions of `scDRS` with different MAGMA gene window sizes while keeping other parameters as the default.

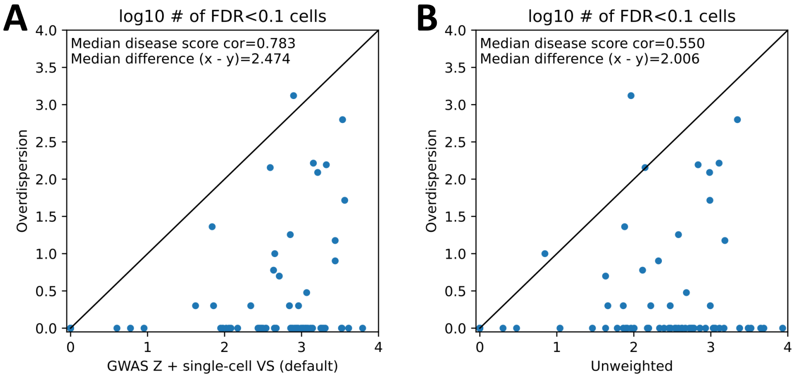


**Supplementary Figure 14. Comparison to overdispersion score.** We compared the default version (weighted average using both GWAS and single-cell weights) and the unweighted version to the overdispersion score, which tests for both overexpression and underexpression of the disease genes in the relevant cell population (Methods). Each dot corresponds to one of the 74 diseases, the x-axis denotes the number of associated cells (FDR<0.1) using the default version or the unweighted version and the y-axis denotes results using the overdispersion score. For each of these comparisons, we also computed the Pearson's correlation of the `scDRS` disease score across cells for each trait, and report the median correlation across traits. We also report the median improvement (x-values minus y-values) across traits. We determined that both the default version and the unweighted version attained higher power while being consistent with the overdispersion score.

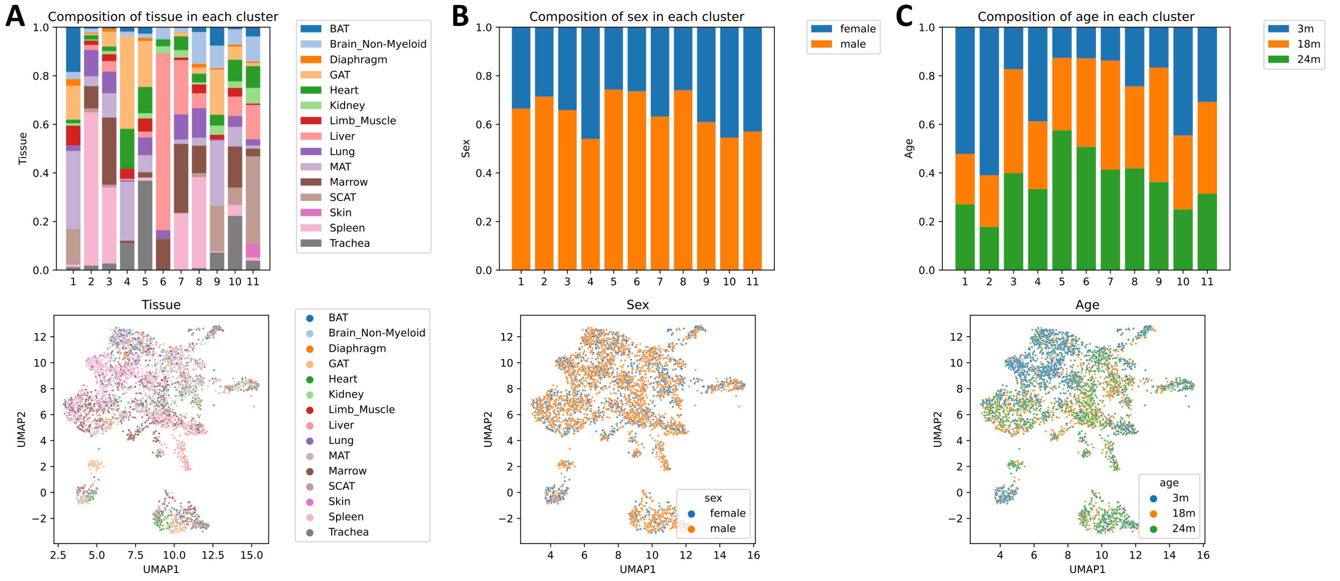


**Supplementary Figure 15.** **Covariate compositions for clusters in Fig. 4A and the corresponding UMAP visualization.** (**A**) Tissue. (**B**) Sex. (**C**) Age.

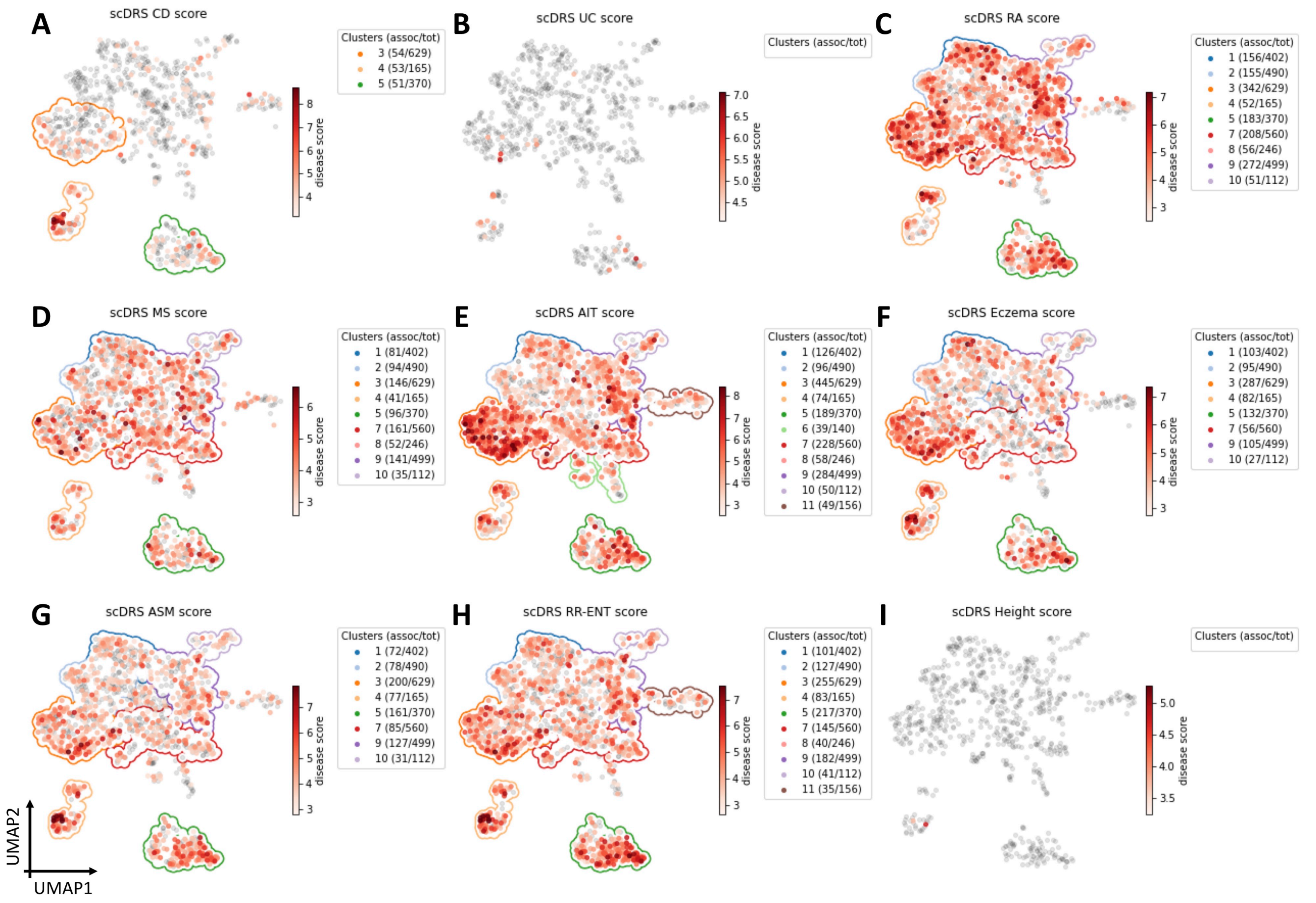


**Supplementary Figure 16.** **Subpopulations of TMS FACS T cells associated with the other 8 autoimmune diseases (besides IBD and HT reported in Fig. 4B,C) and height, a negative control trait.** Significantly associated cells (FDR<0.1) are denoted in red, with shades of red denoting `scDRS` disease scores; non-significant cells are denoted in grey. The number of associated cells and total number of cells are provided in parentheses for clusters with more than 25 associated cells.

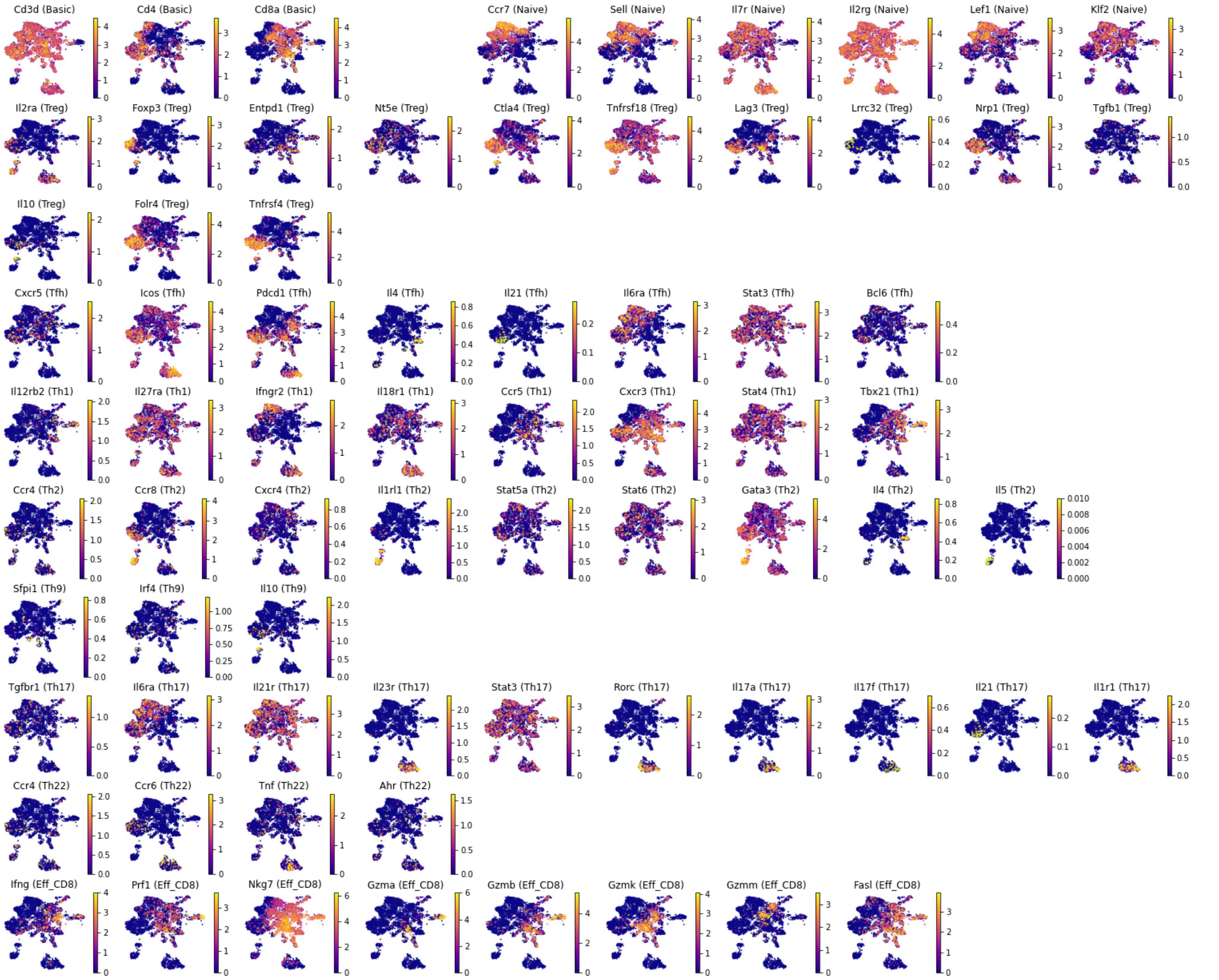


**Supplementary Figure 17. UMAP visualization of T cell marker gene expression in TMS FACS T cells.** Expression of marker genes for general T cells (*CD3*, *CD4*, *CD8A*), naive T cells, regulatory T cells (Treg), follicular helper T cells (Tfh), T helper 1 cells (Th1), T helper 2 cells (Th2), T helper 9 cells (Th9), T helper 17 cells (Th17), T helper 22 cells (Th22), and effector $CD8^+$ T cells (Eff_CD8). 3 subtypes (Tfh, Th9, Th22) were excluded from the subsequent subtype annotation. Tfh was excluded because the corresponding marker genes were not consistently expressed across cell populations and the 2 genes with highly localized expression, *ICOS* and *PDCD1*, can also be expressed in other effector T cell subtypes[78,79]. Th9 was excluded because of low expression of marker genes and the only gene with highly localized expression, *IL10*, can also be expressed in Tregs[80]. Th22 was excluded because the only gene with highly localized expression, *TNF*, can also be expressed in Th17 cells[81].

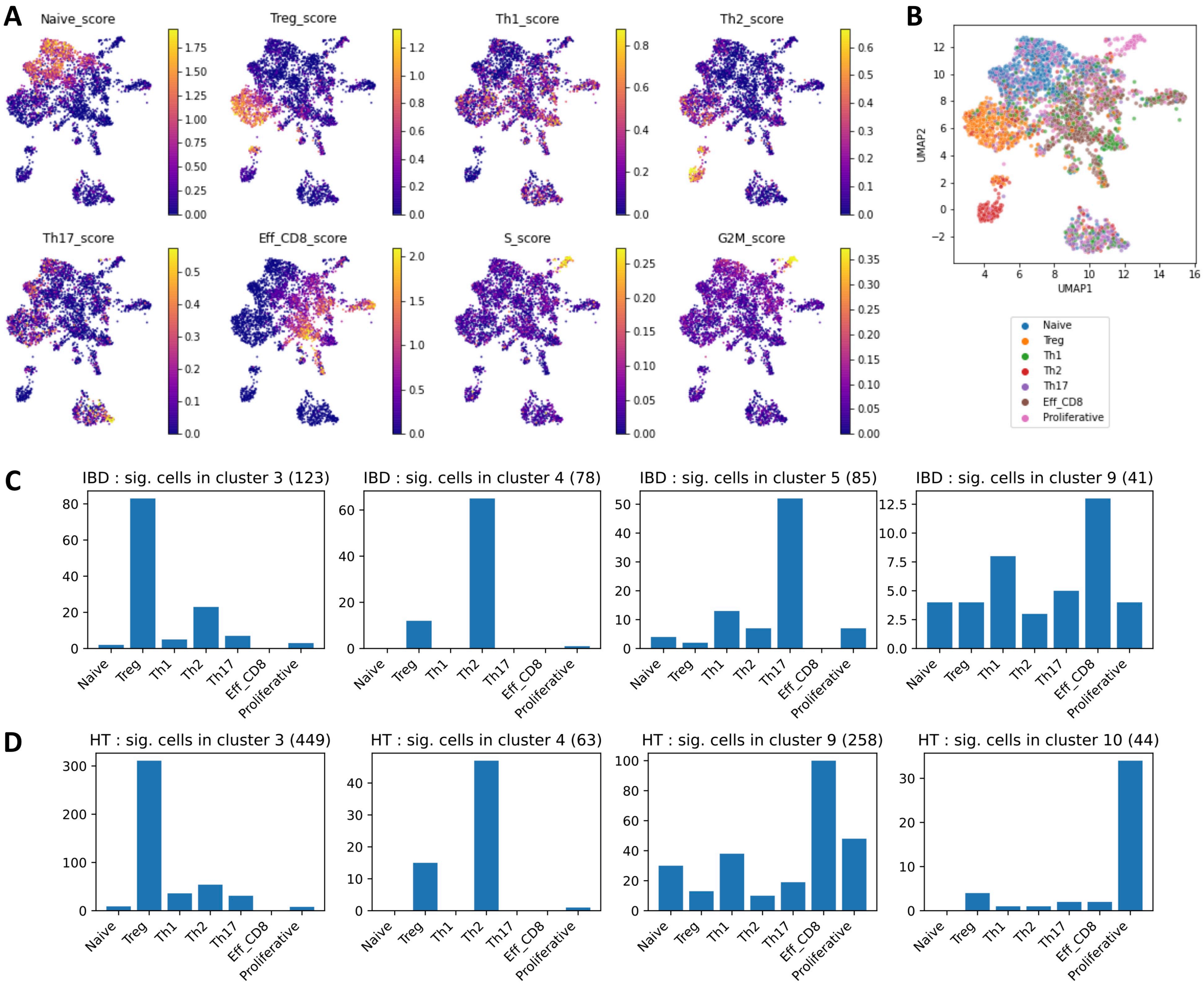


**Supplementary Figure 18.** **T cell subtype classification based on marker genes.** **(A)** Subtype scores for individual T cells, computed as the average expression of marker genes using the "scanpy.tl.score_genes" function in Scanpy[74]. Curated subtype markers (Supp. Fig. 17) were used for the subtype scores and cell-cycle markers[82] (Supp. Table 10) were used for the 2 cell-cycle scores (S_score and G2M_score). **(B)** Subtype classification based on subtype scores in panel A. The scores were first normalized (centered by median and scaled by SD across cells) and each cell was assigned a subtype label based on the highest normalized score. The 2 cell-cycle scores were merged as one term "Proliferative". **(C-D)** Subtype compositions for associated cells for IBD and HT.

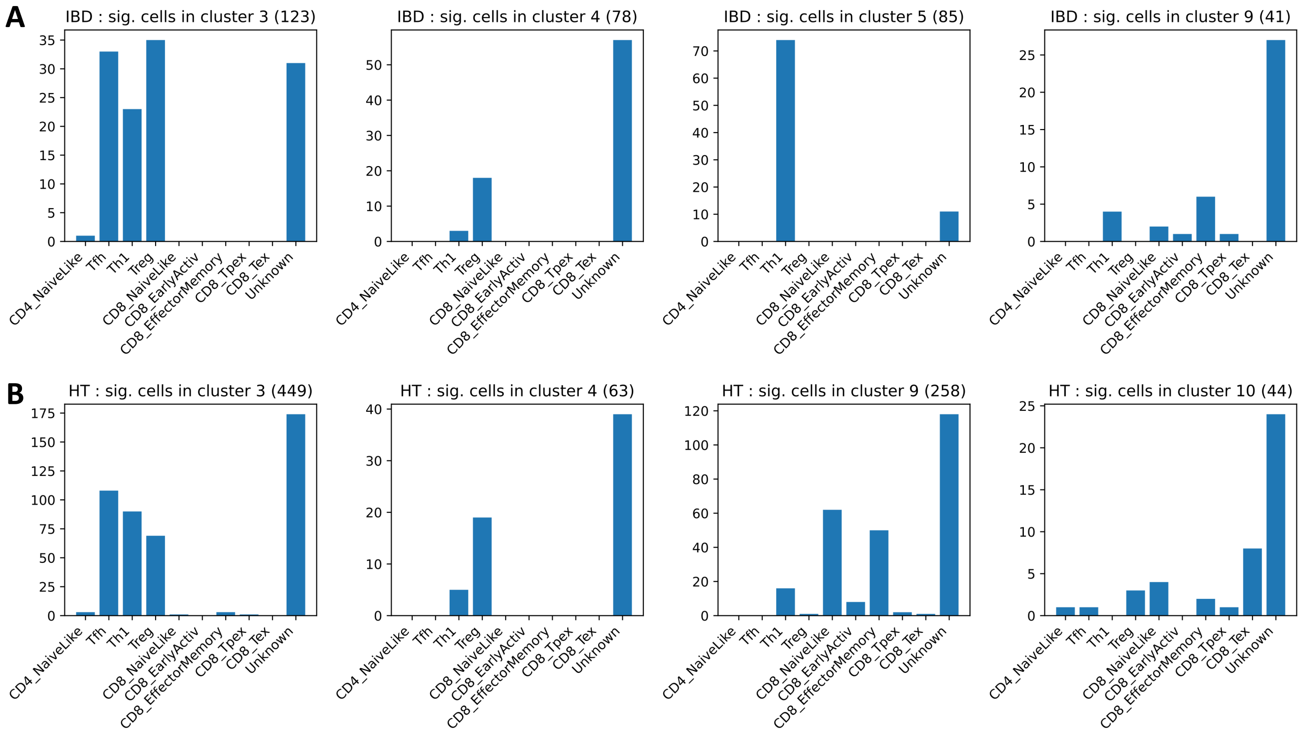


**Supplementary Figure 19. T cell subtype classification based on projecTILE[83].** The tumor-infiltrating T lymphocytes (TIL) atlas reference[83] was used and cells with confidence smaller than 0.8 were labeled as "Unknown".

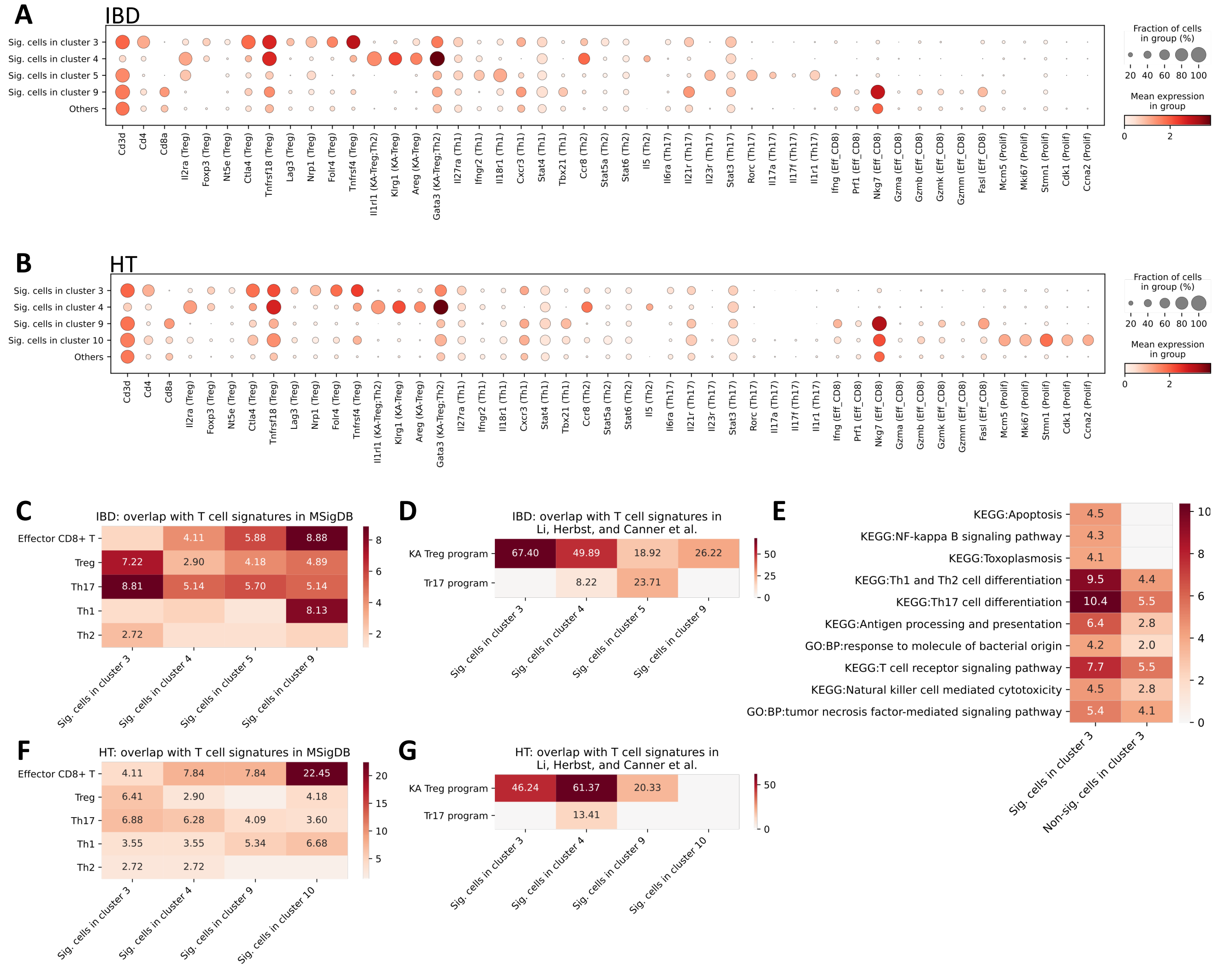


**Supplementary Figure 20. Additional results on subpopulations of T cells associated with IBD and HT. (A-B)** Dotplots for marker gene expression in disease-associated T cells in each of the 4 top clusters for IBD and HT. T cell subtypes of the marker genes were provided in parentheses. KA-Treg refers to a *KLRG1*$^+$ *AREG*$^+$ effector-like Treg program[84]. **(C)** Overlap between T cell signatures and the top 300 specifically expressed genes for each IBD-associated T cell subpopulation (Supp. Table 10; Methods). The color and number in each cell represent the $-\log_{10}$ enrichment p-values (Fisher's exact test). **(D)** Overlap between Treg programs[84] (KA Treg refers to a *KLRG1*$^+$ *AREG*$^+$ effector-like Treg program; Tr17 refers to a Th17-like Treg program; Supp. Table 10) and the top 300 specifically expressed genes of each IBD-associated T cell subpopulation. The color and number in each cell represent the $-\log_{10}$ enrichment p-values (Fisher's exact test). **(E)** Pathways enriched in the top 300 specifically expressed genes of the IBD-associated cells and non-associated cells in cluster 3 respectively. The color and number in each cell represent the $-\log_{10}$ enrichment p-values (based on Enrichr implemented in GSEAPY[85,86]). **(F-G)** Results for HT similar to panels C and D.

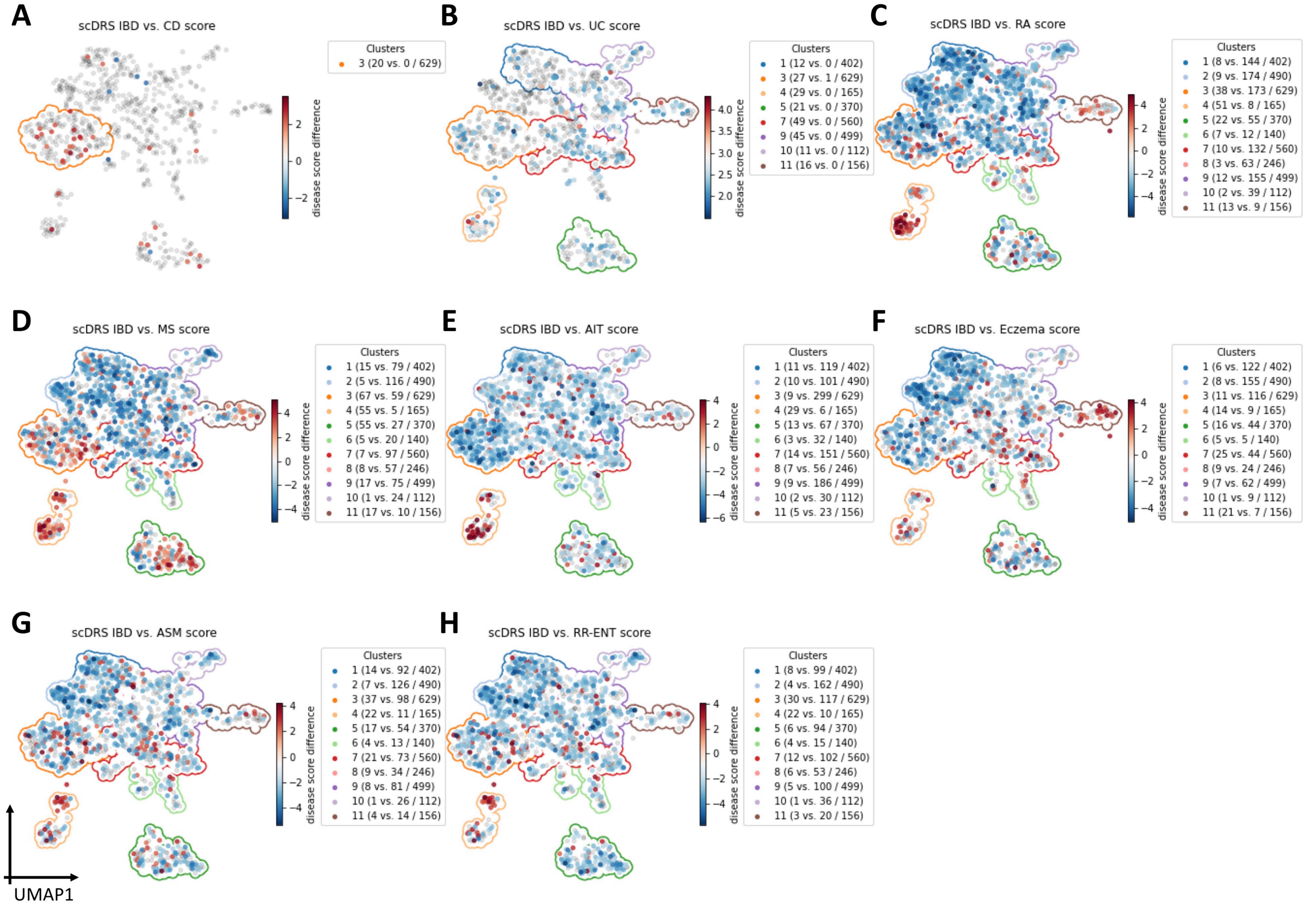


**Supplementary Figure 21. Differences in individual cell-level associations between IBD and the other 8 autoimmune diseases (besides HT reported in Fig. 4D).** Differentially associated cells (absolute score difference>2) are denoted in red and blue, with shades of colors denoting `scDRS` disease score differences; other cells are denoted in grey. The number of IBD-enriched cells, enriched cells for the other disease, and all cells in the cluster are provided in parentheses for clusters with more than 25 differentially associated cells. Associations of T cells to IBD are moderately different from associations of T cells to other autoimmune diseases.

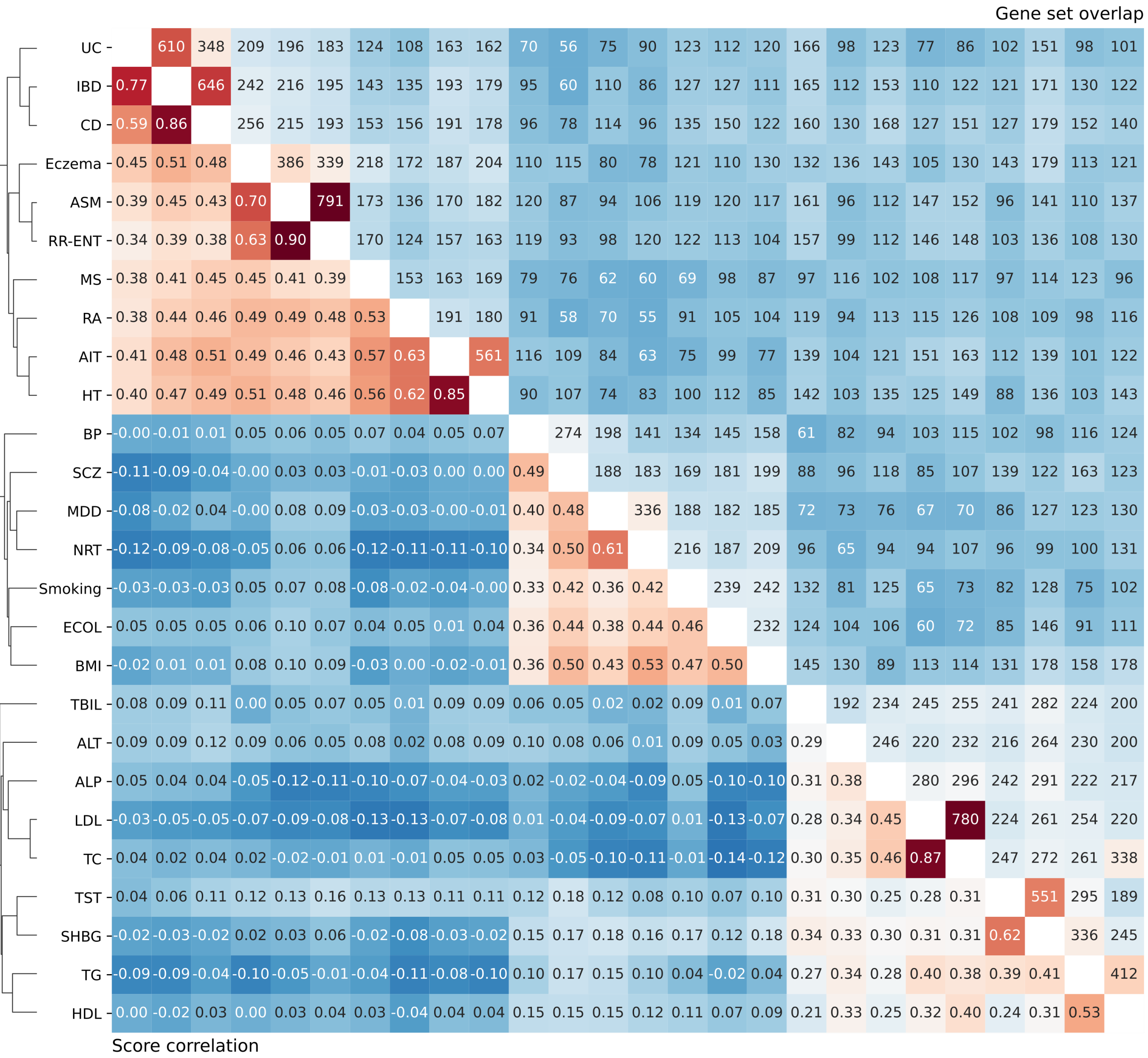


**Supplementary Figure 22. Numbers of overlapping genes (upper triangle) and correlations of the `scDRS` disease scores across all TMS FACS cells (lower triangle) between the 26 autoimmune, brain, and metabolic traits analyzed in the main paper.** Traits are ordered via hierarchical clustering of the `scDRS` score correlation and the clustering dendrogram was provided. The level of gene set overlap is moderate. `scDRS` disease score correlations distinguish diseases/traits from the 3 categories as well as subgroups of diseases/traits in the same category.

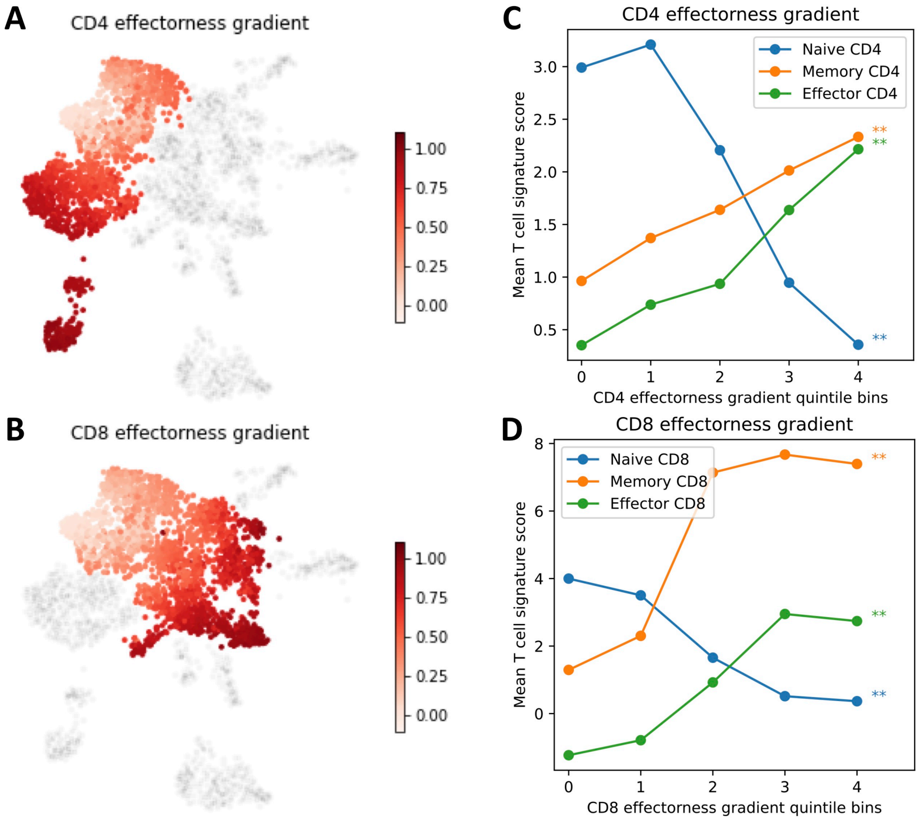


**Supplementary Figure 23. Results for CD4 and CD8 effectorness gradients.** (**A**) UMAP visualization of CD4 effectorness gradient across CD4$^+$ T cells. The effectorness gradient was represented in the red color and non-CD4$^+$ T cells were colored in grey. (**B**) UMAP visualization of CD8 effectorness gradient across CD8$^+$ T cells. The effectorness gradient was represented in the red color and non-CD8$^+$ T cells were colored in grey. (**C**) Correlation across CD4$^+$ T cells between CD4 effectorness gradient and signatures for naive, memory, and effector CD4$^+$ T cells (`scDRS` disease scores for applying these signature gene sets to the TMS FACS data). The x-axis denotes CD4 effectorness gradient quintile bins and the y-axis denotes average `scDRS` disease score for each bin and each signature gene set. * denotes $P < 0.05$ and ** denotes $P < 0.005$. (**D**) Correlation across CD8$^+$ T cells between CD8 effectorness gradient and signatures for naive, memory, and effector CD8$^+$ T cells. The x-axis denotes CD8 effectorness gradient quintile bins and the y-axis denotes average `scDRS` disease score for each bin and each signature gene set. * denotes $P < 0.05$ and ** denotes $P < 0.005$.

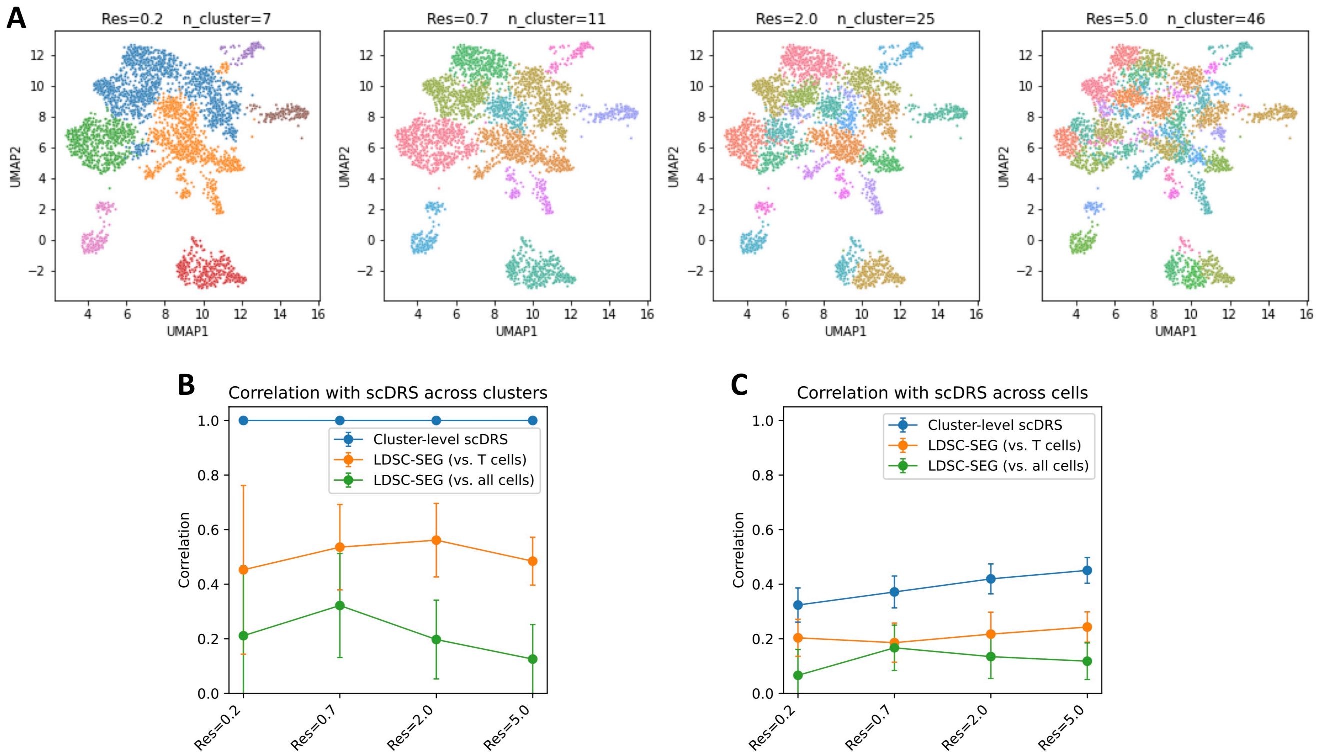


**Supplementary Figure 24. Comparison of individual cell level T cell results to cluster-level analyses.** We reclustered the set of T cells with different cluster resolutions (0.2, 0.7, 2, 5; 0.7 corresponds to Fig. 4A), followed by performing LDSC-SEG analysis on specifically expressed genes (SEGs) for each cluster and each of the 10 autoimmune diseases; we considered two ways for computing SEGs for a given cluster: by comparing to other T cells (vs. T cells) or to all other cells in the TMS FACS data (vs. all cells). We considered a third baseline that obtains $-\log_{10}$ p-values for a given cluster by averaging the `scDRS` log10 p-values of cells within the cluster (cluster-level `scDRS`). We note that the gap between `scDRS` and cluster-level `scDRS` is due to finite clustering resolution, while the gap between cluster-level `scDRS` and LDSC-SEG is due to the difference between `scDRS` and LDSC-SEG for capturing cluster-level disease heritability enrichment. **(A)** T cells clustered at different resolutions. **(B)** Correlation of $-\log_{10}$ p-value *across clusters* between `scDRS` and the 3 comparison methods. For `scDRS`, the cluster-level $-\log_{10}$ p-values were obtained by averaging the $-\log_{10}$ p-value of cells within the same cluster (identical to cluster-level `scDRS`). The three methods were highly correlated with `scDRS` at cluster level, suggesting LDSC-SEG and `scDRS` produced similar results. **(C)** Correlation of $-\log_{10}$ p-value *across cells* between `scDRS` and the 3 comparison methods. For the 3 comparison methods, cell-level $-\log_{10}$ p-values were obtained by assigning the same cluster-level $-\log_{10}$ p-value to all cells within the cluster. The 3 methods were less correlated with `scDRS` at individual cell level, suggesting cluster-level analyses were not able to capture the individual cell-disease associations detected in the `scDRS` individual cell-level analysis.

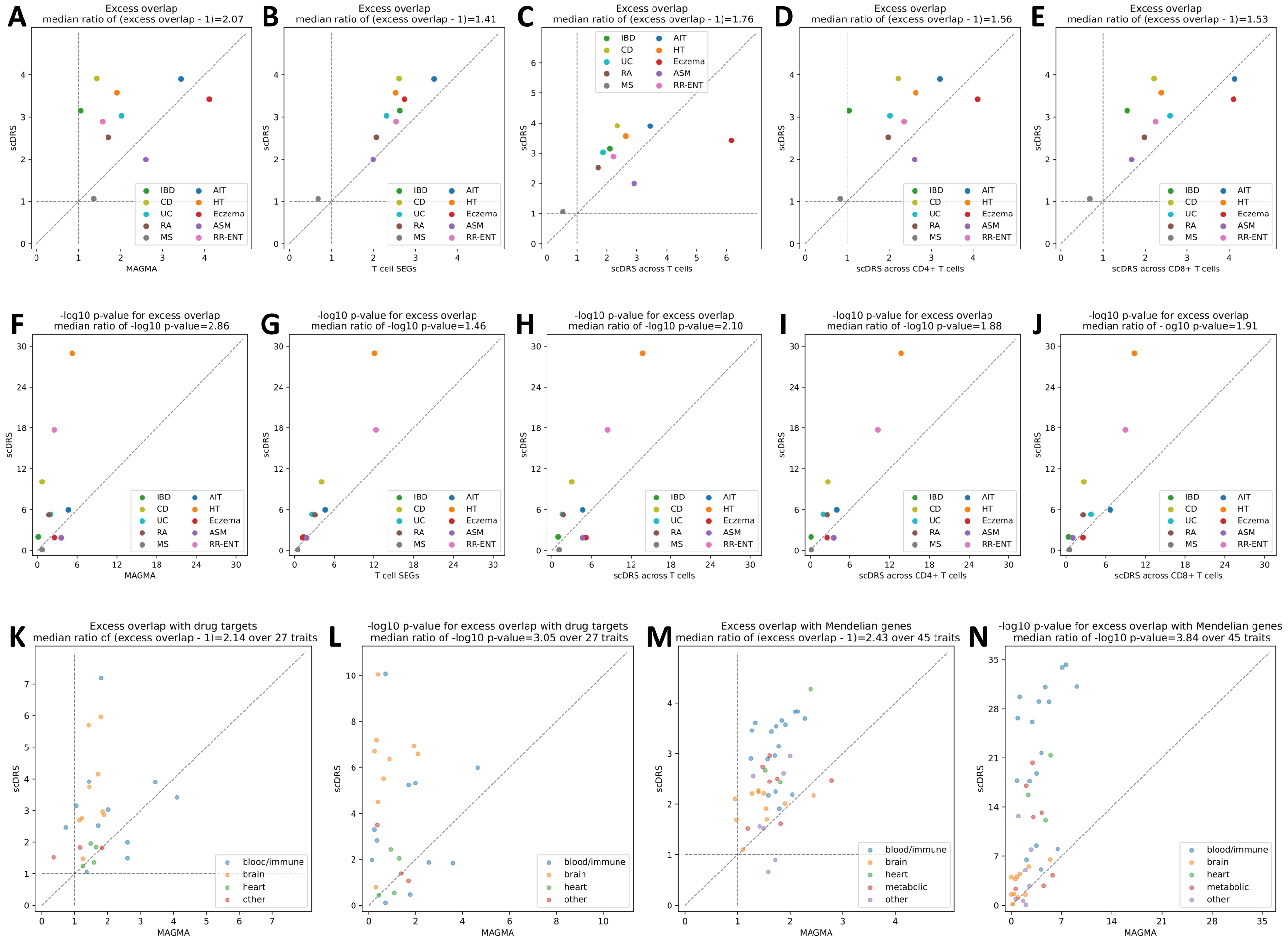


**Supplementary Figure 25. Additional results on disease gene prioritization.** **(A-J)** Comparison to alternative disease gene prioritization methods for the 10 autoimmune diseases. The first row shows levels of excess overlap between the prioritized disease genes and the gold standard gene sets while the second row shows the corresponding $-\log_{10}$ p-values for excess overlap. Each dot corresponds to a disease, the y-axis shows results for the proposed prioritization method (correlating gene expression levels with the `scDRS` disease score across all TMS FACS cells), and the x-axis shows results from comparison methods, including (from left to right) top 1,000 MAGMA genes, top 1,000 genes specifically expressed in T cells (vs. the rest of cells in TMS FACS), prioritization based on correlation across T cells (instead of all TMS FACS cells), prioritization based on correlation across CD4$^+$ T cells (instead of all TMS FACS cells), and prioritization based on correlation across CD8$^+$ T cells (instead of all TMS FACS cells). **(K-L)** Overlap with drug target genes for 27 diseases. **(M-N)** Overlap with Mendelian disease genes for 45 diseases. The median ratio of $-\log_{10}$ p-values and (excess overlap $-$ 1) between the y- and x-values (median of ratios) was provided in the figure title.

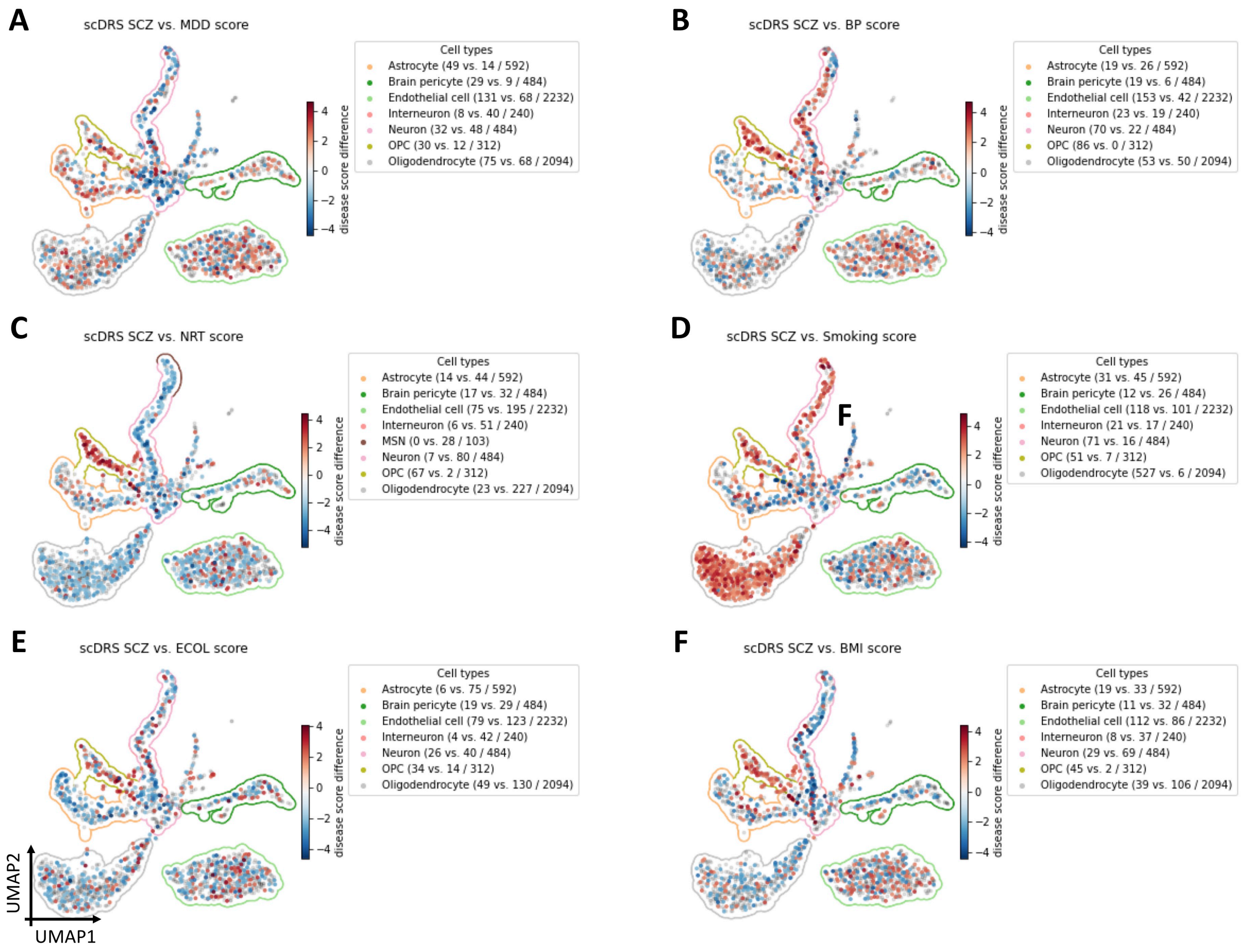


**Supplementary Figure 26. Differences in individual cell-level associations between SCZ and the other 6 brain diseases/traits across the TMS FACS brain non-myeloid cells.** Differentially associated cells (absolute score difference>2) are denoted in red and blue, with shades of colors denoting `scDRS` disease score differences; other cells are denoted in grey. The number of SCZ-enriched cells, enriched cells for the other disease/trait, and all cells in the cluster are provided in parentheses for TMS cell types with more than 25 differentially associated cells. Associations of brain cells to SCZ are moderately different from associations of brain cells to other brain diseases/traits. Particularly, oligodendrocytes are more strongly associated with SCZ than Smoking.

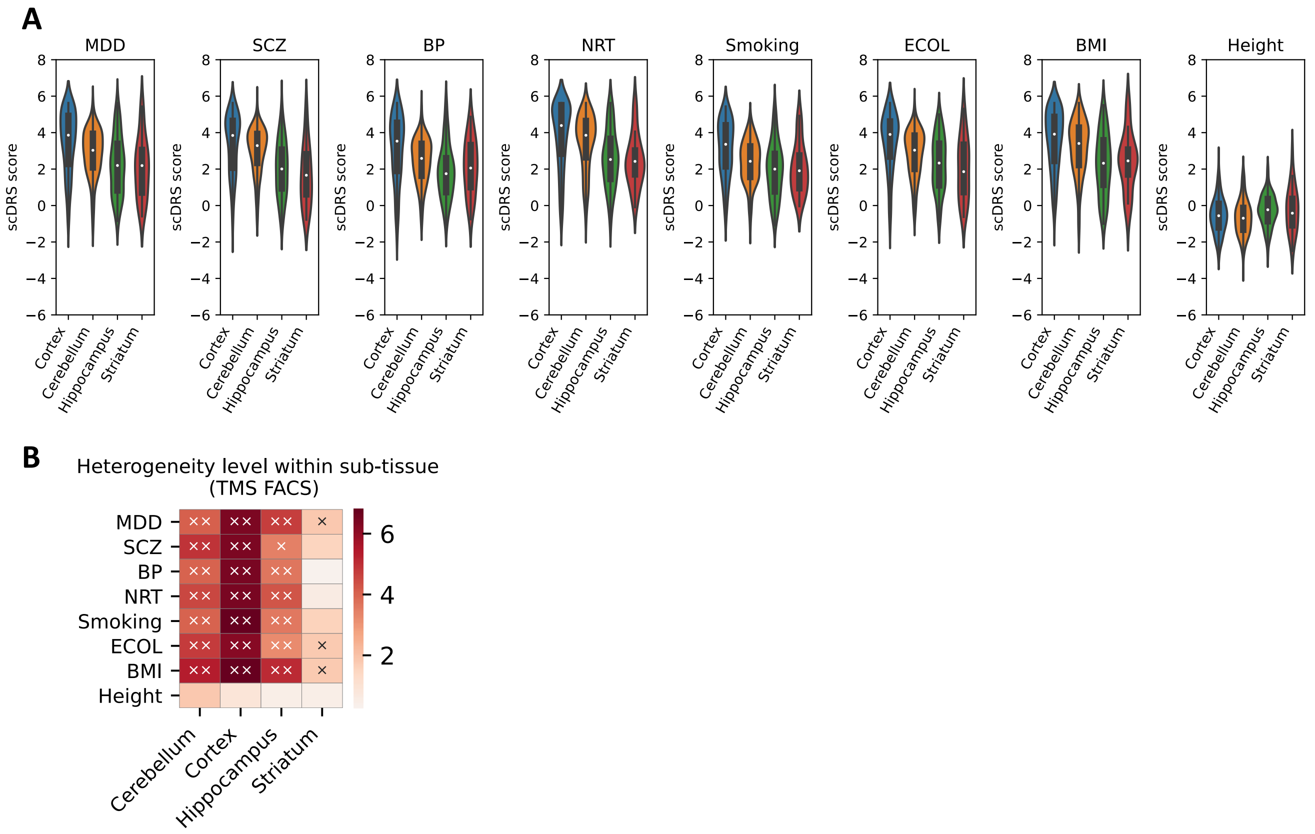


**Supplementary Figure 27. Associations of TMS FACS neurons with brain-related diseases/traits. (A)** Violin plots of the `scDRS` disease score for TMS FACS brain neurons (undetermined neurons, excluding interneurons and MSNs) in different brain subtissues and for different diseases/traits. Height was included as a negative control. **(B)** Within-subtissue heterogeneity of neurons in association with different diseases/traits. Heatmap colors represent the heterogeneity z-score (MC z-score) and cross symbols represent significant within-subtissue disease association heterogeneity (× denotes $P$ <0.05 and ×× denotes $P$ <0.005, MC test).

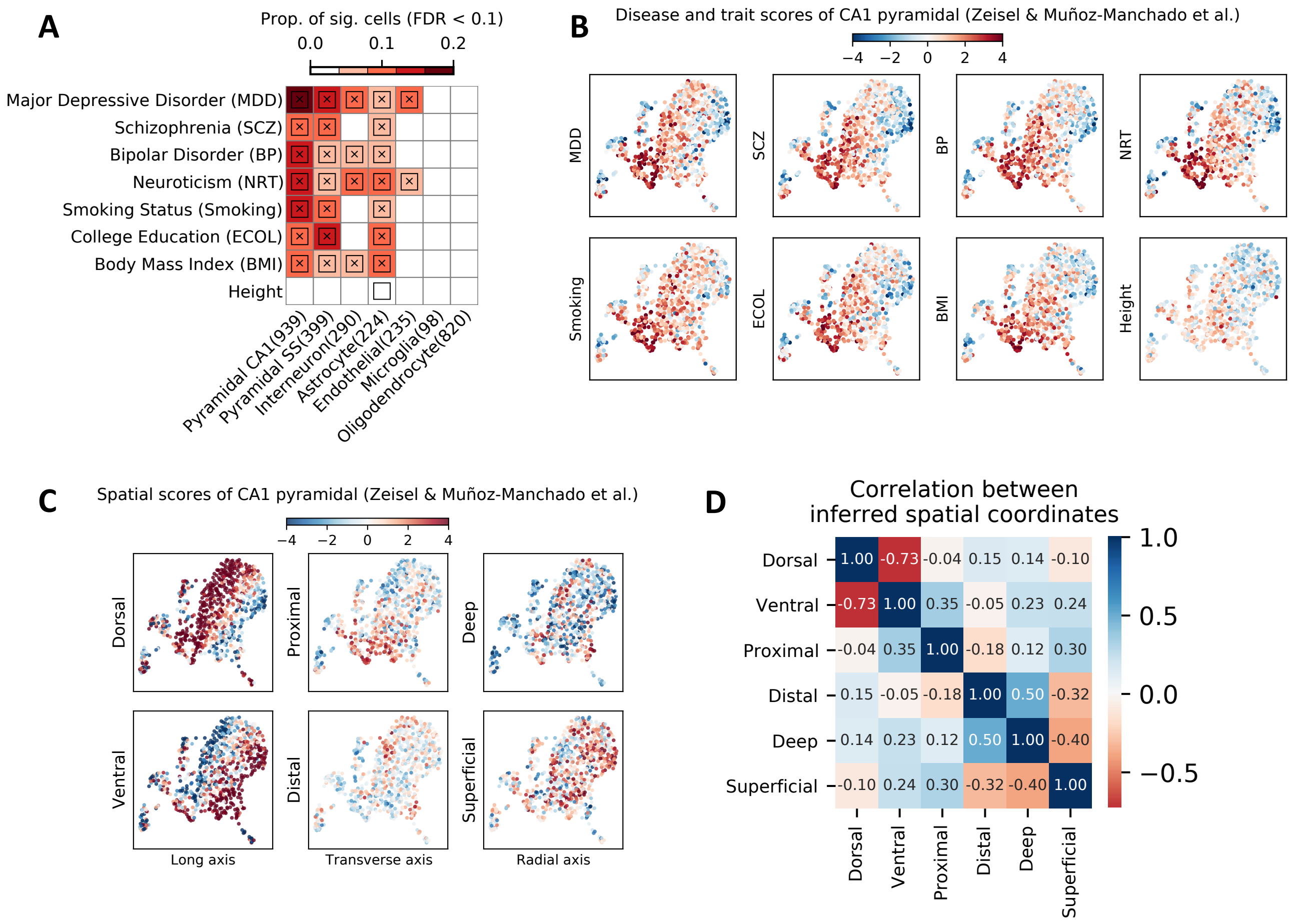


**Supplementary Figure 28. Additional results for the Zeisel & Muñoz-Manchado et al. data on associations to brain-related diseases/traits. (A)** Disease associations at the cell type level. Each row represents a disease/trait and each column represents a cell type. Heatmap colors for each cell type-disease pair denote the proportion of significantly associated cells (FDR<0.1 across all cells for a given disease). Squares denote significant cell type-disease associations (FDR<0.05 across all pairs of 7 cell types and 8 diseases/traits). Cross symbols denote significant heterogeneity in association with disease across individual cells within a given cell type (FDR<0.05 across all pairs). Heatmap colors and cross symbols are omitted for cell type-disease pairs with non-significant cell type-disease associations; the plotting style is the same as in Fig. 3. **(B)** UMAP visualization of the CA1 pyramidal neurons in the Zeisel & Muñoz-Manchado et al. data with color representing the `scDRS` disease score (extending results in Fig. 5A). Height was included as a negative control trait. **(C)** UMAP visualization of the CA1 pyramidal neurons with color representing the inferred spatial coordinates. **(D)** Pairwise correlations across cells for the 6 inferred spatial coordinates in the Zeisel & Muñoz-Manchado et al. data set.

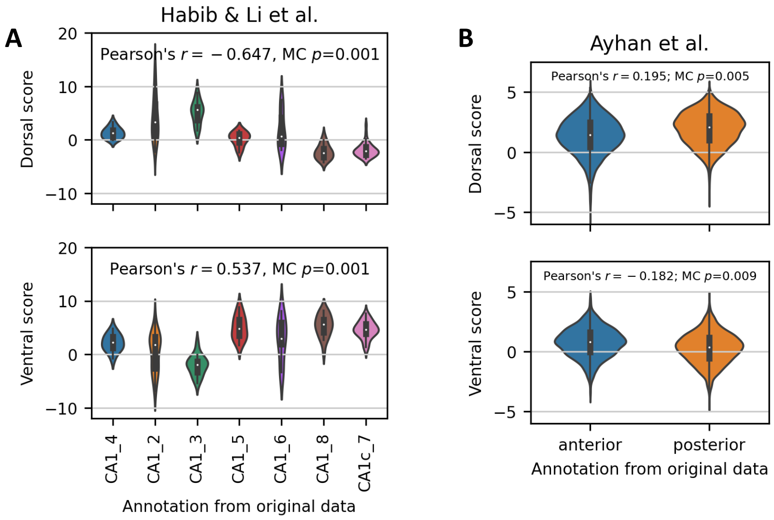


**Supplementary Figure 29. Validation of inferred spatial coordinates of CA1 pyramidal neurons using data sets with annotated spatial annotations.** Violin plots of the inferred cell-level spatial coordinates (y-axis) against the provided spatial annotations (x-axis). **(A)** Habib & Li et al. data. The x-axis labels were ordered from dorsal to ventral neurons according to the provided spatial annotation (Fig. 2B in Habib & Li et al.[62]). Both the dorsal and ventral scores are significantly associated with the provided spatial annotation ($P$<0.001, MC test based on Pearson's correlation between the provided spatial annotation (ordinal ranks of the 7 categories) and the inferred spatial score). **(B)** Ayhan et al. data. Anterior corresponds to the ventral region while posterior corresponds to the dorsal region. Both the dorsal and ventral scores are significantly associated with the spatial annotation ($P$<0.01, MC test based on Pearson's correlation between provided spatial annotation (ordinal ranks of the 2 categories) and the inferred spatial score). Annotated spatial coordinates on the transverse and radial axes were not available in these data sets.

A

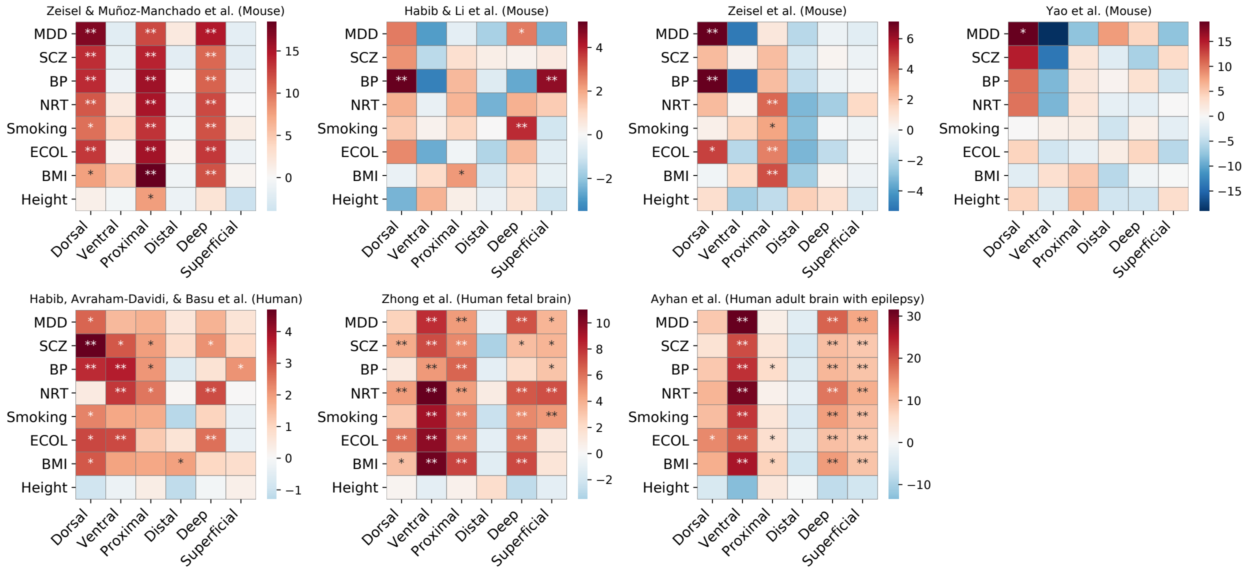


B

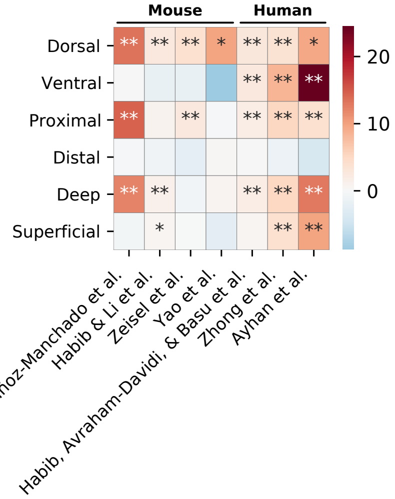


C

| Location | Mouse | Human |
|---|---|---|
| Dorsal | ✓ | ✓ |
| Ventral | | ✓ |
| Proximal | ✓ | ✓ |
| Distal | | |
| Deep | ✓ | ✓ |
| Superficial | | ✓ |

**Supplementary Figure 30. Complete results of correlations between scDRS disease scores and inferred spatial coordinates across CA1 pyramidal neurons in 7 single-cell data sets (extending results in Fig. 5B). (A)** Results for regressing the scDRS disease scores against the inferred spatial coordinates for each disease/trait and each inferred spatial coordinate. Color represents the $t$-statistics and stars represent significant associations (* denotes $P < 0.05$ and ** indicates $P < 0.005$, MC test; Methods). For clarification, Zeisel & Muñoz-Manchado et al. refers to the data from Zeisel & Muñoz-Manchado et al. 2015 *Science*[60] and Zeisel et al. refers to the data from Zeisel et al. 2018 *Cell*[61]. **(B)** Summary of results in panel A. Heatmap color represent the average $t$-statistics across the 7 brain-related diseases/traits (excluding height) for each data set and stars represent significant associations by combining p-values across datasets using Fisher's combined probability test. **(C)** Summary of the association between brain-related diseases and the inferred spatial coordinates for the mouse and human data sets in panel B.

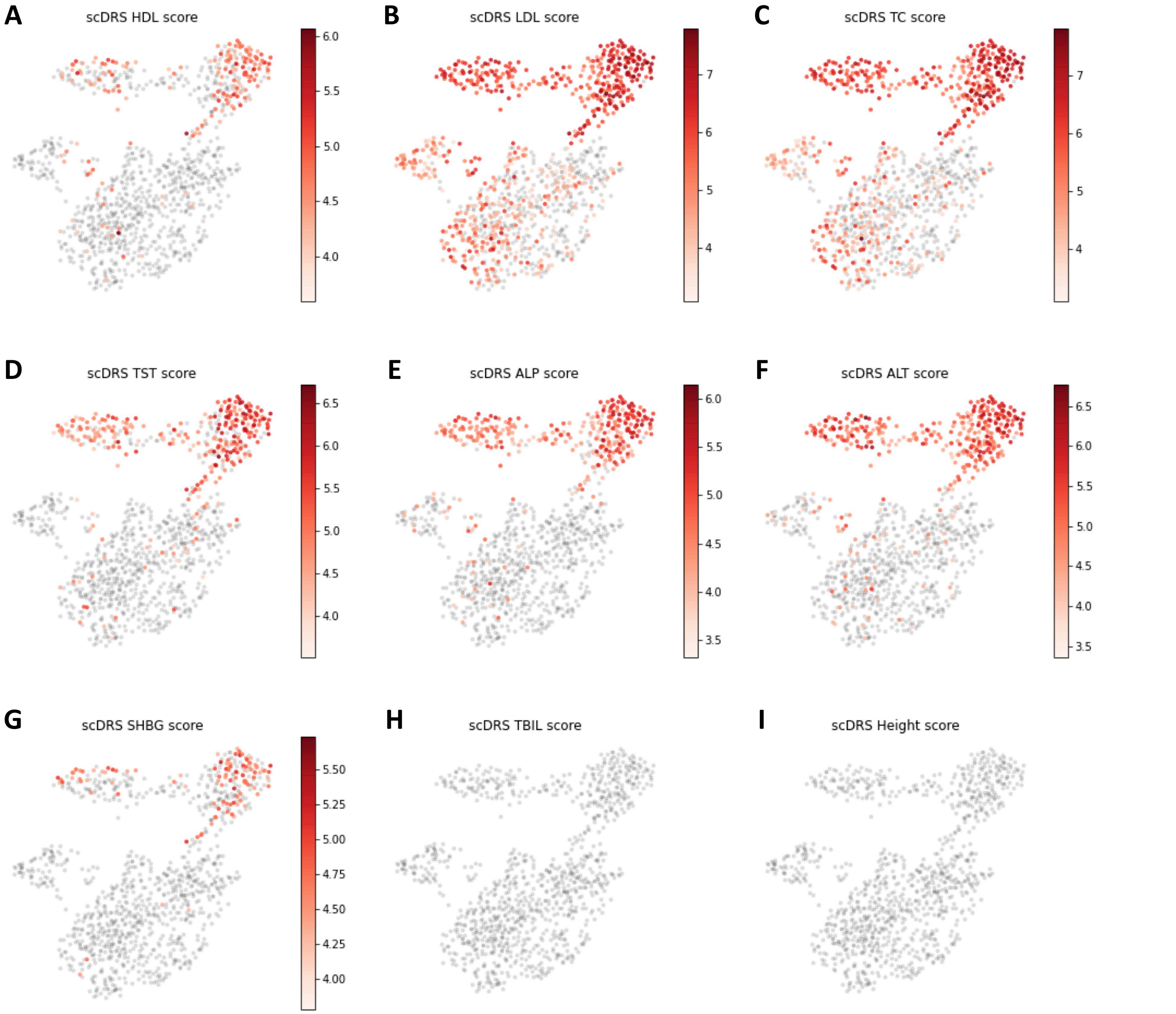


**Supplementary Figure 31. Subpopulations of TMS FACS hepatocytes associated with the other 8 metabolic diseases (besides TG reported in Fig. 5C) and height.** Significantly associated cells (FDR<0.1) were colored in red with shades of red representing the `scDRS` disease score; non-significant cells were colored in grey. The color bar was removed for traits without significant associations.

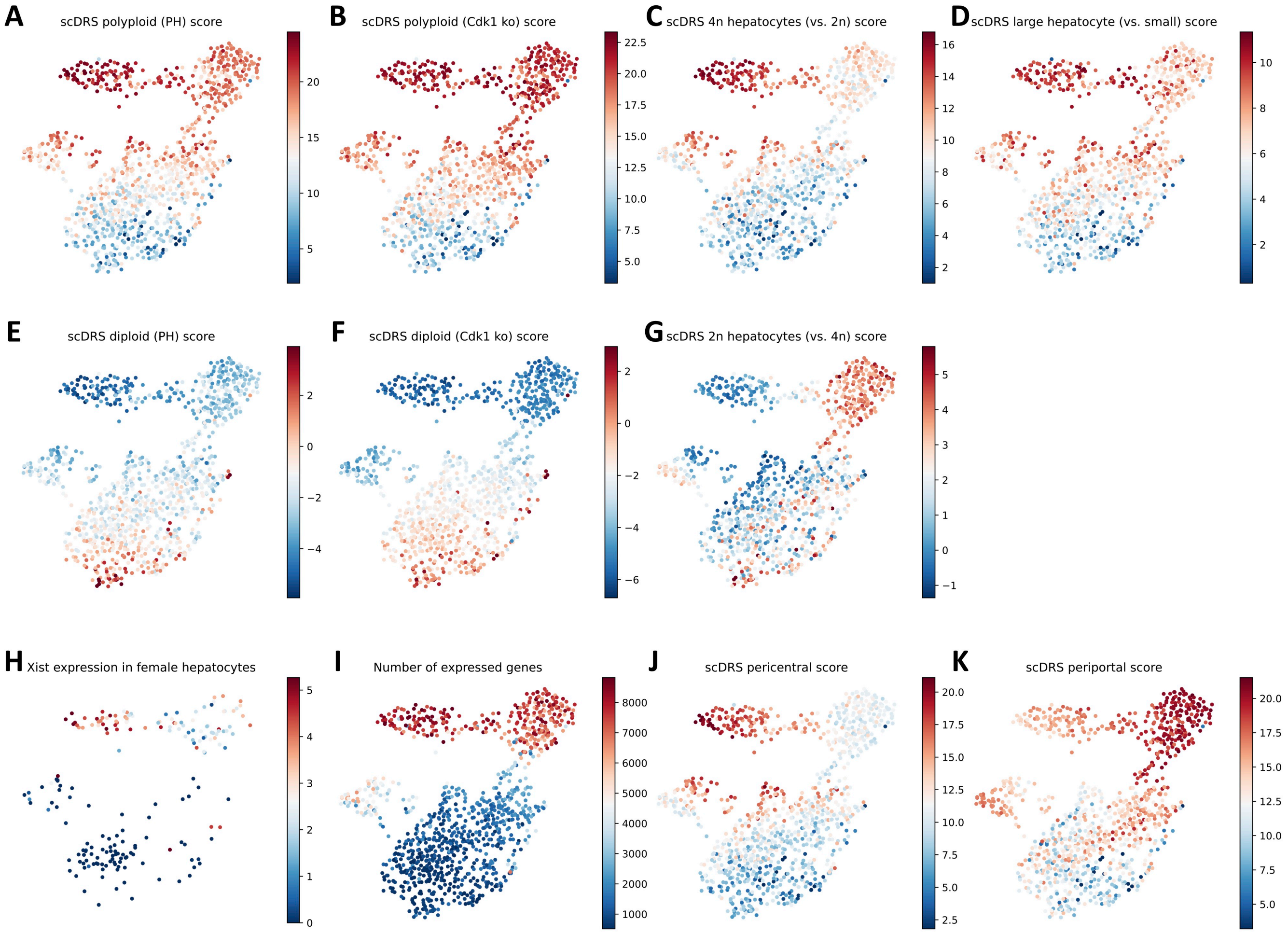


**Supplementary Figure 32.** **Signature scores for hepatocyte ploidy level and zonation. (A-D)** `scDRS` score for polyploid hepatocyte signatures: partial hepatectomy (PH) vs. pre-PH, Cdk1 knockout vs. control, 4n vs. 2n hepatocytes, large vs. small hepatocytes. **(E-G)** `scDRS` score for diploid hepatocyte signatures: pre-PH vs. PH, control vs. Cdk1 knockout, 2n vs. 4n hepatocytes. **(H)** Expression of *Xist* in female hepatocytes (expected to have high expression in high-ploidy female hepatocytes). **(I)** Number of genes (expected to be high in high-ploidy hepatocytes). **(J)** `scDRS` score for pericentral hepatocyte signatures. **(K)** `scDRS` score for periportal hepatocyte signatures.

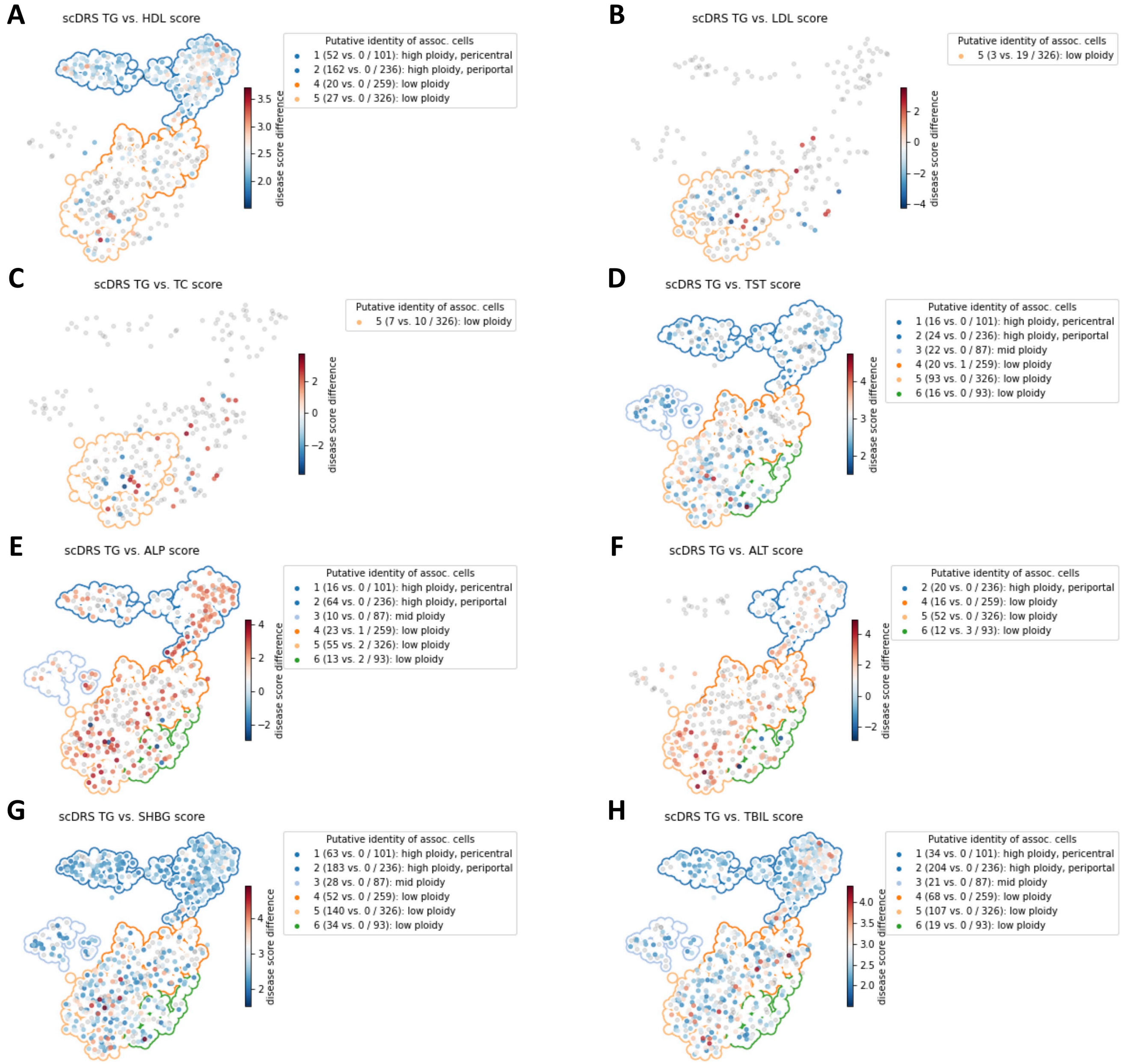


**Supplementary Figure 33. Differences in individual cell-level associations between TG and the other 8 metabolic traits across the TMS FACS hepatocytes.** Differentially associated cells (absolute score difference>2) are denoted in red and blue, with shades of colors denoting `scDRS` disease score differences; other cells are denoted in grey. The number of TG-enriched cells, enriched cells for the other trait, and all cells in the cluster are provided in parentheses for TMS cell types with more than 25 differentially associated cells. Associations of hepatocytes to TG are similar to associations of hepatocytes to other metabolic traits.

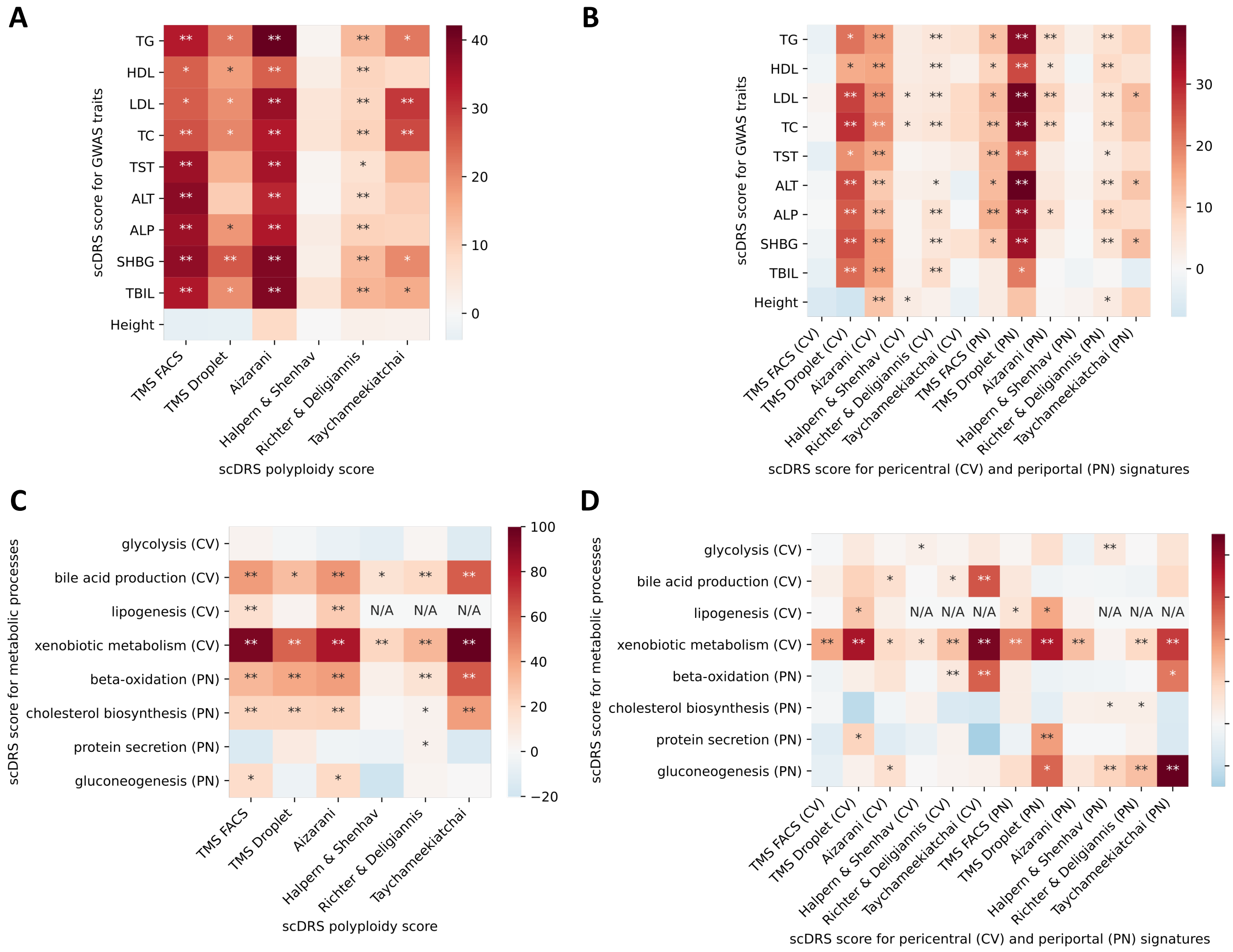


**Supplementary Figure 34.** **Complete results of joint regression analysis for GWAS metabolic traits and putative zonated metablic processes across the 6 data sets (extending results in Fig. 5D).** **(A-B)** Results for the 9 metabolic traits and height, a negative control triat. The polyploidy score (panel A) and both the pericentral and periportal score (panel B) were consistently associated with the 9 metabolic traits across the data sets. The strong association ($P < 0.005$) between the pericentral score and height in the Aizarani et al. data may be because that we inferred the pericentral score using mouse gene signatures, which are less conserved in human (as also mentioned in the original paper[20]). **(C-D)** Results for the 8 metabolic pathways. Overall, as shown in panel D, the pericentral score was associated with pericentral-specific pathways (first 4 rows) while the periportal score was associated with periportal-specific pathways (last 4 rows). * denotes $P < 0.05$ and ** denotes $P < 0.005$.

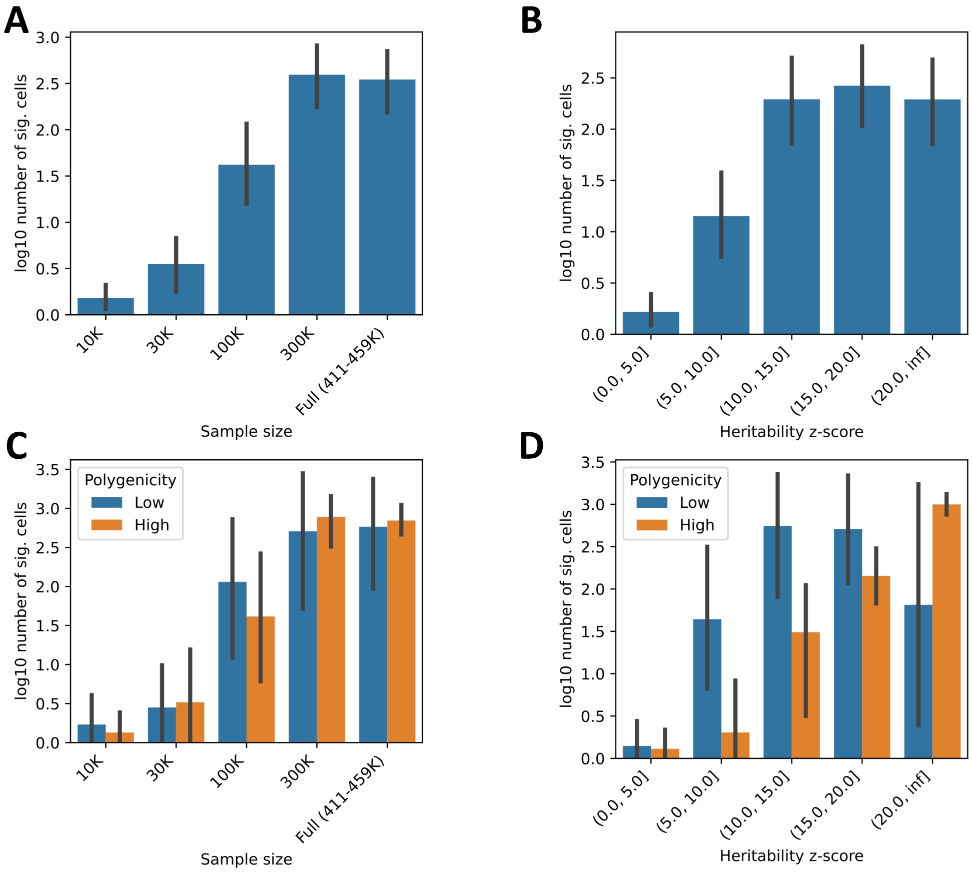


**Supplementary Figure 35. Impact of GWAS power on `scDRS` results.** We assessed the impact of GWAS power on the `scDRS` results by applying `scDRS` to the TMS FACS data and GWAS summary statistics computed at different sample sizes across 35 UK Biobank diseases/traits (subsamples of 10K, 30K, 100K, 300K samples, in addition to the full set of samples varying form 411K to 459K across traits). We roughly defined GWAS power using either sample size or z-score for non-zero SNP-heritability (heritability z-score) and evaluated the `scDRS` power based on number of significant cells (FDR<0.1). **(A-B)** log10 number of discoveries for the 35 traits grouped by GWAS sample size (panel A) or GWAS heritability z-score (panel B). **(C-D)** Results stratified by polygenicity for the 14 traits with polygenicity information; high for the top 50% high-polygenicity traits. 95% confidence intervals were provided for traits in each group. We determined that `scDRS` discoveries increase with GWAS power as defined using either metric, and that a GWAS sample size greater than 100K or heritability z-score greater than 5 is desirable for `scDRS` to produce a reasonable number of discoveries across diseases/traits (although less stringent thresholds can be used for less polygenic traits). The 35 traits are AIT, ALP, ALT, ASM, BMD-HT, BMI, BMR, Breast cancer, CVD, DBP, ECOL, EOS, EY, Eczema, HT, HTN, Hair color, HbA1c, Height, LDL, LYM, MCH, MONO, NCH, PLT, RBC, RDW, RR-ENT, SBP, Smoking, TBIL, TC, TG, WBC, WHR. The 14 traits with the polygenicity information are: AIT, ASM, BMD-HT, EOS, Eczema, RBC, RDW (low-polygenicity), and BMI, ECOL, Height, NCH, SBP, Smoking, WBC (high-polygenicity).
